# The mechanical regulation of RNA binding protein hnRNPC in the failing heart

**DOI:** 10.1101/2021.08.27.457906

**Authors:** Fabiana Martino, Nandan Mysore Varadarajan, Ana Rubina Perestrelo, Vaclav Hejret, Helena Durikova, Vladimir Horvath, Francesca Cavalieri, Frank Caruso, Waleed S. Albihlal, André P. Gerber, Mary A. O’Connell, Stepanka Vanacova, Stefania Pagliari, Giancarlo Forte

**Affiliations:** International Clinical Research Center (ICRC) of St Anne’s University Hospital, CZ-65691 Brno, Czech Republic; Faculty of Medicine, Department of Biology, Masaryk University, CZ-62500 Brno, Czech Republic; Competence Center for Mechanobiology in Regenerative Medicine, INTERREG ATCZ133, CZ-62500 Brno, Czech Republic; Central European Institute of Technology (CEITEC), Masaryk University; National Centre for Biomolecular Research, Masaryk University, Brno, Czech Republic; Centre for Cardiovascular and Transplant Surgery, Brno, Czech Republic; ARC Centre of Excellence in Convergent Bio-Nano Science and Technology, and the Department of Chemical Engineering, The University of Melbourne, Parkville, Victoria 3010, Australia; Dipartimento di Scienze e Tecnologie Chimiche, Università degli Studi di Roma Tor Vergata, via della Ricerca Scientifica 1, 00133, Rome, Italy; Dept. Microbial Sciences, Faculty of Health and Medical Sciences, University of Surrey, Guildford GU2 7XH, United Kingdom

**Keywords:** Heart failure, alternative splicing, mechanotransduction, RNA-binding proteins, hnRNPC

## Abstract

Cardiac pathologies are characterized by intense remodeling of the extracellular matrix (ECM) that eventually leads to heart failure. Cardiomyocytes respond to the ensuing biomechanical stress by re-expressing fetal contractile proteins via transcriptional and post-transcriptional processes, like alternative splicing (AS). Here, we demonstrate that the heterogeneous nuclear ribonucleoprotein C (hnRNPC) is upregulated and relocates to the sarcomeric Z-disk upon ECM pathological remodeling. We show that this is an active site of localized translation, where the ribonucleoprotein associates to the translation machinery. Alterations in hnRNPC expression and localization can be mechanically determined and affect the AS of numerous mRNAs involved in mechanotransduction and cardiovascular diseases, like Hippo pathway effector YAP1. We propose that cardiac ECM remodeling serves as a switch in RNA metabolism by impacting an associated regulatory protein of the spliceosome apparatus. These findings offer new insights on the mechanism of mRNAs homeostasis mechanoregulation in pathological conditions.

## INTRODUCTION

The onset and progression of aging-associated pathologies is paralleled by continuous local extracellular matrix (ECM) remodeling. This remodeling serves as a compensatory strategy for tissues to cope with altered conditions ^1^. ECM remodeling is perceived and transmitted within a cell through the modulation of cytoskeleton-propagated intracellular tension ^2^ that eventually affects tissue-specific cell responses and function ^3,4^. Cardiac pathologies are a prime example of how ECM remodeling can impact on cell functionality, here affecting muscle contractility and organ pumping ^5^. Independently of the etiology, cardiac remodeling confers a major shift in cardiomyocyte contractile function by inducing changes in signaling axes ^6^, metabolic pathways ^7^, and the overall epigenetic landscape ^8^. Such changes eventually trigger modifications in the cardiomyocyte genetic program to increase cell contractility and favor cell survival ^9^. Reactivation of the fetal cardiac genetic program is a hallmark of the pathological heart, as observed in pressure overload-induced hypertrophy ^10^.

Besides transforming the transcriptional landscape, post-transcriptional events, such as alternative splicing (AS), RNA editing and transport, translation and degradation are also affected in cardiac disease ^11^. In particular, changes in AS in the pathological heart lead to alterations in the expression level of numerous sarcomeric and ion channel genes associated with cardiac pathologies, being determinants of cardiomyopathies and, eventually, of heart failure (HF) ^12^. The mechanisms presiding over RNA homeostasis permit prompt cellular adaptation to changes in the external environment. This process is mainly controlled via RNA binding proteins (RBPs) that physically associate with and guide RNAs through all steps of post-transcriptional control. Given the importance of RBPs in RNA metabolism, their expression must be tightly controlled.

Among the RBPs, heterogeneous nuclear ribonucleoproteins (hnRNPs) represent a large family of proteins that comprise nuclear ribonucleoprotein complexes consisting of RNA and proteins, and help control mRNA dynamics ^13^. Under stress conditions, hnRNPs may relocalize to a different cellular compartment, undergo post-transcriptional modification, and/or exhibit altered protein expression; all of these events can affect the metabolism of essential transcripts needed for cell adaptation ^14,15^. Indeed, the mislocalization and deregulation of hnRNPs has been associated with the onset of several pathologies, including cancer, Alzheimer’s disease and fronto-temporal lobe dementia ^16^.

Emerging data suggest that RNA homeostasis can be controlled by mechanical cues ^17–19^, but the molecular basis of such processes is largely unknown. One hypothesis is that mechanical stress arising from the surrounding environment could affect RNA dynamics by either altering RBP function or localization ^20,21^. However, this process requires that components of the RNA processing apparatus respond to mechanical cues generated during ECM remodeling.

In this study, we show ischemic and chronic cardiac pathologies are accompanied by the altered expression of the RBP heterogeneous nuclear ribonucleoprotein C (hnRNPC). The protein adopts a peculiar sarcomeric distribution and associates to the translation machinery upon pathological ECM remodeling. Moreover, we provide evidence that hnRNPC intracellular localization can be controlled by the increased biomechanical stress generated by cardiac ECM remodeling and show the relocation of a fraction of the protein from the nucleus is sufficient to affect the AS of transcripts coding for components of the mechanosensitive Hippo pathway. This pathway is known to be heavily involved in the progression of cardiac diseases ^22,23^.

Therefore, we propose hnRNPC acts as a mechanosensitive switch affecting RNA metabolism in the pathological heart.

## RESULTS

### The RNA-binding protein hnRNPC is upregulated in the diseased heart

The intense ECM remodeling occurring in cardiac pathologies can alter tissue integrity and impair organ function ^5^, but the underlying molecular mechanisms are unclear. We first asked whether this phenomenon was associated with a common genetic signature in both ischemic and non-ischemic heart diseases. To do so, we analyzed four published expression profiling array datasets obtained from ischemic (mouse myocardial infarction, MI) ^24^ and non-ischemic (mouse and human HF) ^25–27^ cardiac diseases and looked for common molecular function terms among the upregulated genes in the pathological heart (**Supplementary Data 1**).

We found that 11-17% of the significantly upregulated genes in MI and HF hearts (FC>1.5) shared the RNA binding (GO:0003723) annotation (**Fig. 1a; Supplementary Data 1**). When browsing the genes encoding RBPs, we found 82 RBPs that were upregulated in both mouse datasets and four that were upregulated in both human heart failure (HF) datasets. We identified hnRNPC as the only common upregulated gene among all four datasets (**Fig. 1b**). hnRNPC is a core ribonucleoprotein that is ubiquitously distributed and best known for its role as a splicing regulator ^28–30^. Increased levels of this RBP have been associated with several pathological conditions ^31–33^. Its upregulation in diseased cardiomyocytes has been recently confirmed by single cell RNA-sequencing in a model of ischemia-reperfusion ^34^.

**Figure 1:**
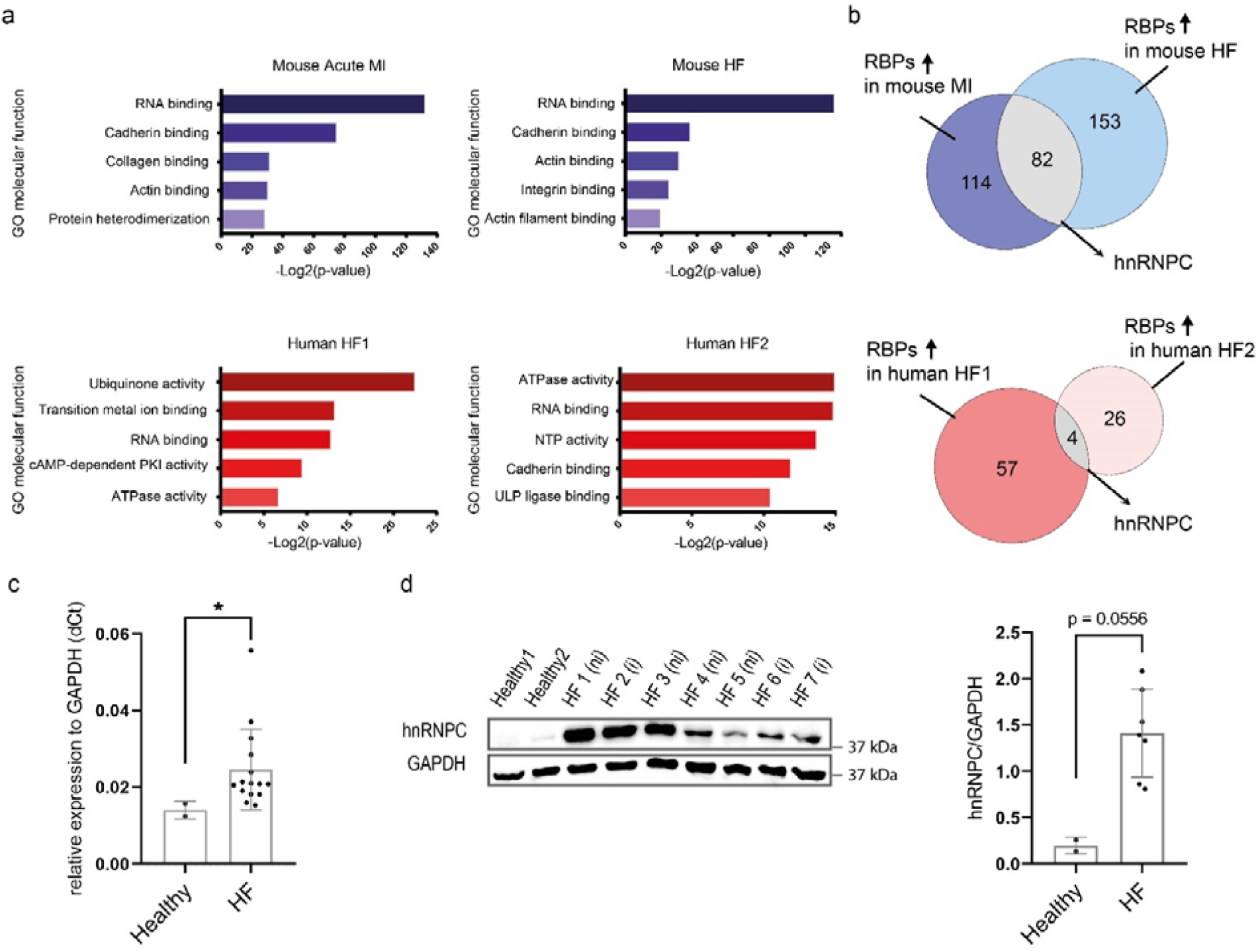
The RNA-binding protein hnRNPC is upregulated in the diseased heart. **a)** Top: Gene ontology (GO) annotation of the significantly upregulated genes (FDR < 0.1; FC > 1.5) identified by expression profiling array of murine hearts (left ventricle) exposed to myocardial infarction (48 h post-MI) as compared to sham operated (left) and of MerCreMer (MCM) murine hearts with peak dysfunction (day 10 post-tamoxifen treatment) as compared to control (right). Bottom: Gene ontology (GO) annotation of the genes found significantly upregulated by expression profiling array analysis of failing (HF) human heart as compared to healthy human heart in two datasets: HF1 (left) (FDR < 0.05; FC > 1.8) and HF2 (FDR < 0.1; FC > 1.5) (right). **b)** Top: Genes found upregulated by *in silico* analysis of profiling array data 48 h post-MI and in HF mouse heart tissues that encode for proteins involved in RNA binding (GO: 0003723). Bottom: Genes found upregulated by *in silico* analysis of profiling array data of HF1 and HF2 encoding for proteins involved in RNA binding (GO: 0003723). **c)** RT-qPCR analysis of hnRNPC RNA expression in HF (N = 15; n = 2) compared to healthy human hearts (N = 2; n = 2). **d)** Western blot analysis (left) and relative quantification (right) of hnRNPC protein expression in samples of HF (1-7) (N = 7; n = 3) and healthy (1-2) (N = 2; n = 3) heart tissues. HF (i) and HF (ni) indicate ischemic and non-ischemic samples, respectively. GAPDH was used for total protein loading normalization. Data are presented as mean ± S.D.; *p < 0.05; p-values were determined by Mann-Whitney test. See also **Supplementary Fig. 1** and **Supplementary Data 1**.

Consistently, our *in silico* expression array analyses showed that hnRNPC was upregulated 1.52 fold 48 h after MI and 1.77 fold in HF in mice, and from 1.68 to 4.2 fold in HF patients (**Supplementary Fig. 1a; Supplementary Data 1**).

Given the consistent alteration in the expression level of hnRNPC in both ischemic and non-ischemic cardiac pathologies, we further analyzed its expression in heart tissue specimens obtained from a cohort of 17 patients diagnosed with end-stage HF and undergoing cardiac transplantation. As a healthy control, we used non-transplantable hearts from deceased (healthy) donors (**Supplementary Fig. 1b, c; Supplementary Table 1**). Both RT-qPCR and western blot analyses confirmed that hnRNPC gene (**Fig. 1c**) and protein (**Fig. 1d**) expression were reproducibly upregulated in failing human hearts. These results support that hnRNPC expression is upregulated in HF. This phenomenon seems to happen independently of the etiology of the pathology, since no significant difference is found in ischemic vs non-ischemic heart diseases (Supplementary Fig. 1d, e).

### hnRNPC interacts with components of the myofibril in pathological heart

The participation of hnRNPs to pre-mRNA processing and mRNA dynamics is defined by their cellular localization and by their interaction with specific interactors. To investigate hnRNPC binding partners and how these might change upon HF, we pulled down the endogenous protein in human end-stage failing and healthy heart samples and quantitatively analyzed the immunoprecipitated complexes by tandem mass tag spectrometry (TMT-MS, **Fig. 2a**). We identified 194 unique proteins interacting with hnRNPC in failing hearts, and 127 in healthy samples: only 55 interactors were common to both conditions (**Fig. 2b; Supplementary Data 2; Supplementary Methods**).

**Figure 2:**
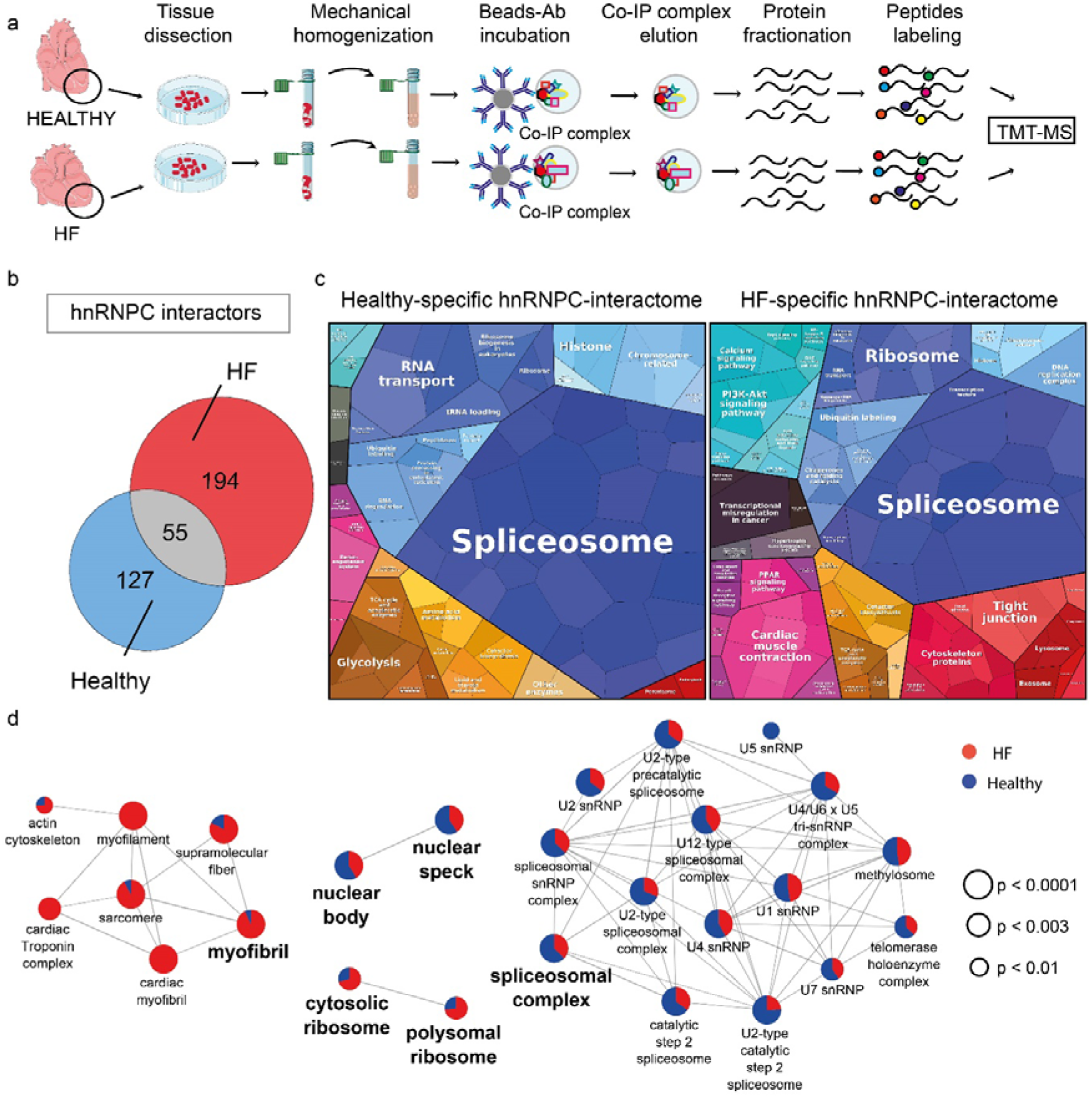
hnRNPC interacts with components of the myofibril in pathological heart. **a)** Graphical representation of the strategy adopted to identify hnRNPC interactors in the healthy human and failing heart (HF) *in vivo* by endogenous protein immunoprecipitation followed by tandem mass tag spectrometry (TMT-MS). **b)** The number of unique and shared proteins found in the hnRNPC interactome in human healthy (N=2) and HF (N=2) heart tissue by TMT-MS. **c)** Proteomaps (based on the KEGG database) representing the quantitative composition of the hnRNPC interactome in healthy and HF human hearts with a focus on protein function. Proteins are weighted for their abundance in the immunoprecipitated samples. **d)** Network representation of the cellular components Gene Ontology (GO) categories enriched in the hnRNPC interactome in healthy (red) and HF (blue) heart tissue. The size of the node indicates the Bonferroni adjusted Term p-values. See also **Supplementary Fig. 2** and **Supplementary Data 2**.

We weighted the specific HF and healthy interactor proteins for their abundances in the hnRNPC-IP complexes and used the resulting data to build proteomaps based on the KEGG database ^35^. This approach allowed us to visualize how the hnRNPC interactome composition changes in the pathology with a focus on the function of its binding partners (**Fig. 2c** and **Supplementary Data 2**). We found that hnRNPC mainly interacted with components of the spliceosome complex in both healthy and diseased hearts, consistent with its acknowledged function in RNA splicing ^29,30^. More interestingly, HF-specific interactors returned a common annotation for cardiac muscle contraction and cytoskeletal proteins (**Fig. 2c**), suggesting that hnRNPC might display a distinct localization in the pathological heart (**Supplementary Fig. 2a**).

We next performed gene ontology analysis of hnRNPC interactors in healthy and diseased heart to identify which cellular component they belonged to. This analysis revealed that HF-specific hnRNPC interactors included proteins belonging to the myofibril. For example, components of the myofilament, sarcomere and the cardiac troponin complex were found bound to the RBP, almost exclusively in HF (**Fig. 2d; Supplementary Fig. 2b**). Together with the annotation for cytosolic and polysomal ribosome, which was significant for hnRNPC interactors in HF samples, these data support a differential localization of the protein in the pathological heart.

### hnRNPC localizes to the Z-disk of the sarcomeric apparatus in the pathological heart

hnRNPC is predominantly localized in the nucleus. Given its interaction with components of the contractile apparatus and ribosomal proteins in the failing heart, we hypothesized hnRNPC could have a different localization under this condition. To confirm our hypothesis, we used confocal microscopy to investigate hnRNPC intracellular localization in the human myocardium. Besides being localized in the nucleus of cardiomyocytes, we also detected hnRNPC in the cytoplasm of the contractile cells in the failing heart, with a characteristic striated pattern closely following the distribution of cardiac troponin T (**Fig. 3a**). Using structured-illumination microscopy (SIM) to resolve the subcellular distribution of the protein in the human pathological heart, we observed that hnRNPC was localized at the Z-disc of the sarcomeres in HF (**Fig. 3b; Supplementary Fig. 3a**) with a lateral resolution of ∼125 nm. The shuttling of a fraction of the protein in the diseased heart was confirmed by western blot analysis of nuclear and cytoplasmic extracts. This experiment also indicated a significant reduction in hnRNPC nuclear expression in the pathological heart (**Supplementary Fig. 3b**).

**Figure 3:**
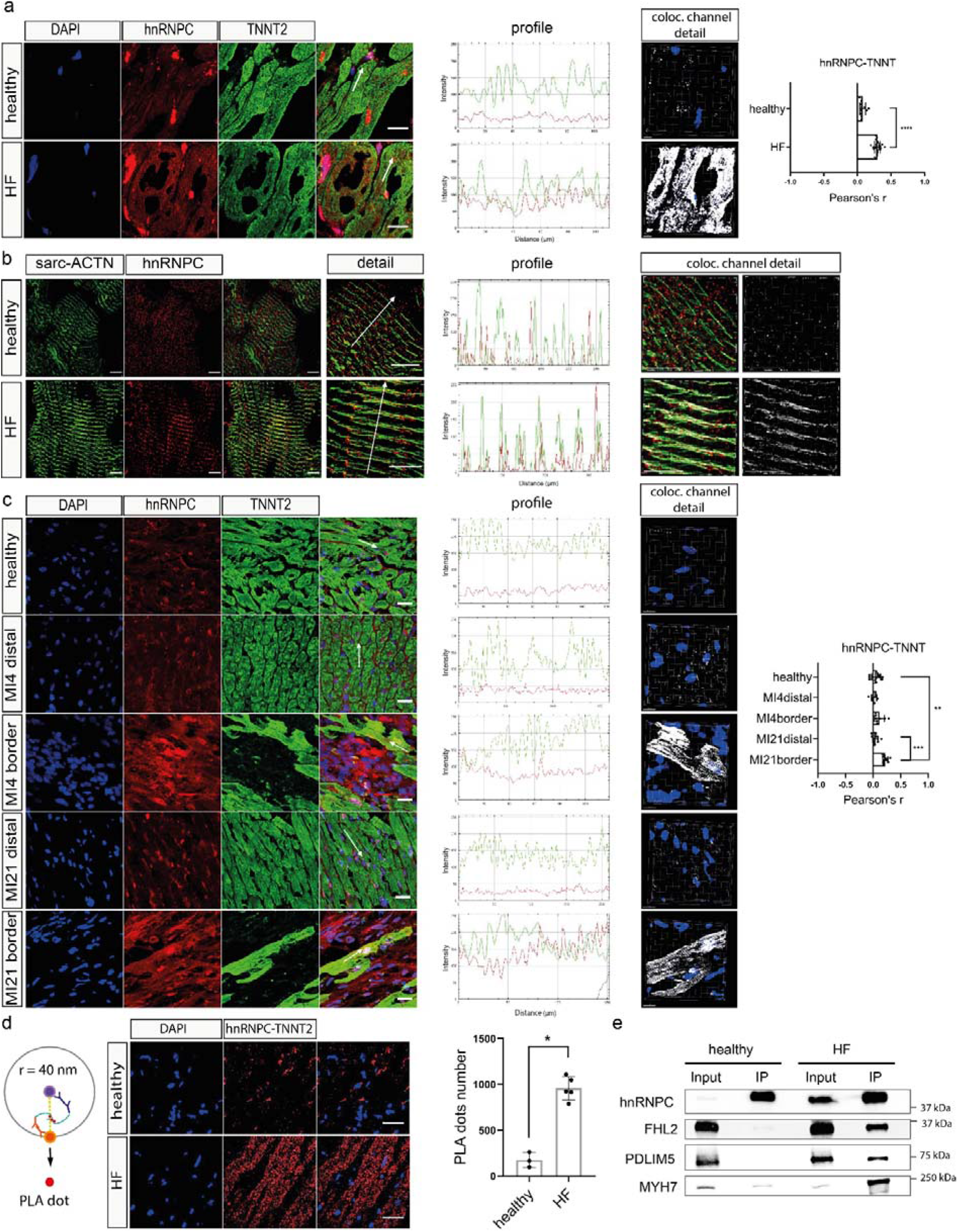
hnRNPC is localized at the Z-disk of the sarcomeric apparatus in the pathological heart. a) Representative confocal microscopy images depicting hnRNPC (red) localization in healthy (N=2) or failing (HF) (N=3) human heart tissue. Cardiomyocytes are stained with cardiac troponin T2 antibody (TNNT2, green) and the nuclei are counterstained with DAPI (blue). The image analysis shows hnRNPC (red) distribution across the sarcomere, as identified by TNNT2 staining. The area analyzed is indicated by the white arrow in the microscopic image. The detail shows the colocalization (white) of red and green channels. Pearson’s coefficient was calculated by Imaris (n ≥ 11). Data are presented as mean ± S.D.; ****p < 0.0001; Mann-Whitney test. Scale bar = 20 μm; Scale bar of detail = 5 μm. **b)** Representative super-resolution image and image analysis of hnRNPC (red) sarcomeric distribution in failing HF. The sarcomere is counterstained by sarcomeric alpha actinin (sarc-ACTN, green). The area analyzed is indicated by the white arrow in the microscopic image. The detail shows the colocalization (white) of red and green channels. Scale bar = 5 μm. **c)** Representative confocal images showing the localization and relative profile of hnRNPC (red) in healthy and at the infarction proximity (border zone), or distal areas of infarcted mouse heart 4- and 21-days post-MI. Cardiomyocytes are stained with cardiac troponin T (TNNT2, green) and the nuclei are counterstained with DAPI (blue). The area analyzed is indicated by the white arrow in the microscopic image. The detail shows the colocalization (white) of red and green channels. Pearson’s coefficient was calculated by Imaris (n ≥ 8). Data are presented as mean ± S.D.; ***p < 0.001; **p < 0.01; Mann-Whitney test. Scale bar = 10 μm. **d)** Representative confocal images and relative quantification of the results obtained from Proximity Ligation Assay (PLA) by using antibodies against hnRNPC and TNNT2 in healthy (N = 3; n ≥ 12) and HF (N = 5; n ≥ 13) human heart tissue. Red dots identify areas where the proteins are found to be within 40 nm of each other. The nuclei are counterstained with DAPI (blue). Data are presented as mean ± S.D.; *p < 0.05; Mann-Whitney test. Scale bar = 50 μm. **e)** Western blot analysis of three sarcomeric proteins (FHL2, PDLIM5 and MYH7) in hnRNPC immunoprecipitated samples (IP) or total lysate (input) in healthy and failing human heart (HF). See also **Supplementary Fig. 3** and **Supplementary Data 3**.

To understand whether hnRNPC sarcomeric distribution can be considered a common feature of cardiac pathologies, we analyzed its localization in a mouse model of MI generated by left anterior descending (LAD) coronary artery ligation in C57BL/6 mice ^36^ (**Supplementary Fig 3c, d**). We stained the infarction border and the distal zone of the infarcted myocardium with an antibody directed against hnRNPC at day 4 post-MI, when the tissue is stormed by intense inflammatory response and at day 21 post-MI, when the acute phase is resolved and the scar is stabilized ^5^. As expected, hnRNPC showed a characteristic nuclear localization in both the healthy and ischemic heart. When co-stained with troponin T2 (TNNT2), hnRNPC displayed a clear sarcomeric pattern in cardiomyocytes located at the infarct border zone 21 days post-MI. A similar pattern could be detected at the earlier time-point (day 4 post-MI). Image analysis clarified the co-localization between hnRNPC and TNNT2 was only significant at day 21 post-MI as compared to sham-operated, day 4 post-MI. No such phenomenon was found far from the infarct, at the distal zone (**Fig. 3c**). These results suggest that hnRNPC translocates to the sarcomeres in cardiomyocytes at the border zone of the infarction where it interacts with components of the sarcomere. This process seems to start right after the ischemic insult and consolidates during the chronic MI stage. To further confirm hnRNPC sarcomeric localization in failing hearts, we performed a proximity ligation assay (PLA) with TNNT2 (**Fig. 3d; Supplementary Data 2; Supplementary Methods**). We found that hnRNPC and TNNT2 are located within 40 nm of each other in HF and that the number of interactions, as indicated by PLA dots, was significantly lower in healthy hearts (**Fig. 3d**). We also confirmed the physical interaction between hnRNPC and other sarcomeric proteins (FHL2, PDLIM5, MYH7) identified by TMT-MS in HF but not in the healthy heart by co-immunoprecipitation analysis (**Fig. 3e**).

In order to further corroborate hnRNPC interaction with FHL2 and PDLIM5, we pulled the two endogenous proteins down in 2 HF samples and resolved the interacting proteins (**Supplementary Data 3**). hnRNPC presence was confirmed in both IP preparations by MS analysis and by western blot (**Supplementary Fig. 3e; Supplementary Data 3**). Interestingly, FHL2, PDLIM5 and hnRNPC interactomes returned 143 common binding partners in the failing heart (**Supplementary Fig. 3f**) with a common annotation for actin cytoskeleton and myofibril component (**Supplementary Fig. 3g; Supplementary Data 3**).

Altogether, these results indicate that hnRNPC localizes to the contractile apparatus in both ischemic and non-ischemic diseased hearts. The sarcomeric distribution of hnRNPC in HF suggests this protein might have distinct functions in physiological and pathological conditions.

### hnRNPC binds transcripts encoding components of cardiomyocytes contractile apparatus in HF

To understand the role of hnRNPC at the sarcomere in HF, we next determined which transcripts it physically binds. To do so, we carried out RNA-binding protein-immunoprecipitation (RIP) followed by high-throughput sequencing (seq) in three human HF samples (**Fig. 4a; Supplementary Methods**). We first confirmed the specificity of the antibody in control RIP-seq experiments in which a mouse IgG antibody showed no hnRNPC immunoprecipitation (**Supplementary Fig. 4a**). Next, we found that the distribution of reads on genomic regions (intron/exon, 5’UTR, 3’UTR, CDS) in the hnRNPC-IP library mostly mapped to intronic regions (63%), followed by coding DNA sequences (CDS, 22%), 3’ (13%) and 5’ (2%) untranslated regions (UTRs) (**Fig. 4b**). After browsing the POSTAR2 database, we confirmed that 2,487 out of 2,629 RNA targets found in our RIP-seq experiment harbored at least one binding site for hnRNPC ^37^ (**Supplementary Fig. 4b**).

**Figure 4:**
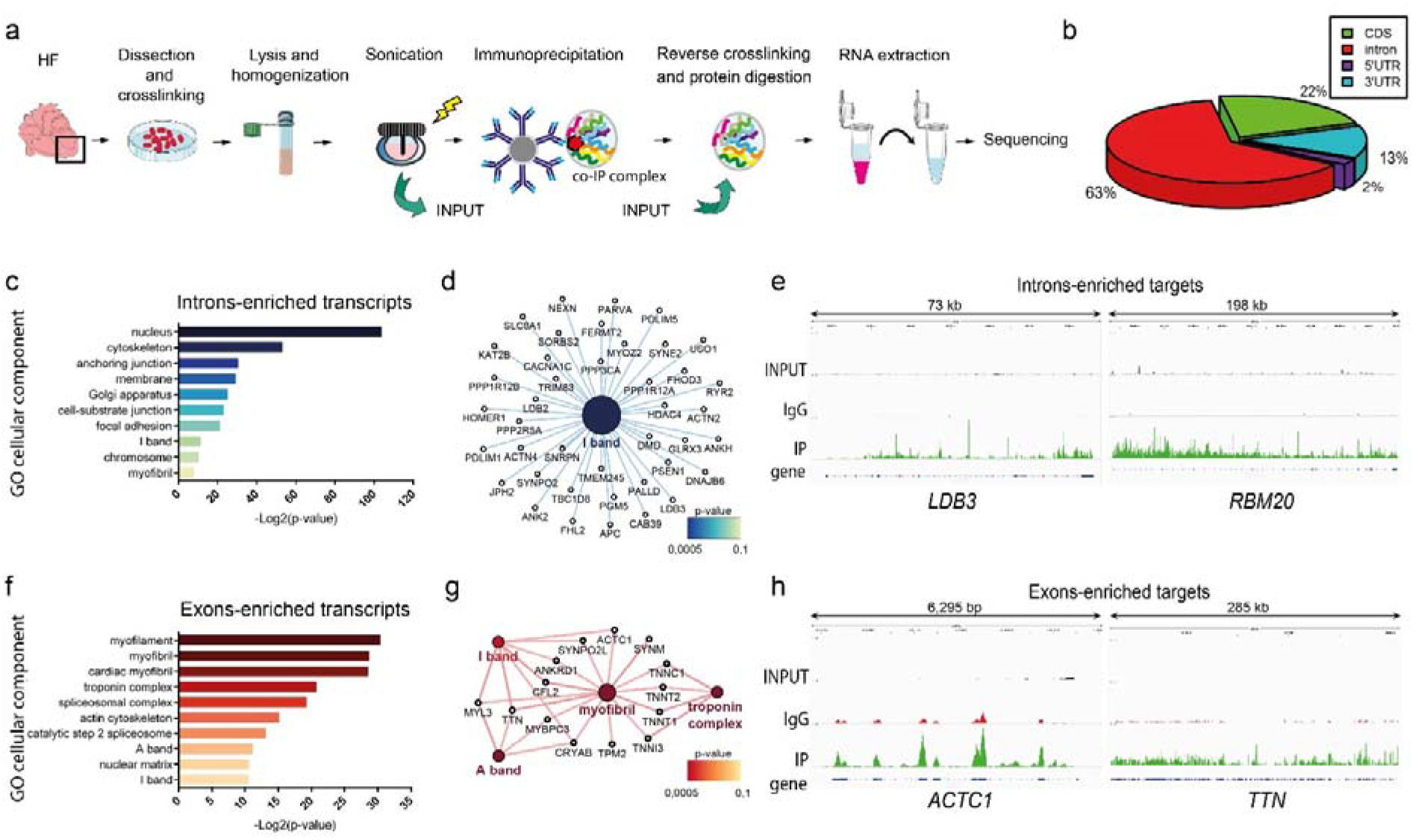
hnRNPC binds transcripts encoding components of cardiomyocytes contractile apparatus in HF. **a)** Graphical representation of the strategy adopted to identify hnRNPC RNA targets in human heart failure (N = 3) (HF) *in vivo* by endogenous RNA immunoprecipitation followed by sequencing. **b)** Genomic distribution of the read counts showing the percentage of reads mapping on specific genomic regions (intron/exon, 5’UTR, 3’UTR, CDS) in immunoprecipitated samples (N = 3). **c)** Cellular components Gene Ontology (GO) categories significantly enriched in intron-enriched transcripts (log2 intron/exon ratio > 1) bound to hnRNPC in HF. **d)** GO network of I band cellular components and the intron-enriched transcripts within this network that bind hnRNPC (p-value corrected with Bonferroni = 0.00037). The size of the node is proportional to the number of hnRNPC targets belonging to the ontology. **e)** Individual gene representation of intron-enriched hnRNPC targets: read coverage is displayed in INPUT (black), IgG (red) and IP (green) samples. **f)** Cellular components GO categories significantly (p-value < 0,05) enriched in exon-enriched transcripts (log2 intron/exon ratio < -1) bound to hnRNPC in HF. **g)** GO network of cellular components of interest enriched (Group p-value corrected with Bonferroni = 9.43E-07) in exon-enriched RNAs that bind hnRNPC in HF. The size of the node is proportional to the number of hnRNPC targets belonging to the ontology. **h)** Individual gene representation of exon-enriched hnRNPC targets: read coverage is displayed in INPUT (black), IgG (red) and IP (green) samples. See also **Supplementary Fig. 4** and **Supplementary Data 4**.

Next, we calculated the number of reads mapping to a specific region (intron/exon, 5’UTR, 3’UTR, CDS) for each gene. By this approach, we could define two groups of targets: the transcripts enriched in intronic and those enriched in exonic reads (exon, CDS, 5’ and 3’ UTRs) (**Supplementary Fig. 4c; Supplementary Data 4; Supplementary Methods**). The large proportion of intronic hits is consistent with the documented role of hnRNPC in pre-mRNA splicing ^28^. Although RBPs can regulate pre-mRNA splicing by binding to exonic splicing enhancers or silencers ^38^, we hypothesized that the exon-enriched targets found in the pathological heart could represent mature RNAs that might be informative as to hnRNPC activity at the sarcomere.

We then performed gene ontology analysis of hnRNPC-targets to understand to which cellular component they belonged to. Here, we identified that the intron-enriched targets (1,681) mostly belonged to nucleus, cytoskeleton and adhesion categories, while 18.8% of all annotated I-band components were hnRNPC targets (**Fig. 4c, Fig. 4d**). Among the intron-enriched targets belonging to the I-band category, we found transcripts such as LDB3, CACNA1C, DMD, PDLIM5, for which an alteration in AS has been previously assigned to the occurrence of cardiac pathologies ^12,39^. Moreover, we found splicing regulators among the hnRNPC targets (including RBM20, MBNL2, CELF1-2, RBFOX1-2) that are involved in heart development and disease ^40,41^. These findings indicate that hnRNPC could exert both a direct and indirect role in splicing control of sarcomeric transcripts affecting the post-transcriptional regulation of other RBPs. Selected hnRNPC intron-enriched targets, showing high read coverage on intronic regions, as obtained by Integrative Genomics Viewer (IGV), are shown in **Fig. 4e** and in **Supplementary Fig. 4d**.

We performed a similar cellular component enrichment analysis of the hnRNPC exon-enriched targets and identified a significant enrichment for the myofibril, troponin complex, A and I bands of the sarcomeres (**Fig. 4f**). Titin (*TTN*), cardiac muscle alpha actin (*ACTC1*) and troponin I (*TNNI3*) contractile apparatus transcripts were found among hnRNPC targets (**Fig. 4g; Supplementary Fig. 4e**). When we analyzed the enriched biological processes by these exon-enriched hnRNPC targets, we found that these targets were mainly involved in muscle structure development (22), filament sliding (9), contraction (14) and myofibril assembly (9) (**Supplementary Fig. 4f**). By comparison, the intron-enriched targets were principally associated with the regulation of signal transduction (472), cell communication (513), cytoskeletal organization (209) and heart development (113) (**Supplementary Fig. 4g**).

These data demonstrate that hnRNPC interacts with intron-enriched transcripts, confirming its well-known function as a splicing regulator. On the other hand, hnRNPC also binds exon-enriched transcripts that are enriched in RNAs encoding myofibril components. This finding suggests a possible implication of hnRNPC in translational regulation of selected targets upon its relocalization to the sarcomeres in HF.

### ECM pathological remodeling induces hnRNPC association to the translation machinery at the sarcomere

Because hnRNPC exon-enriched targets comprised mainly mRNAs encoding sarcomeric proteins and we found hnRNPC localized at the sarcomere in HF, we queried the role of hnRNPC at this site. *TTN, ACTC1* and *TNNI3* transcripts, found among hnRNPC exon-enriched targets, are translated at the sarcomeres in the rat heart, in a process dubbed as ‘localized translation’ ^42,43^. Localized translation allows for the spatio-temporal control of contractile protein synthesis at the sarcomeric site ^43,44^. Previous reports have shown ribosomes are present at the sarcomeres in the rat heart, where localized translation ensures that contractile proteins are readily replaced during sarcomere remodeling ^43,45^. Consistently, our TMT-MS analysis identified nine ribosomal proteins in the hnRNPC interactome in the failing heart (**Supplementary Fig. 5a**) which accounted for the ribosome annotation found in GO cellular component analysis (**Fig. 2d**). To confirm the presence of the ribosomes at the sarcomeric structure, we labeled the evolutionarily conserved RPS6 ribosomal protein (which was a potential hnRNPC interactor in our TMT-MS analysis) in healthy and HF cardiac tissue sections. We found a typical sarcomeric cross-striated pattern, as confirmed by TNNT2 decoration, in both groups (**Supplementary Fig. 5b**). We observed a similar pattern in the sarcomeres of cardiomyocytes residing both at the distal and at border zones of myocardial infarction, as well as in healthy cardiomyocytes in the murine heart (**Supplementary Fig. 5c**).

As we found that hnRNPC relocated to the sarcomeres in HF and confirmed the presence of the ribosomes at this site in the human heart, we asked whether sarcomeres could be a site of localized translation in human cardiomyocytes. Therefore, we derived contractile cardiomyocytes from human induced pluripotent stem cells (iPSC-CMs) through an established protocol ^46^. In differentiated cells, we first confirmed RPS6 presence at the sarcomeres by co-localization analysis of the protein with cardiac TNNT2 (**Fig. 5a**). Next, we performed a ribopuromycylation assay that exploits puromycin mimicry of aminoacylated tRNA to visualize the active sites of translation (**Supplementary Fig. 5d**). Here, puromycin (PMY) could be incorporated in the contractile apparatus of human iPSC-CMs, thus demonstrating human cardiomyocyte sarcomeres are active sites of protein translation (**Fig. 5b**).

**Figure 5:**
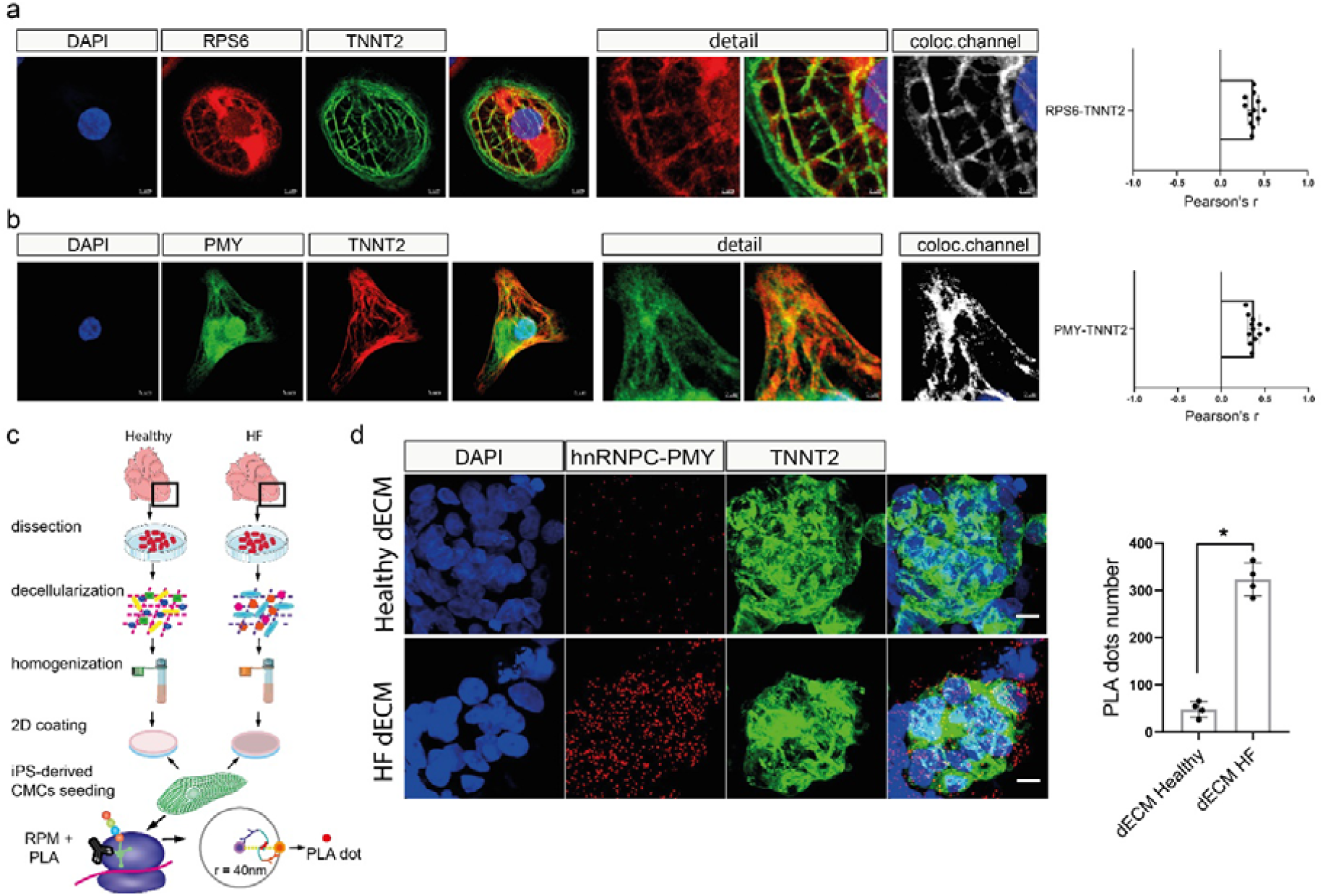
hnRNPC associates to the translation machinery at the sarcomere following ECM pathological remodeling. **a)** Representative confocal image of ribosomal protein S6 (RPS6, red) localization in iPSC-derived contractile cardiomyocytes (N = 3). Sarcomeric units are identified by cardiac troponin T (TNNT2, green) staining and nuclei are counterstained with DAPI (blue). The detail shows a magnified area of a single cardiomyocyte in which the colocalization of red and green channels has been quantified using Imaris software (n = 12). Data are presented as mean ± S.D. Scale bar = 5 μm; Scale bar of detail = 2 μm. **b)** Representative confocal microscopy images of single iPSC-derived cardiomyocytes labeled with anti-puromycin antibody (PMY, green) after puromycin incorporation (N = 3). Sarcomeric units are identified by cardiac troponin T (TNNT2, red) staining and the nuclei are counterstained with DAPI (blue). Colocalization analysis of the green and red channels was quantified by Imaris software (n = 12). Data are presented as mean ± S.D. The detail shows a magnified area of a single cardiomyocyte. Scale bar = 5 μm; Scale bar of detail: 2 μm. **c)** Description of the ribopuromycilation and Proximity Ligation Assay (PLA) used to investigate the occurrence of localized translation at the sarcomeres in iPSC-derived cardiomyocytes grown onto decellularized matrices obtained from either healthy or failing human hearts. **d)** Representative confocal microscopy images of PLA using antibodies against PMY and hnRNPC in iPSC-derived beating cardiomyocytes grown onto HF or healthy decellularized heart matrices. The cardiomyocytes were labeled with cardiac troponin T (TNNT2, green). Red dots indicate that the proteins are within 40 nm of each other. The nuclei are counterstained with DAPI (blue). Data are presented as mean ± S.D. (N = 4; n = 6). *p < 0.05; Mann-Whitney test. Scale bar = 10 μm. See also **Supplementary Fig.5**.

Next, we queried whether hnRNPC associates to the translation machinery when localized at the sarcomeres in the pathological heart. To do this, we adopted an original reductionist approach that proved effective at reproducing some key mechanical features of pathological cardiac ECM remodeling by decellularizing healthy and diseased human heart tissue (dECMs) ^23,47^. We used the homogenized dECMs to coat the surface of cell culture plates. Next, we seeded contractile iPSC-CMs on healthy and pathological dECMs and combined ribopuromycilation with PLA assay to investigate whether hnRNPC localized close to the active translation sites, as indicated by PMY staining (**Fig. 5c**). The analysis showed the presence of PLA dots, corresponding to hnRNPC–PMY proximity, in TNNT2-positive cells cultured on pathological dECMs but not in iPSC-CMs grown on healthy dECMs (**Fig. 5d**). These data confirm that sarcomeres are active sites of RNA translation and that hnRNPC localizes to those sites as a result of cardiac ECM pathological remodeling.

### hnRNPC intracellular localization is controlled by cell spreading and cytoskeletal tension

Thus far we have shown that cardiac ECM pathological remodeling induces hnRNPC shuttling to the active sites of translation at the sarcomere. Pathological remodeling is a complex phenomenon modifying the chemical composition and the mechanical properties of cardiac ECM^5,23^. We tried to disentangle the role of mechanical cues in hnRNPC localization. We hence adopted surfaces with the same chemical composition and controlled rigidities (1.5 vs 28 kPa) and repeated the ribopuromycilation assay followed by PLA on iPSC-CMs. The elasticity values chosen do not exactly match those found in the living heart, but are in the same kPa range. More importantly, these values have been shown to induce mechanosensor activation in other cell systems ^48–50^. Here we found hnRNPC localization to the sites of active translation, as detected by PMY interaction, was sensitive to the stiffness of the substrate, since PLA signal was significantly different on substrates with different elasticities. This result suggests that substrate stiffness *per se* can induce hnRNPC shuttling (**Supplementary Fig. 6a**).

In order to corroborate this result, we challenged normal human dermal fibroblasts (NHDFs), a model of highly mechano-responsive cells, with different mechanical stimuli and analyzed hnRNPC distribution. Here, we used a fibrin-based 3D culture system, whose mechanical properties can be tuned in a physiological range by modulating the fibrinogen concentration in the presence of constant levels of thrombin ^51^ (**Supplementary Fig. 6b**). We embedded NHDFs into the hydrogels and analyzed hnRNPC localization after 48 hours. We additionally stained F-actin and cell nuclei to monitor cell ability to spread within the hydrogels, indicative of the degree of cytoskeletal tension. We observed that NHDFs spread well in soft (2 mg/ml and 5 mg/ml fibrinogen) hydrogels, but failed to elongate as the stiffness of the fibrin gel increased (17 mg/ml fibrinogen). At the highest fibrinogen concentration (50 mg/ml), the cells were unable to acquire an elongated morphology (**Supplementary Fig. 6c**). Image analysis demonstrated that hnRNPC nuclear localization correlated with the NHDF volume and with the cellular ability to spread within soft hydrogels, with hnRNPC translocation to the cytoplasm characteristic of cells confined into stiff fibrin gels (**Fig. 6a; Supplementary Fig. 6c**). These data suggest that hnRNPC nuclear localization might be controlled by the ability of cells to develop intracellular tension and spread within the surrounding milieu.

**Figure 6:**
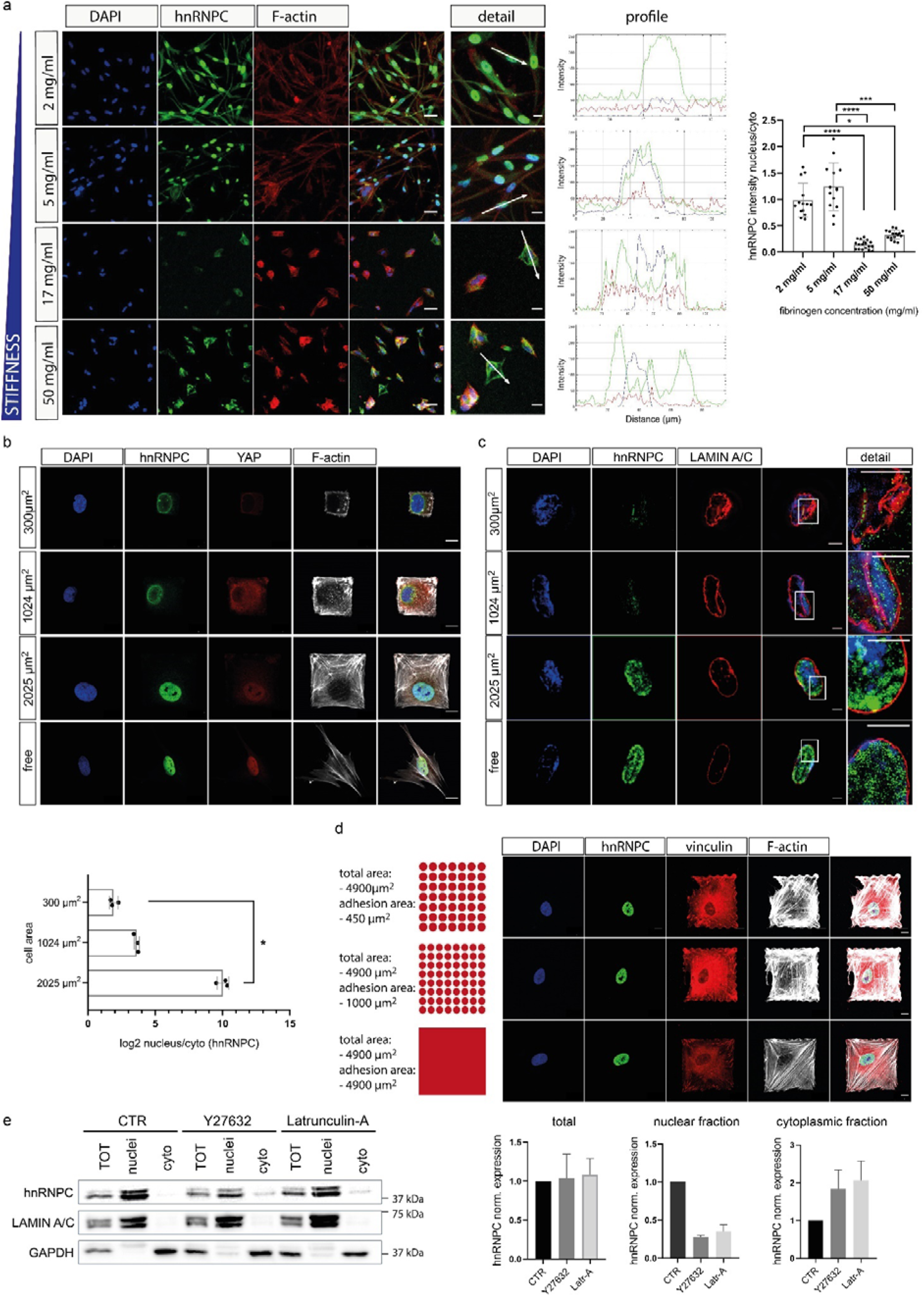
hnRNPC intracellular localization is mechanically controlled. **a)** Left: Representative confocal microscopy images showing hnRNPC (green) expression in normal human dermal fibroblasts (NHDFs) embedded for 48 h in fibrin-based hydrogels with an increasing fibrinogen concentration and stiffness. Cellular F-actin was labeled with Phalloidin (red) and the nuclei were counterstained with DAPI (blue). Image analysis shows the quantification of signal distribution in NHDFs embedded for 48 h in fibrin-based hydrogels with controlled stiffness. Right: Quantification of the hnRNPC in the nucleus/cytoplasm distribution in fibrin-gels with increasing fibrinogen concentration and stiffness. For each experimental condition, data are presented as mean ± S.D. from N ≥ 7 independent fields of acquisition; *p < 0.05; ***p < 0.001; ****p < 0.0001, Kruskal-Wallis test followed by Dunn’s multiple comparison test. **b)** Top: confocal analysis of individual cells grown onto 300, 1024, 2025⍰μm^2^ micropatterns coated with fibronectin. The cells were labeled with anti-hnRNPC (green), Alexa Fluor 647 Phalloidin, anti-Lamin A/C (red) and the nuclei were counterstained with DAPI (blue). Bottom: Barplot representation of hnRNPC nucleus/cytoplasm distribution in single cells seeded onto 300, 1024, 2025 µm^2^ micropatterns (N = 3; n ≥ 8); *p < 0.05; Kruskal-Wallis test followed by Dunn’s multiple comparison test. **c)** Representative super-resolution images of individual cells grown onto 300, 1024, 2025⍰μm^2^ micropatterns coated with fibronectin. The cells were labeled with anti-hnRNPC (green) and anti-Lamin A/C (red). Nuclei were counterstained with DAPI. **d)** Left: Annotation of micropattern properties (cell area and adhesion area) and a schematic top view of the fibronectin distribution in the micropatterns. Red color indicates fibronectin-covered area. Right: Confocal images of single NHDFs grown onto the indicated micropatterns and labeled with anti-vinculin (red), anti-hnRNPC (green), Alexa Fluor 546 Phalloidin and DAPI (blue) (N = 3; n = 5). **e)** Western blot analysis (left) and quantification (right) of hnRNPC expression in cellular (TOT), nuclear and cytoplasmic fractions of NHDFs treated or not (CTR) with Latrunculin-A and Y27632 inhibitors. GAPDH was used for loading normalization of the total and cytoplasmic fractions. Lamin A/C was used to normalize protein loading in the nuclear fraction. Band intensities corresponding to hnRNPC in the nuclear and cytoplasmic fractions were quantified using Bio-Rad Image Lab software. Data are presented as mean ± S.D. (N = 3). See also **Supplementary Fig. 6**.

To corroborate our hypothesis, we used fibronectin-coated micro-patterned surfaces to precisely control single cell area ^49,50,52^. We stained NHDFs with a specific antibody against hnRNPC and used YAP as an example of a mechanosensitive protein whose localization depends on intracellular tension and cell spreading ^49,50^. Confocal analysis showed that hnRNPC localized at the nucleus in cells that were confined to bigger areas (2,025⍰μm^2^) or free to spread. By contrast, the protein shuttled to the perinuclear region of cells constrained on smaller surfaces (1,024 and 300 μm^2^) (**Fig. 6b; Supplementary Fig. 6d**). To resolve the intracellular distribution of the protein, we counterstained the nuclear structure with Lamin A/C and visualized the cells by structured illumination microscopy (SIM). Here we found that hnRNPC egressed the nucleus and acquired a perinuclear localization in constrained cells, with a peculiar distribution close to Lamin A/C nuclear invaginations (**Fig. 6c**). We repeated the same experiment using iPSC-CMs and again observed that hnRNPC nuclear localization was also sensitive to cell spreading in contractile cells (**Supplementary Fig. 6e**).

Having shown that hnRNPC nuclear localization required the cell to spread in 2D and 3D, we next performed in depth experiments to clarify whether the shuttling of the protein was driven by the availability of adhesion sites within the local ECM or by the transmission of intracellular tension. We first designed single cell fibronectin-coated micropatterned arrays that keep the cell area constant (4,900 μm^2^), while reducing the number of fibronectin adhesion spots ^49^. When we decreased the adhesion area (1,000 and 450⍰μm^2^), we saw no significant effect on hnRNPC nuclear localization (**Fig. 6d**).

We concluded that adhesion did not affect hnRNPC localization, and decided to inhibit the transmission of intracellular tension by treating NHDFs with pharmacological inhibitors of either the RhoA/ROCK pathway (Y27632) or F-actin polymerization (Latrunculin A) ^2,53^. We analyzed hnRNPC localization in the nucleus and cytoplasm by cell fractionation followed by western blotting, and detected a significant decrease in nuclear hnRNPC and the concomitant increase in its cytoplasmic expression in cells treated with the pharmacological inhibitors (**Fig. 6e**). These data are consistent with the hypothesis that the inhibition of intracellular tension determines a reduction in hnRNPC nuclear expression and the shuttling of a fraction of the protein to the cytoplasm.

### hnRNPC regulates the AS of genes involved in mechanotransduction

We have so far demonstrated that a fraction of hnRNPC is distributed to the sarcomeres in HF, and its expression in the nucleus is reduced (**Supplementary Fig. 3b**). This data suggests a decreased contribution of the ribonucleoprotein to the splicing process (**Fig. 2c**). Moreover, we found that altered cytoskeletal tension affects hnRNPC subcellular localization. As hnRNPC is associated to the spliceosome and mainly nuclear in physiological conditions, we studied which changes in terms of AS events would cause the reduced contribution of hnRNPC to the splicing process. We therefore knocked down (KD) hnRNPC in NHDFs using two independent siRNAs (**Fig. 7a**) and analyzed the abundance of RNA splicing variants by high throughput genome-wide RNA-seq. We then adopted the multivariate analysis of transcript splicing (MATs) statistical algorithm to quantify the changes in AS driven by hnRNPC downregulation. We identified the differential splicing of 2,032 RNA transcripts (FDR< 0.01 and |ΔPSI| ≥ 0.1) in KD cells, as compared to the scrambled control transfected cells (**Supplementary Data 5; Supplementary Methods**). Consistent with previous reports ^30,54^, most of the splicing events affected by hnRNPC KD consisted of skipped exons (SE: 78,95%), followed by mutually exclusive exons (MXE: 14,85%), alternative 3’-5’ splice sites (A3SS: 3,76% and A5SS: 2,15%) and intron retention (IR: 0,3%) (**Fig. 7b**). Pathway enrichment analysis of the 2,032 differentially spliced genes in hnRNPC KD cells revealed that the splicing of numerous genes involved in cell mechanotransduction, integrin-mediated cell adhesion and focal adhesion was affected (**Fig. 7c**). In particular, 15 components of the Hippo signaling pathway, a mechanosensitive molecular axis involved in heart development ^22^ and diseases ^52,55^, displayed altered AS upon hnRNPC silencing (**Supplementary Data 6**).

**Figure 7:**
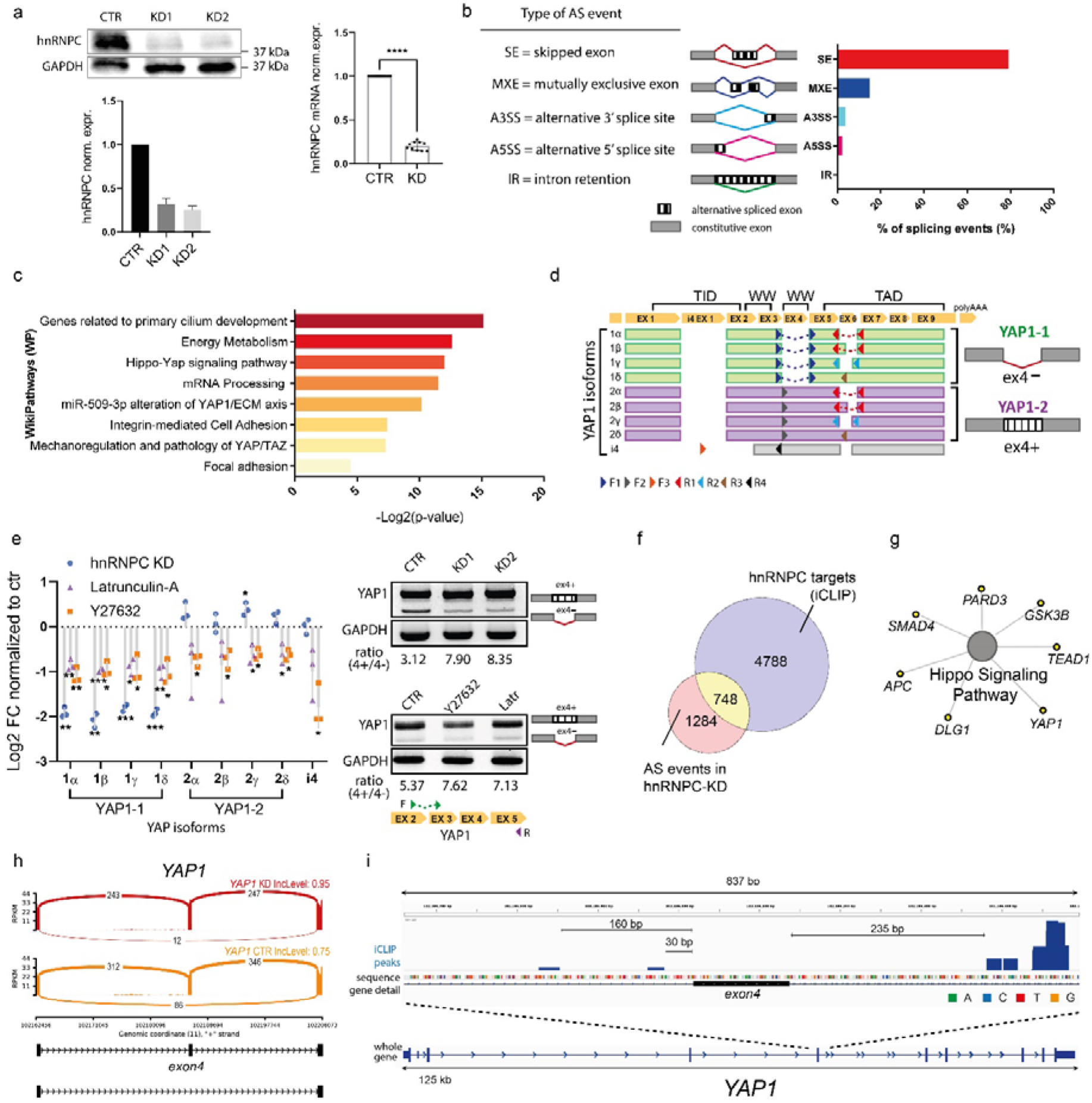
hnRNPC regulates the alternative splicing of genes involved in mechanotransduction. **a)** Left: Western blot analysis (top) and relative quantification (bottom) of hnRNPC protein expression in normal human dermal fibroblasts (NHDF) transfected with hnRNPC siRNA constructs (KD1 and KD2) or with a control siRNA construct (CTR). Protein expression was assessed 96 h after the transfection. GAPDH was used for total protein loading normalization. Data are presented as mean ± S.D. (N = 3). Right: histogram representation of the results obtained by RT-qPCR analysis of hnRNPC RNA expression in NHDFs 96 h after transfection with hnRNPC (KD) or control siRNAs (CTR). Data are presented as mean ± S.D. (N = 10). ****p < 0.0001, Mann-Whitney test. **b)** The percentage of splicing events occurring in hnRNPC KD cells classified into the five major categories of alternative splicing (AS), as obtained by Multivariate analysis of transcript splicing (MATs). **c)** Gene ontology (GO) analysis showing the main pathways found significantly enriched (p-value < 0.05) in genes displaying altered splicing in hnRNPC KD cells using the WikiPathways database. **d)** Schematic representation of protein-coding YAP1 isoforms. Left) Coding regions (CDS) of four YAP1-1, four YAP1-2, and YAP-i4 isoforms are aligned and corresponding protein domains (TEAD-binding domain (TBD), WW domains, TRANS-activating-domain (TID) with N-terminal coiled-coil domain and N terminal PDZ motif) are shown. The arrows represent the approximate binding sites of forward (F) and reverse (R) primers used to detect the specific isoforms. Right) schematic representation of the AS event (exon 4 skipping) that distinguishes YAP1-1 from YAP1-2 isoforms. **e)** Left) Dotplot representation of YAP isoforms mRNA expression (Log2FC) in NHDF upon hnRNPC-depletion (blue) or upon 24h treatment with latrunculin-A (purple) or rock inhibitor (Y27632) (orange). Each result is compared to its relative control. Superimposed bars represent the as mean ± S.D. (N = 3). *p < 0.05; **p < 0.01; ***p < 0.001; One-sample t-test. Right) RT-PCR of YAP1 in NHDF upon hnRNPC depletion (Top) or upon 24h treatment with inhibitors of tension (Latrunculin (Latr) and Y27632) (Bottom). Top band: exon 4-included YAP1 mRNA. Bottom band: exon 4-excluded YAP1 mRNA. **f)** Venn diagram representation of the common transcripts identified as hnRNPC targets by hnRNPC iCLIP-seq data that were also found differentially spliced (AS events) in hnRNPC KD in NHDF cells. **g)** GO network representation of the RNA targets of hnRNPC displaying altered splicing upon hnRNPC silencing and belonging to Hippo signaling Pathway (HSA04390). **h)** Sashimi plot depicting YAP1 skipped exon (SE) AS event (exon 4) in hnRNPC KD NHDFs (red) and controls (orange). The exon inclusion level (IncLevel) is indicated. **i)** Genome browser view of YAP1 gene displaying the iCLIP data (crosslink events per nucleotide) of hnRNP C (blue) (average of N = 3) in proximity of exon 4.

Among the 15 genes belonging to Hippo pathway which displayed altered splicing upon hnRNPC depletion we found the main effector, Yes Associated Protein 1 (YAP1). *YAP1* gene encodes for multiple protein isoforms (**Fig. 7d**). The distinct co-transcriptional fingerprints of the splicing variants are known to depend upon the presence of one (YAP1-1) or two (YAP1-2) WW domains ^56,56^. The exon 4, which encodes for the second WW domain in YAP1-2 isoforms, was found differentially spliced in hnRNPC-depleted cells by RNA-seq. We thus analysed the inclusion of exon 4 in YAP1 transcript upon hnRNPC KD by calculating the relative abundance of YAP1 isoforms including (YAP1-2) or not (YAP1-1) exon 4 by RT-qPCR and by conventional PCR. Interestingly, we found that hnRNPC depletion promotes the inclusion of exon 4 in YAP1 transcript and affects the overall balance among YAP1-1 and YAP1-2 splicing variants (**Fig. 7d, e Supplementary Fig. 7a**). These results were confirmed in hnRNPC-depleted iPSC-CMs (**Supplementary Fig. 7b, c**). Additionally, we obtained similar results by treating NHDF with inhibitors of cytoskeletal tension (**Fig. 7e**). Hence we established that the mechanical displacement of hnRNPC from the nucleus affects the splicing of YAP1 by promoting exon 4 inclusion (**Fig. 7e**).

In order to validate this set of data, we analyzed the expression of the variable exon 7 of CD44 transcript, whose alternative splicing has been proven mechanosensitive ^57^. As expected, here we found the inclusion of variable exon 7 in CD44 transcript increases upon hnRNPC depletion as well as upon treatment with inhibitors of tension in NHDFs (**Supplementary Fig. 7d**). We could only confirm this result in one out of the two iPSC-CMs depleted for hnRNPC (**Supplementary Fig. 7e**). This discrepancy could be explained by the variability in differentiation efficiency in distinct iPSC cultures or by the use of two unrelated siRNAs in the KD experiments.

We also validated the reduced expression of MYLK exon 11 and COL6A3 exon 3 found by RNA-seq in hnRNPC KD cells by RT-qPCR (**Supplementary Fig. 7f, g**).

Next, we compared the differentially spliced genes in hnRNPC KD cells to the RNAs known to harbor hnRNPC binding sites (as documented in the POSTAR database). We found as many as 1,738 among the 2,032 transcripts had at least one hnRNPC binding site and could, in principle, interact with this RBP (**Supplementary Fig. 7h**). In order to check that those were true hnRNPC targets, we performed individual-nucleotide resolution Cross-Linking and ImmunoPrecipitation (iCLIP). This approach allowed us to map precisely hnRNPC-RNA interactions and relate them to the previously detected hnRNPC-dependent splicing events in NHDF cells (**Supplementary Fig. 8a, b; Supplementary Data 7; Supplementary Methods**). A total of 1,470,396 uniquely mapped hnRNPC crosslink events were identified, which cluster into 13,567 binding sites in 5,536 transcripts (**Supplementary Data 7**). As a control, we found the majority of hnRNPC crosslink events to be located within introns, with high coverage of Alu elements, in good agreement with previous results obtained in other cell types ^28,29^ (**Supplementary Fig. 8c, d**). While 58% of the crosslink events mapped to coding sequences, several non-coding hnRNPC targets were also identified, including MALAT-1, a long non coding RNA whose interaction with hnRNPC has already been described ^58^ (**Supplementary Fig. 8e-g**).

Next, we matched the results of AS events in hnRNPC KD cells with those obtained by iCLIP. In particular, we searched for all transcripts directly bound by hnRNPC to determine whether they displayed AS in hnRNPC-depleted cells. We ultimately detected 748 transcripts physically interacting with hnRNPC in NHDFs that were differentially spliced in the absence of the protein (**Fig. 7f; Supplementary Data 7**). Among those, we confirmed the presence of seven components of the Hippo pathway (**Fig. 7g; Supplementary Data 6**), on which hnRNPC binding occurred in intronic regions surrounding exons alternatively spliced in hnRNPC-depleted cells (**Fig. 7h, i; Supplementary Fig. 9a, b)**. This was the case for YAP1 transcript, for which, the RNA-binding protein iCLIP coverage in the proximity of the hnRNPC-dependent exon 4 could be confirmed (**Fig. 7h, i**). The data were validated by independent iCLIP-PCR assay for three selected hnRNPC targets: YAP1, CD44 and MYLK (**Supplementary Fig. 10 a, b**). In summary, these results suggest that hnRNPC is directly involved in the regulation of AS of pre-mRNAs encoding genes involved in mechanotransduction.

### hnRNPC regulates the AS of genes involved in cardiovascular diseases

We next used our iCLIP and alternative splicing analyses to validate the RNA-IP results obtained in the diseased human heart **(Fig. 4)**. We figured this approach might help us unveil the direct targets of hnRNPC in the pathological heart, where the RBP levels and localization are perturbed **(Fig. 1c, d; Fig. 3a, b)**. We matched the three datasets to identify 389 common transcripts which are physically bound by hnRNPC in the failing hearts and undergo alternative splicing in its absence (**Fig. 8a; Supplementary Data 8**). Among them, we again confirmed the presence of the components of the Hippo pathway (**Supplementary Data 6**).

**Figure 8.**
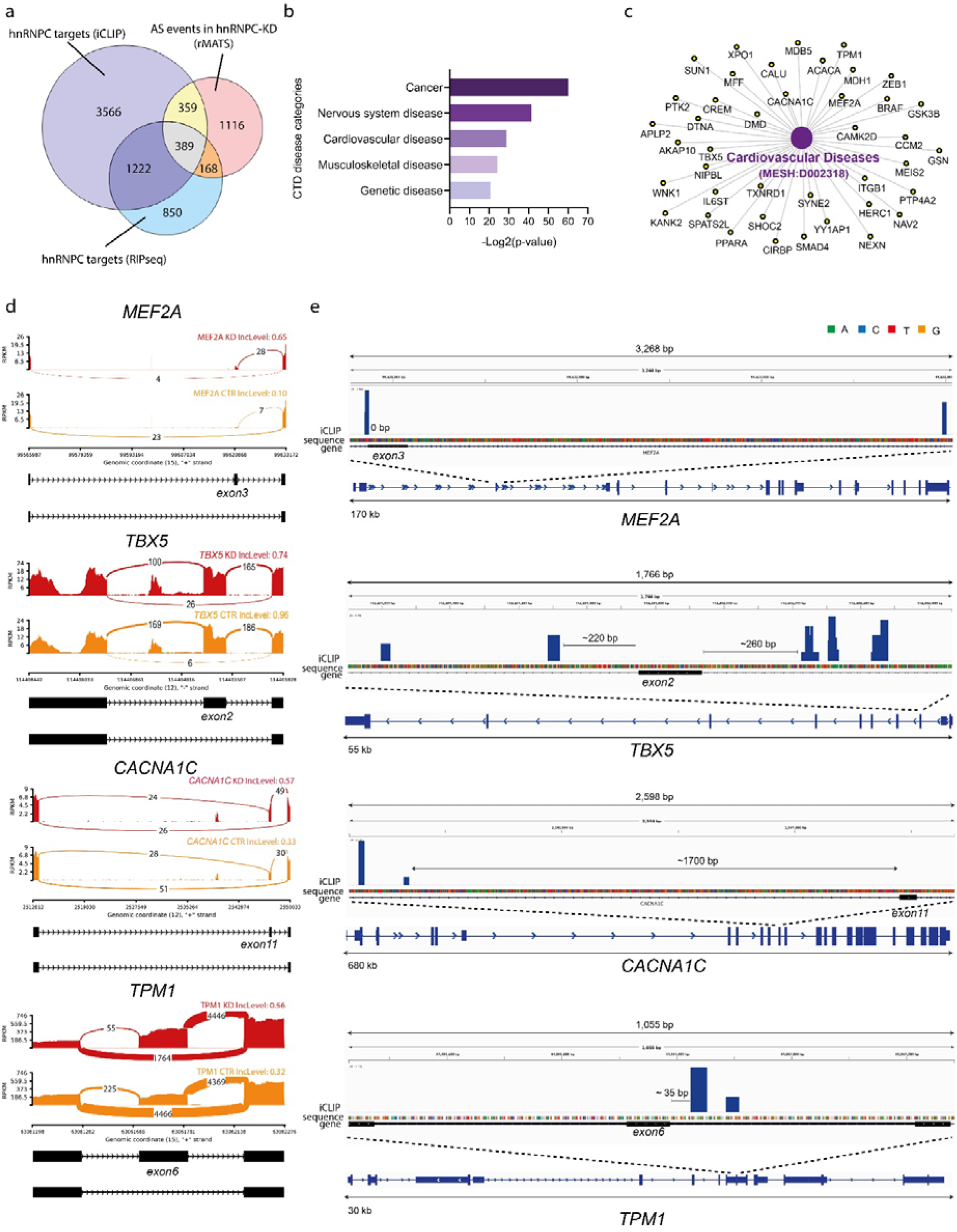
hnRNPC regulates AS of genes involved in cardiovascular diseases. **a)** Venn diagram representation of the transcripts common to the following datasets: iCLIP-seq in NHDFs, RNAseq AS-analysis in hnRNPC KD NHDF and hnRNPC RNA-IP in failing human hearts. **b)** Gene Ontology (GO) analysis of the 389 transcripts common to the three datasets as obtained by Comparative Toxicogenomics Database (CTD). **c)** GO network representation of the transcripts belonging to Cardiovascular Disease (MESH: D002318) category and common to the three datasets analyzed: iCLIP-seq, AS (rMATS) in *HNRNPC* KD NHDF and hnRNPC RNA-IP in failing human hearts. **d)** Sashimi plots depicting representative skipped exon AS events in *MEF2A*; *TBX5*; *CACNA1C* and *TPM1* belonging to Cardiovascular Disease category of the CTD database in hnRNPC KD NHDFs (red) and controls (orange). The exon inclusion level (IncLevel) is indicated. **e)** Genome browser view of individual genes displaying the iCLIP data (crosslink events per nucleotide) of hnRNP C (blue) (average of N = 3) in proximity of specific exons. See also **Supplementary Data 6-8**.

The Comparative Toxicogenomics Database (CTD) annotation of the transcripts targeted for gene– disease associations yielded cardiovascular, cancer and neurological diseases as the main represented categories (**Fig. 8b; Supplementary Data 8**). In particular, 41 hnRNPC direct targets belonging to the cardiovascular disease category displayed AS in hnRNPC KD cells (**Fig. 8c; Supplementary Data 8**). We confirmed the importance of hnRNPC depletion in exon inclusion by visualizing the splice junction of selected transcripts belonging to this category and by highlighting hnRNPC binding sites on those transcripts (**Fig. 8d, e**).

Next, we investigated whether the splicing events determined by hnRNPC depletion *in vitro* are relevant in HF. Hence, we analysed YAP1 exon 4 splicing in our cohort of patients diagnosed with HF and found an increase in exon 4 inclusion in the majority of the samples considered (**Supplementary Fig. 11a**). The variability of this result is expected due to the intrinsic complexity of the cardiac tissue (i.e.: heterogeneous cellular composition of the failing human heart, the presence of fibrotic tissue).

We then adopted the MATs statistical algorithm to quantify the changes in YAP1 exon 4 AS in RNA-seq datasets obtained from patients diagnosed with heart failure and previously published (GSE108157 and GSE141910). Here, we confirmed that failing human hearts display a higher inclusion of exon 4 in *YAP1* transcript, thus supporting our data (**Supplementary Fig. 11b**). Finally, we used the same RNA-seq datasets to further corroborate the clinical relevance of the AS events affected by hnRNPC depletion on a larger scale and found that numerous AS events determined by hnRNPC KD in NHDFs also occurred in the failing heart, independently of the etiology of the pathology (**Supplementary Fig.11c; Supplementary Data 9**).

Taken together, these results indicate that pathological ECM remodeling in the failing heart induces hnRNPC shuttling to the sarcomeres, where the RBP localizes in close proximity to the translation apparatus. A reduction in hnRNPC nuclear localization and in its contribution to the splicing process affects the AS of mechanosensitive transcripts and RNAs involved in cardiovascular diseases.

## DISCUSSION

Cardiac pathologies are characterized by a profound modification of the composition and mechanics of the local ECM — a phenomenon referred to as ventricular remodeling. This process impairs cardiomyocyte contractility and thus contributes to the progressive deterioration of heart function ^5^. The active cardiomyocyte response to ventricular remodeling is considered as a compensatory mechanism that aims to minimize the detrimental effects of the increased biomechanical stress on cardiac output, while protecting the integrity and the pumping activity of the organ ^59^.

Here we performed an *in silico* analysis of published expression profiling array datasets obtained from ischemic and non-ischemic heart, with the aim of identifying molecular mediators that help ensure cell adaptation to the new demands of the pathological organ by reshaping gene expression. We observed a general upregulation of RBP expression in the pathological heart in humans and in mice, independently of the etiology of the pathology. We identified hnRNPC among the RBPs consistently altered in the diseased heart; the upregulation of this abundant nuclear protein, which has so far mainly been studied for its role as a splicing factor ^28–30^, has been independently confirmed by single cell RNA sequencing in cardiomyocytes ^34^. The upregulation is more pronounced at the protein level, suggesting post-transcriptional processes might be at play in order to increase protein stability.

We took hnRNPC forward for further analyses – employing quantitative proteomics, super-resolution microscopy and PLA to disclose that hnRNPC shuttles out of the nucleus and localizes to the sarcomeres of cardiomyocytes in the pathological heart. We confirmed this observation by biochemical analyses, and found that the protein settles at the Z-disc, where it physically interacts with components of the contractile apparatus, including Troponin T2, PDLIM5, FHL2 and MYH7.

hnRNPC upregulation has been associated with different pathologies, including atherosclerosis and pre-atherosclerotic intimal hyperplasia ^33^, neurodegenerative diseases ^31^ and some forms of cancer (e.g. gastric cancer) ^60^; however, modifications in its localized function have not been described in pathological conditions. Indeed, most hnRNPs shuttle between the nucleus and the cytoplasm; hence their interaction with specific binding partners can be considered as an indicator of their function at specific cell compartments. Interestingly, in chemoresistant gastric cancer, hnRNPC was found upregulated and localized to the cytoplasm and membrane, but its function in these compartments was not questioned ^60^. Our mass spectrometry analysis of hnRNPC-interacting proteins in diseased human heart samples confirmed that hnRNPC protein interacts with the spliceosome, as previously demonstrated ^61^. However, the presence of spliceosome components was reduced in hnRNPC interactome in failing hearts compared to healthy hearts. Interestingly, the presence of proteins belonging to the cytoskeleton, the sarcomere and the ribosome specifically in the failing heart interactome suggested a different function for hnRNPC in diseased conditions.

To determine what this new function might be, we performed RIP-seq and identified several specific exon-enriched hnRNPC-bound RNAs in the failing heart, mostly encoding for sarcomeric proteins. Some of these mRNAs, including titin (*TTN*), cardiac muscle alpha actin (*ACTC1*) and cardiac troponin I3 (*TNNI3*) are translated locally at the sarcomeric site ^42,43^. Together with the evidence that hnRNPC interacts with ribosomal proteins at the sarcomere, these data led us to hypothesize that hnRNPC shuttling to the contractile apparatus in the failing heart could be associated with the localized translation of transcripts encoding for contractile proteins.

Localized translation has been described in different cell types, including neurons ^62^, fibroblasts ^63^ and myoblasts ^64^, as a strategy aimed at maximizing the synthesis of needed proteins directly at the site of exploitation. Indeed, hnRNPC was previously found in RNA granules in neurons together with mRNAs involved in synaptic remodeling, which are transported to the subsynaptic site and translated in response to an input ^65^. Sarcomeres serve as sites of active translation in the rat heart, where the translation machinery and mRNAs encoding for sarcomeric proteins are distributed in a cross-striated pattern ^43^. We used iPSC-derived cardiomyocytes to visualize the active incorporation of PMY (an indicator of active translation) at the sarcomere, where ribosome components are found. By coupling decellularized extracellular matrices obtained from healthy and diseased human hearts with ribopuromycilation and PLA, we found that the pathological ECM induces hnRNPC recruitment to the sarcomeric active sites of translation.

Because cardiac ECM remodeling is associated with mechanical stress, we used 3D hydrogels with controlled stiffness and micropatterned surfaces to demonstrate that a fraction of hnRNPC can shuttle outside of the nucleus when the transmission of intracellular tension is hindered. We confirmed that its shuttling to the cytoplasm is precipitated in constrained cells not able to propagate intracellular tension. It thus seems that the mechanical stress associated with ECM remodeling serves as a switch for hnRNPC redistribution from the nucleus to the sarcomere, presumably to modulate its role in RNA homeostasis. Given hnRNPC shuttling out of the nucleus and its reduced interaction with spliceosomal components, we investigated the effect of hnRNPC depletion on pre-mRNAs AS. To mimic a situation in which hnRNPC function in splicing is reduced, we induced its shuttling from the nucleus by pharmacological inhibitors of tension in NHDFs, which are known to be extremely sensitive to mechanical cues. Together with the results obtained by hnRNPC knock down in the same cell type and iPSC-derived cardiomyocytes, these experiments indicate hnRNPC depletion from the nucleus affects the AS of mechanosensitive transcripts and genes involved in cardiovascular diseases. Next, we used iCLIP-seq to confirm the presence among hnRNPC direct targets of components of the Hippo pathway, a molecular axis involved in organogenesis and whose dysregulation has been associated to numerous pathologies ^66^, including those affecting the heart ^22,52,55,67^. In particular, we found that the mechanically induced hnRNPC displacement from the cell nucleus or its depletion affect the inclusion of YAP1 exon 4. This splicing event is of particular relevance, since the exon encodes for the second WW domain, a domain known to directly control the protein interaction with TEAD transcription factor and thus its DNA binding activity ^49^. This switch towards the expression of YAP1 isoforms with two WW domains (YAP1-2) has been recently described by our research group in human pathological hearts^56^ and confirmed in other two independently published RNA-seq datasets.

Going forward, questions remain as to which upstream factors are involved in hnRNPC regulation in failing hearts and what is the specific role of hnRNPC at the translation sites (repressor/activator).

Indeed, although our data suggest an involvement of hnRNPC in localized translation at the sarcomeric site in HF, future work is needed to elucidate whether hnRNPC plays a direct role in this process and its relevance in the establishment and progression of cardiac diseases.

The presence of the hnRNPC transcript in our RIP- and iCLIP-seq analyses suggests that hnRNPC could regulate its own RNA post-transcriptionally. RBPs were previously proposed to bind and regulate the fate of their transcripts in an autoregulatory feedback ^68^. Further studies are thus needed to elucidate whether hnRNPC adopts this mechanism to regulate its expression and function in cardiac pathologies.

In conclusion, our results on the regulation of hnRNPC localization and function unveil a novel mechanism by which RNA homeostasis could undergo mechanical regulation following pathological cardiac ECM remodeling (**Fig. 9**). Future experiments that study the mechanical control of RNA homeostasis and the players involved are now warranted, as the ability of RNA binding proteins (RBPs) to serve as possible mechanosensors could explain how ECM mechanics can directly affect cell differentiation and function by controlling RNA fate in a stage-specific manner.

**Figure 9:**
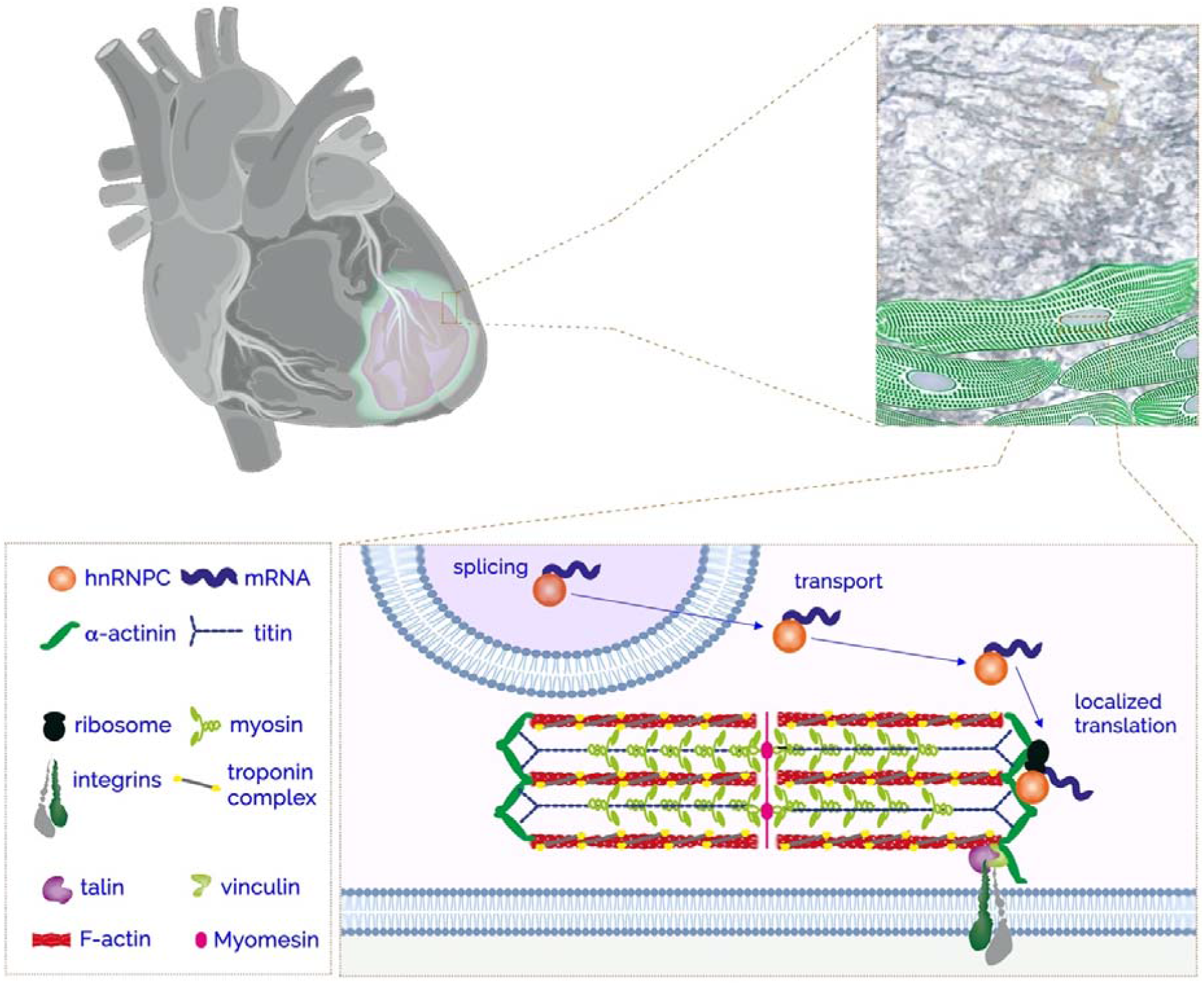
Graphical representation of hnRNPC displacement in the pathological heart.

## Methods

### Study approval

This study was performed in accordance with the ethical standards of the Centre of Cardiovascular and Transplantation Surgery and was approved by the Ethics Committee of St. Anne’s University Hospital, Brno, Czech Republic. The experiments were done according to the Declaration of Helsinki (2000) of the World Medical Organization. Experiments involving mice were approved by the Instituto de Biologia Molecular e Celular – Instituto de Engenharia Biomédica (IBMC-INEB) Animal Ethics Committee and the National Direção Geral de Veterinária (permit no: 022793), and conformed with Directive 2010/63/EU of the European Parliament.

### Patient-derived heart samples

Diseased heart tissue samples (N=17) were obtained from left ventricle of patients diagnosed with end-stage heart failure undergoing cardiac transplantation or ventricular assist device implantation. The classification of ischemic (N = 8) and non-ischemic (N = 9) heart failure is defined based on the patient anamnesis, as per the International Statistical Classification of Diseases and Related Health Problems (ICD-10). Healthy hearts (N=3) obtained from deceased organ donors without a history of cardiac disease were used as a control (**Supplementary Table 1**).

### Mouse model of myocardial infarction (MI)

MI was experimentally induced by ligating the left anterior descending coronary artery in C57BL/6 adult mice (9-10 weeks), as previously described ^36^ (**Supplementary Methods**). Mice hearts cohort was composed of 4 days post-MI (acute MI), 21 days post-MI (chronic MI), sham-operated (no ligation of the coronary artery) (**Supplementary Fig. 3b**). All animal experiments were approved by the local Ethics Committees.

### Proximity Ligation Assay

The proximity ligation assay (PLA) was performed on fixed (4% PFA) heart tissue slices and cells using DuoLink PLA technology probes and reagents (**Supplementary Table 2**) and following the manufacturer’s protocol (**Supplementary Methods**).

### Ribopuromycilation

Ribopuromycilation protocol (RPM) to visualize active sites of translation was performed following the original RPM protocol – Procedure A^66,67^ (**Supplementary Methods; Supplementary Fig. 5d**). The RPM experiments were performed on beating cardiomyocytes between days 20 and 30 of differentiation.

### Cell culture, differentiation, transfection, treatment and micropatterning

Culture of the human induced pluripotent stem cell line DF 19-9-7T (iPSC) (WiCell, Madison, WI, USA) and normal human dermal fibroblasts (NHDF, ATCC) as well as cardiac differentiation is detailed in **Supplementary Methods**. Cardiac troponin T (**Supplementary Table 2**) was used as sarcomeric marker of iPSC-derived cardiomyocytes. Treatment of cells with inhibitors of cytoskeletal tension, 3D cultures in fibrin gels and micropatterning are detailed in **Supplementary Methods**.

*HNRNPC* knockdown was performed as previously described^27^. Detailed procedures for siRNA transfection, qRT-PCR and western blotting are described in **Supplementary Methods**.

### Protein Immunoprecipitation (IP)

For the IP of hnRNPC-binding proteins, 100 mg of tissue from the human heart apex was used for each IP and control sample (**Fig. 2a**). The sample was washed in ice-cold PBS, dissected and mechanically homogenized in 1ml of Lysis Buffer using 3.0 mm Zirkonia beads and Beadbug homogenizer. Dynabeads Protein G (50 µl/sample) were incubated with 200 µl primary antibody (5 µg/sample) for 45 min at 4 °C. Heart lysates were incubated with antibody conjugated Dynabeads overnight at 4°C. The proteins were then eluted by incubation with 100 µl 8M Urea on a shaker (20 min at RT). The eluates were then processed for mass spectrometry analysis (**Supplementary Methods; Supplementary Table 2**).

### Mass spectrometry

After protein immunoprecipitation, the samples were digested with trypsin using the FASP method. The digested peptides were labeled using TMT pro16-plex isobaric mass tagging reagents (Thermo Scientific) and the fractions were analyzed on an Orbitrap Fusion™ Lumos™ Tribrid™ mass spectrometer interfaced with an Easy-nLC1200 liquid chromatography system (Thermo Fisher Scientific). The IP/IgG abundances were calculated for each sample. A protein was considered successfully immunoprecipitated when a ratio > 1.5 was detected in both biological replicates. See also **Supplementary Methods** and **Supplementary Table 2**.

### RNA Immunoprecipitation (RIP)

To immunoprecipitate hnRNPC-binding RNAs, 100 mg tissue from the human heart apex was used for each IP and control sample. In brief, the heart samples were dissected, cross-linked in 1% (v/v) formaldehyde, mechanically dissociated in 1 ml Polysome Lysis Buffer (PLB) using 3.0 mm Zirkonia beads and a Beadbug homogenizer and sonicated. Dynabeads Protein G (100 µl/sample) were incubated with 10 µg of primary antibody (sc-32308, sc-2025) for 1 h at 4 °C. Cross-linked lysate was incubated with antibody-conjugated Dynabeads overnight at 4°C. Reverse cross-linking was performed and the proteins were digested by incubation with Proteinase K (60 µg/sample) for 1.5 h at 55 °C on a shaker. RNA extraction was then performed following standard phenol:chloroform phase separation. The protocol is depicted in **Fig. 4a** and detailed in **Supplementary Methods**.

### RNA sequencing, RIP-seq and alternative splicing (AS) analysis

A sequencing library was prepared using a NEBNext Ultra II Directional Kit (New England Biolabs, MA, USA). For the RIP-sequencing, the immunoprecipitated RNA was used as an input into the total RNA library preparation protocol; control input RNA was first rRNA depleted using QIAseq Fastselect HMR Kit (Qiagen, Germany). For the alternative splicing analysis, 200-300 ng total RNA was used as an input into the polyA enrichment module protocol. Samples were fragmented and transcribed into cDNA. Following universal adapter ligation, the samples were barcoded using dual indexing primers. The RNA resultant from RIP of heart tissue or resultant from NHDF transfection with hnRNPC siRNA and the respective control construct was sequenced on an Illumina Nextseq 550 sequencer (Illumina, CA, USA) to 25-35 million single-end 75bp reads and >50 million paired-end 75 bp reads per sample, respectively. Detailed sequencing procedure and analysis are available in the expanded methods section (**Supplementary Methods; Supplementary Table 2**).

### Individual-nucleotide resolution UV crosslinking and immunoprecipitation (iCLIP) of hnRNPC1/C2

NHDFs were grown to 85-90% confluency and exposed to 150 mJ/cm2 at 254 nm in a Stratalinker 2400 on ice. Cell pellets were resuspended in 1ml of ice-cold lysis buffer (**Supplementary Methods**). The insoluble fraction was removed by centrifugation at 20.000g for 20 min at 4°C and the supernatant was used for the iCLIP experiment. Dynabeads Protein G (Invitrogen) (100 µl/sample) were incubated with 15 µg of anti HNRNPC1/C2 antibody (Santa Cruz) in a total volume of 100 µl of lysis buffer for 1 h at RT. Antibody coupled beads were mixed with 1ml of soluble cell lysate and incubated for 1hour on a rotator at 4°C. Dephosphorylation, radiolabeling (γ-ATP) and 3’ end adapter ligation was carried out on the protein-RNA complex. The resultant Protein-RNA complex was resolved on a gradient NuPAGE gel and blotted onto nitrocellulose membrane. Post-autoradiography the membrane was cut between 48-100kDa size and Proteinase K treatment was performed to isolate RNA. The cDNA libraries were prepared as previously described ^69^ with slight modifications. The purified library was analyzed by 75bp single end high-throughput sequencing at Illumina HiSeq2500. Protocol and reagents are detailed in **Supplementary Methods and Supplementary Table 2**. A schematic representation of the protocol can be found in **Supplementary Fig. 7f**.

### Decellularization of human myocardial tissue and preparation of ECM coating

The decellularization protocol was performed as previously described^23,70^ (**Supplementary Methods**). The ECM-enriched scaffold was mechanically disrupted and used as 2D coating for cell culture (**Fig. 5c, d**).

### Microscopy and image analysis

Histology, immunohistochemistry and immunocytochemistry are detailed in **Supplementary Methods**. A Zeiss LSM 780 confocal microscope with 40x (1.3 NA) oil-immersion or 20x air (0.8 NA) objective lenses was used for image acquisition. ImageJ and Imaris software were used for image quantification (**Supplementary Methods**). For the Masson’s Trichrome analysis, a slide scanner Zeiss Axio Scan Z1 microscope was used to visualize whole tissues using the bright-field mode and a 10x objective. Super-resolution imaging was performed using a 3D-SIM Deltavision OMX with a 60x objective (1.42 NA).

### Statistical Analyses

All data are expressed as the mean ± standard deviation (S.D.) unless otherwise specified. The statistical analyses were performed by using GraphPad Prism v. 6.0 software package. The null hypothesis was rejected when P < 0.05. The number of biological (N) and technical (n) replicates and the statistical tests used are indicated in figure legends and in the Source Data file.

## Supporting information

Supplementary dats legends

## Data Availability

All data supporting the findings of this study are available as Supplementary Data. Sequencing data have been deposited to GEO database under the reference series GSE155474 and GSE169068. The Mass Spectrometry proteomics data have been deposited to the ProteomeXchange Consortium via the PRIDE partner repository under the dataset identifier PXD020663.

## Acknowledgements

The authors would like to thank Carina Sihlbom and Johannes Fuchs at the Proteomics Core Facility at Sahlgrenska Academy, University of Gothenburg, Sweden for proteomic analysis. They also thank the technical support team at the Center of Translational Medicine, the members of the Mass Spectrometry Core Facility of FNUSA-ICRC, the Core Facility Bioinformatics of CEITEC Masaryk University, the CF Genomics supported by the NCLG research infrastructure (LM2018132 funded by MEYS CR), for their assistance with obtaining the scientific data presented in this paper. The authors are grateful to Jessica Tamanini of Insight Editing London for reviewing and editing the manuscript prior to submission and to Jan Vrbsky for primer design. The authors also thank Perpetua Pinto-do-Ó and Diana S Nascimento for providing the mouse model of MI. Super-resolution microscopy was performed at the Materials Characterization and Fabrication Platform (MCFP) at The University of Melbourne.

## Author contributions

FM: Conceptualization, Experimental Design and Execution; ARP: Experimental Execution; VHE: Bioinformatics analysis; NMV, SV: iCLIP-sequencing; HD: Experimental Execution; VHO: human tissue harvesting and patient anamnesis; FC, FCAR: Superresolution microscopy, manuscript drafting; WSA, APG: RNA-IP protocol setting and data interpretation; MAOC: paper conceptualization and drafting; SP, GF: project supervision, paper conceptualization and drafting.

