## Supplementary dats legends for "The mechanical regulation of RNA binding protein hnRNPC in the failing heart"

**Supplementary Data 1**. Datasets analysed in Figure 1a, b.

- Datasets details

- List of upregulated genes in dataset1 (mouse MI); dataset2 (mouse HF); datasets 3 and 4 (human HF)

**Supplementary Data 2**

List of hnRNPC interactors found by MS analysis in HF samples

List of hnRNPC interactors found by MS analysis in healthy samples

Proteomaps (based on the KEGG database) representing the quantitative composition of the hnRNPC interactome in healthy and HF human hearts

**Supplementary Data 3**

List of FHL2 interactors found by MS analysis in HF samples

List of PDLIM5 interactors found by MS analysis in HF samples

List of the common binding partners of hnRNPC, FHL2 and PDLIM5 in HF samples

GO cellular component analysis of the common interactors of hnRNPC, FHL2 and PDLIM5 in HF samples

**Supplementary Data 4**

List of transcripts bound by hnRNPC in HF samples found by RNA-IP sequencing.

List of exon-enriched targets and intron-enriched targets

**Supplementary Data 5**

List of transcripts undergoing AS (SE; MXE; A5SS; A3SS; IR) in *HNRNPC* KD cells and pathway enrichment analysis by WikiPathways.

**Supplementary Data 6**

1) **Hippo_ASanalysis.** Transcripts belonging to the Hippo pathway displaying alternative splicing in hnRNPC KD cells. KEGG mapper representation of HIPPO pathway: transcripts showing altered AS in hnRNPC KD cells are highlighted in red
2) **Hippo_iCLIPanalysis.** Transcripts belonging to the Hippo pathway bound by hnRNPC in NHDF cells. KEGG mapper representation of HIPPO pathway: hnRNPC targets are highlighted in red
3) **Hippo_AS_iCLIP_Venn.** Venn diagram showing the transcripts belonging to Hippo pathway common to the two datasets
4) **Hippo_RNAIP.** Transcripts belonging to the Hippo pathway bound by hnRNPC in failing hearts. KEGG mapper representation of HIPPO pathway: hnRNPC targets are highlighted in red
5) **Hippo_AS_iCLIP_RNAIP.** Venn diagram showing the transcripts belonging to Hippo pathway common to the three datasets

**Supplementary Data 7**

Number of reads in each sample (N=3);

List of hnRNPC targets;

List of transcripts common to iCLIP analysis and AS analysis;

Number of reads on coding vs non-coding elements;

Number of reads on Alu elements.

**Supplementary Data 8**

List of transcripts common to iCLIP, RNA-IP and AS analyses

List of disease categories found by CTD analyses of the transcripts common to iCLIP, RNA-IP and AS analyses

List of hnRNPC-targets belonging to the cardiovascular disease category
