## Supplementary Materials for "The mechanical regulation of RNA binding protein hnRNPC in the failing heart"

**Supplementary Information**

**Supplementary Figures:**

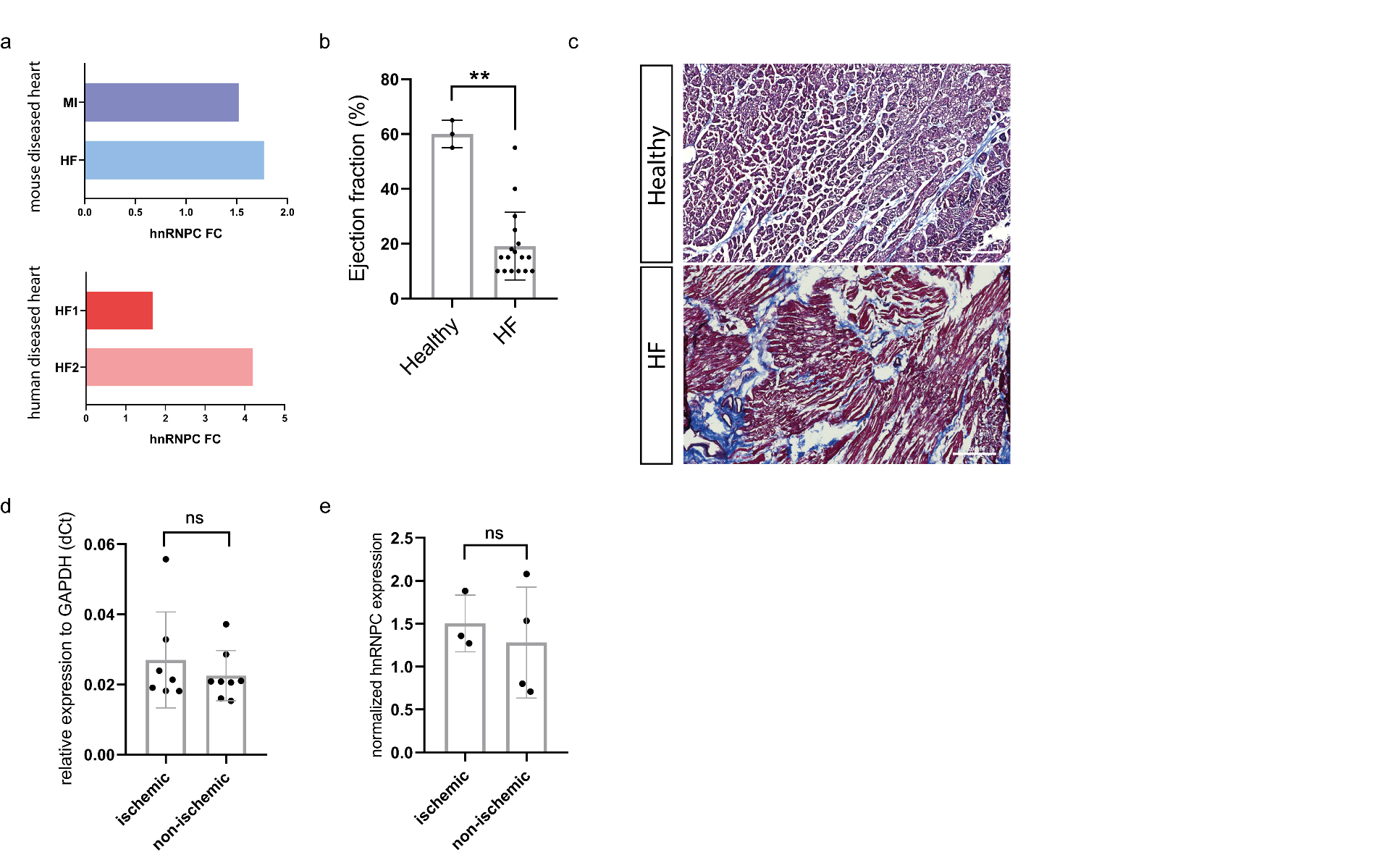
**Supplementary Figure 1**. **a)** Top: Barplot representation of hnRNPC expression in mouse diseased heart datasets. Bottom: hnRNPC expression in human diseased heart failure (HF) datasets. The data are expressed as fold change (FC) as compared to their respective control. **b)** Ejection fraction representation of the patients enrolled in the study. Data are presented as mean ± S.D.; **p < 0.01, Mann-Whitney test. Healthy (N = 3); heart failure (HF, N = 17). **c)** Representative Masson’s Trichrome staining of human heart tissue. Collagen, blue; muscle fibers, red. Scale bar = 500 μm. **d)** Dotplot representation of hnRNPC RNA expression in ischemic HF (N = 7; n = 2) compared to non-ischemic heart tissues (N = 8; n = 2) as obtained by RT-qPCR analysis. **e)** Dotplot representation of hnRNPC protein expression in samples of ischemic (N = 3; n = 3) and non-ischemic (N = 4; n = 3) heart tissues as obtained by western blot (see also Fig. 1d). GAPDH was used for total protein loading normalization.

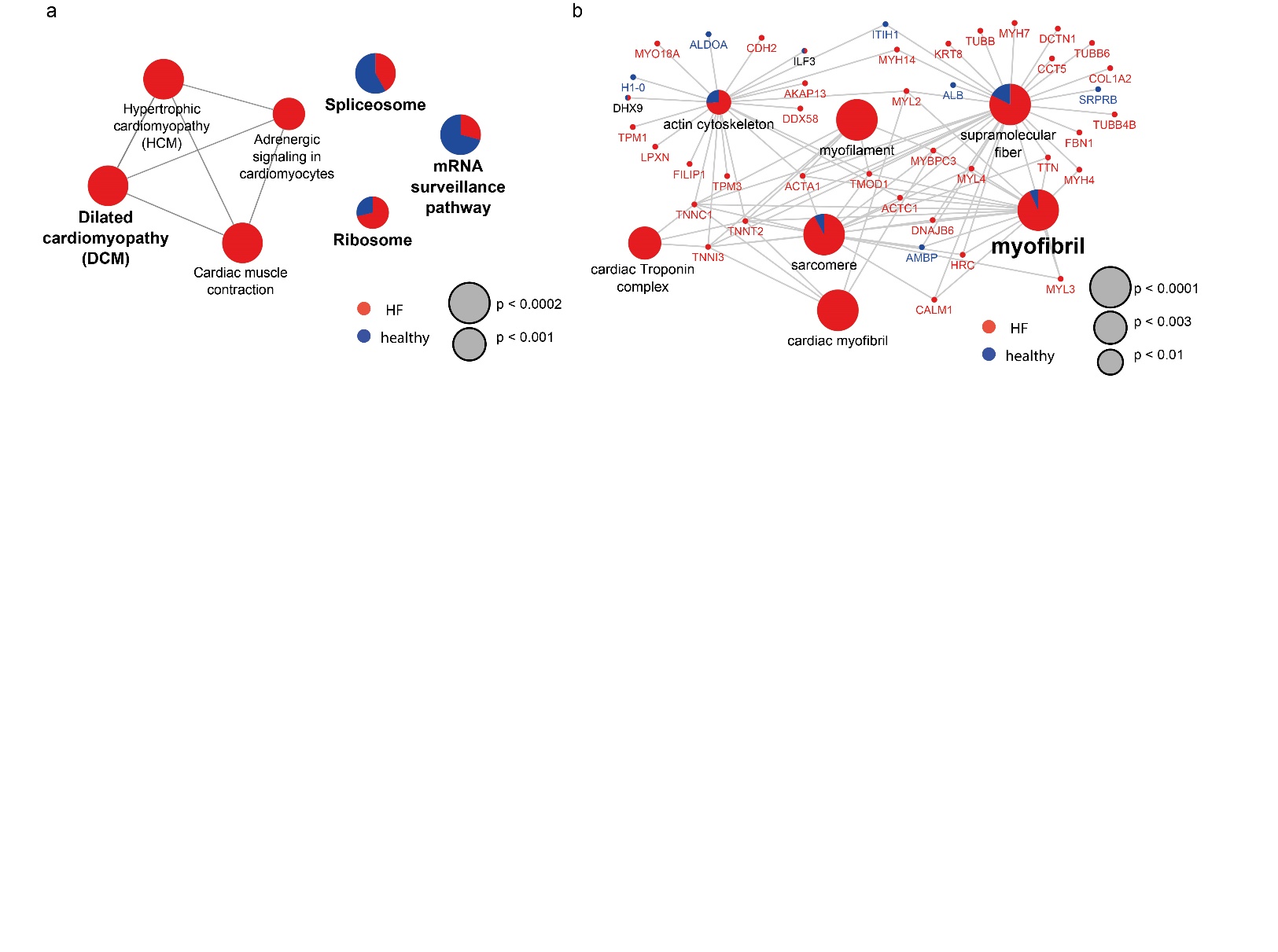

**Supplementary Figure 2.** **a)** Network of the pathways found enriched in failing (N = 2) (red) or healthy (N = 2) (blue) hearts based on KEGG database. The size of the nodes indicate the Bonferroni adjusted Term p-values. **b)** Gene-ontology (GO) network representation of the myofibril cluster found significantly [Group p-value (corrected with Bonferroni): 1.31E-08] enriched in failing (HF, blue) heart tissue. The proteins belonging to each node and found in hnRNPC-IP are depicted. The size of the nodes indicate the Bonferroni adjusted Term p-values.

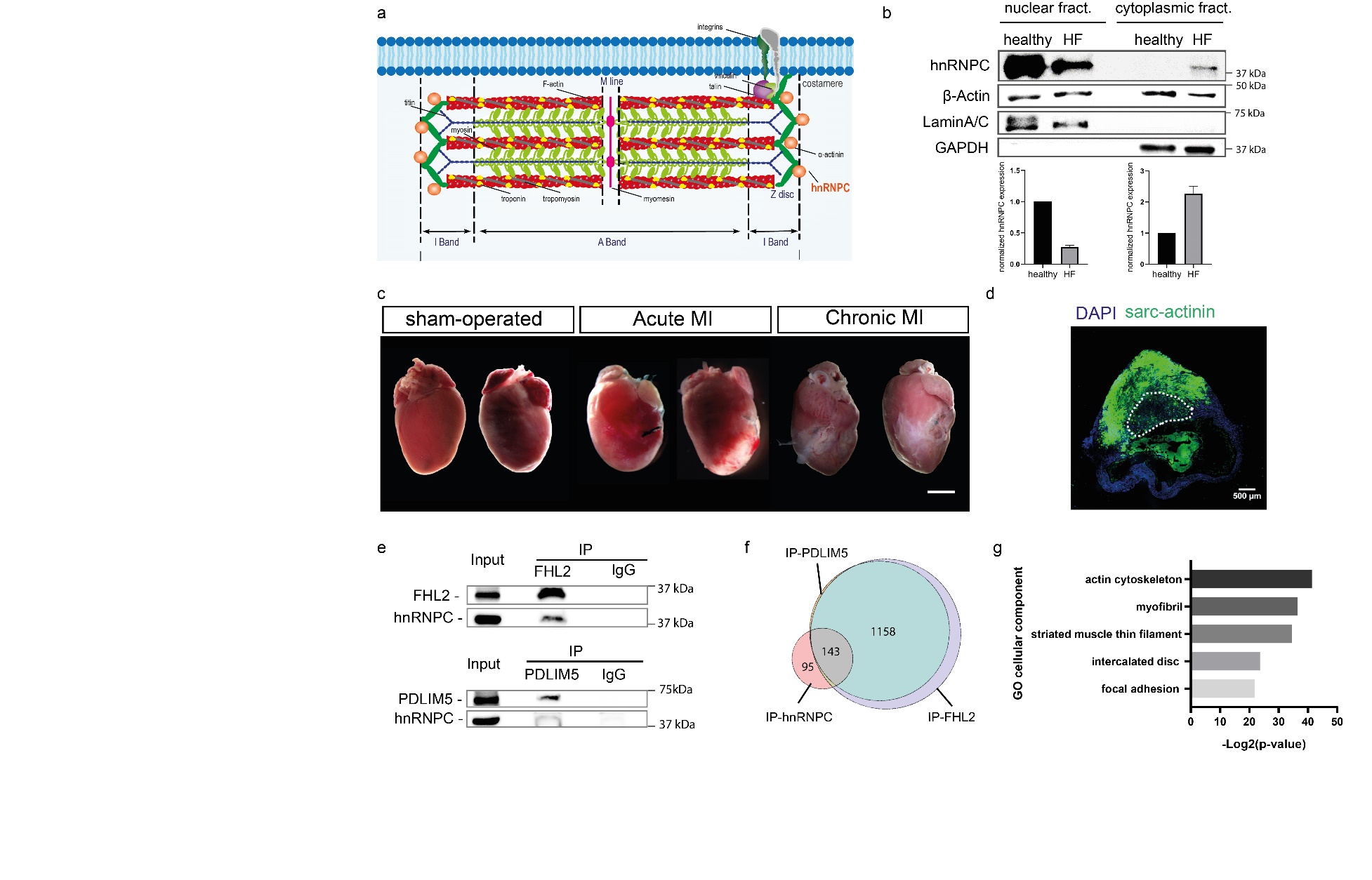

**Supplementary Figure 3.** **a)** Schematic representation of the sarcomere structure and hnRNPC distribution (orange beads). **b)** Western blot analysis (top) and quantification (bottom) of hnRNPC expression in nuclear and cytoplasmic fractions of healthy and diseased (HF) human hearts. B-actin was used for loading normalization. Lamin A/C and GAPDH were used to validate the fractionation protocol. Band intensities corresponding to hnRNPC in the nuclear and cytoplasmic fractions were quantified using Bio-Rad Image Lab software. Data are presented as mean ± S.D. (N = 2). **c)** Representative images of hearts collected from a mouse model of myocardial infarction (MI) 4 days post-surgery (Acute MI) or 21 days post-surgery (Chronic MI) and sham-operated mice. Scale bar = 200 mm. **d)** Representative confocal image of a transverse mouse heart section 21 days post-MI (Chronic MI). The frozen section was stained for sarcomeric actinin (sarc-actinin, green) and counterstained with DAPI. The dashed line indicates the infarct border zone. Scale bar = 500 μm. **e)** Western blot analysis of hnRNPC co-immunoprecipitation with FHL2 and PDLIM5 in diseased human heart (total lysate), FHL2- and PDLIM5- immunoprecipitated samples and negative control (IgG). **f)** Venn diagram representation of the proteins found interacting with hnRNPC, PDLIM5 and FHL2 in immunoprecipitated samples in failing human hearts. **g)** Barplot representation of cellular components Gene Ontology (GO) categories significantly enriched in the binding partners common to hnRNPC, PDLIM5 and hnRNPC in the failing human heart.

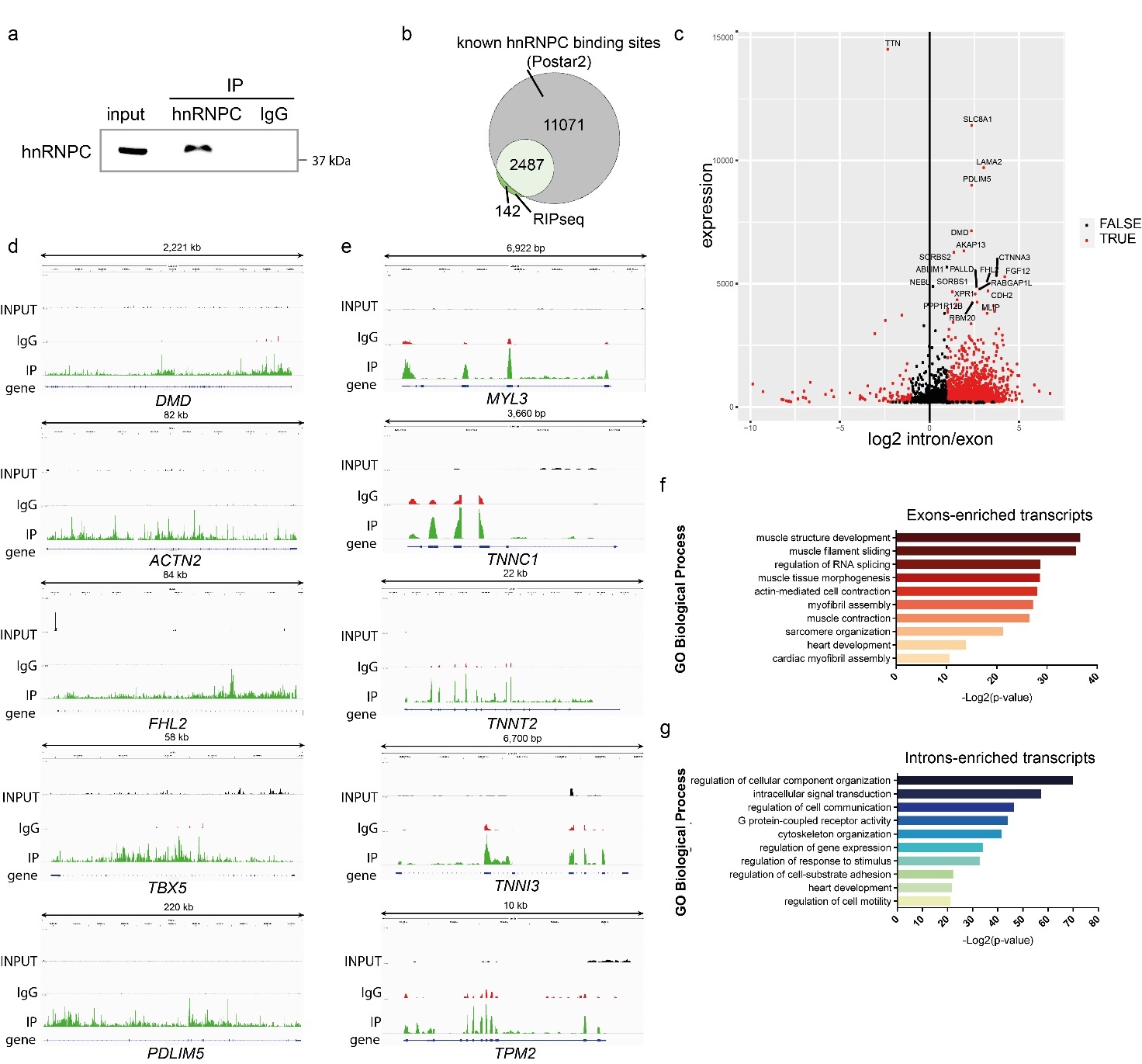

**Supplementary Figure 4**. **a)** Western blot analysis of hnRNPC protein in the total human failing heart lysate (input), the hnRNPC immunoprecipitated sample and the negative control (IgG) following a RIP-seq protocol. **b)** Venn diagram representing the transcripts enriched in hnRNPC-immunoprecipitated samples over the input (green) and the transcripts harboring at least one binding site for hnRNPC, collected from the POSTAR2 database (grey). **c)** Volcano plot representing the transcripts found enriched in hnRNPC immunoprecipitated samples over the input. The x-axis shows the intron-exon ratio on a logarithmic scale: log_2_ (intron/exon) < -1 indicates exons-enriched transcripts whereas log_2_ (intron/exon) > 1 indicates introns-enriched transcripts. **d)** Individual gene representation of representative intron-enriched hnRNPC targets. Read coverage is displayed in INPUT (black), IgG (red) and IP (green) samples. **e)** Individual gene representation of representative exon-enriched hnRNPC targets. Read coverage is displayed in INPUT (black), IgG (red) and IP (green) samples. **f)** Gene Ontology (GO) of the biological processes of interest found significantly enriched (p value < 0.01) in exons-enriched RNAs bound to hnRNPC in failing heart tissue. **g)** GO of biological processes of interest found significantly enriched (p value < 0.01) in introns-enriched RNAs bound to hnRNPC in failing heart tissue.

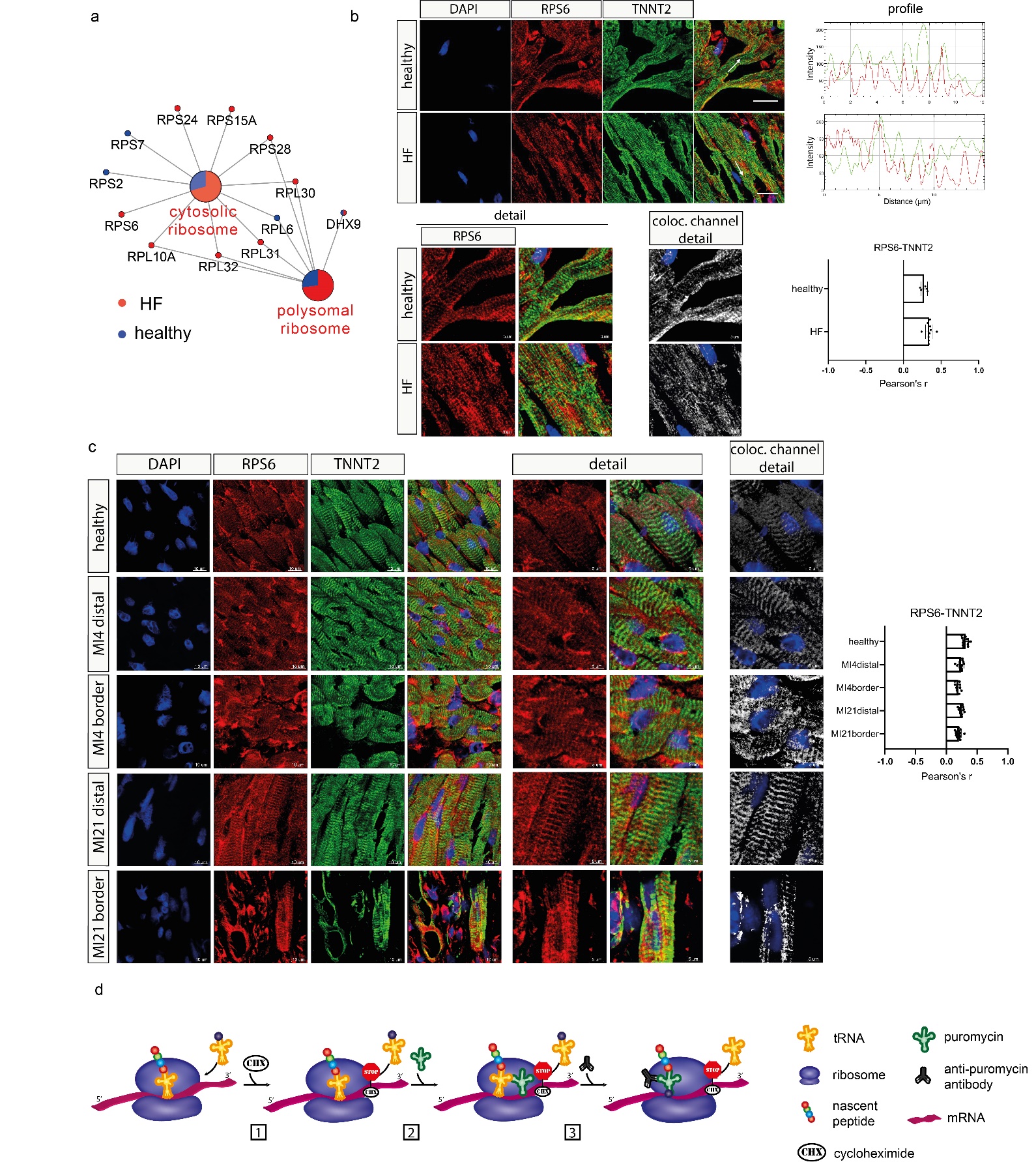

**Supplementary Figure 5**. **a)** GO network representation of the cytosolic and polysomal ribosome categories (GO cellular component) found enriched (p-value < 0.01) in hnRNPC protein interactome in healthy (blue) and failing (red) heart. **b)** Representative confocal microscopy images of ribosomal protein S6 (RPS6) (red) expression in healthy and failing (HF) human heart. Sarcomeric units are identified by cardiac troponin T (TNNT2, green) staining and nuclei are counterstained with DAPI (blue). Image analysis shows the intensity profiles of RPS6 and TNNT2 in representative cells. The superimposed white arrow indicates the region of interest. Bottom: the detail shows the colocalization (white) of red and green signals. Pearson’s coefficient was calculated by IMARIS (n≥5). Data are presented as mean ± SDs. Scale bar = 10 μm; Scale bar of detail: 5 μm **c)** Representative confocal images showing the localization and relative profile of RPS6 (red) in healthy and border or distal areas of the infarcted mouse heart (MI). Cardiomyocytes are stained with cardiac troponin T (TNNT2, green) and the nuclei are counterstained with DAPI (blue). The detail shows the colocalization (white) of red and green signals. Pearson’s coefficient was calculated by IMARIS (n≥10). Data are presented as mean ± SDs. Scale bar = 20 μm. Scale bar detail = 8 μm. **d)** Schematic of the strategy adopted to identify sites of active protein translation in iPSC-derived cardiomyocytes by means of ribopuromycilation assay: 1) elongation inhibition, 2) puromycin incorporation, 3) puromycin detection.

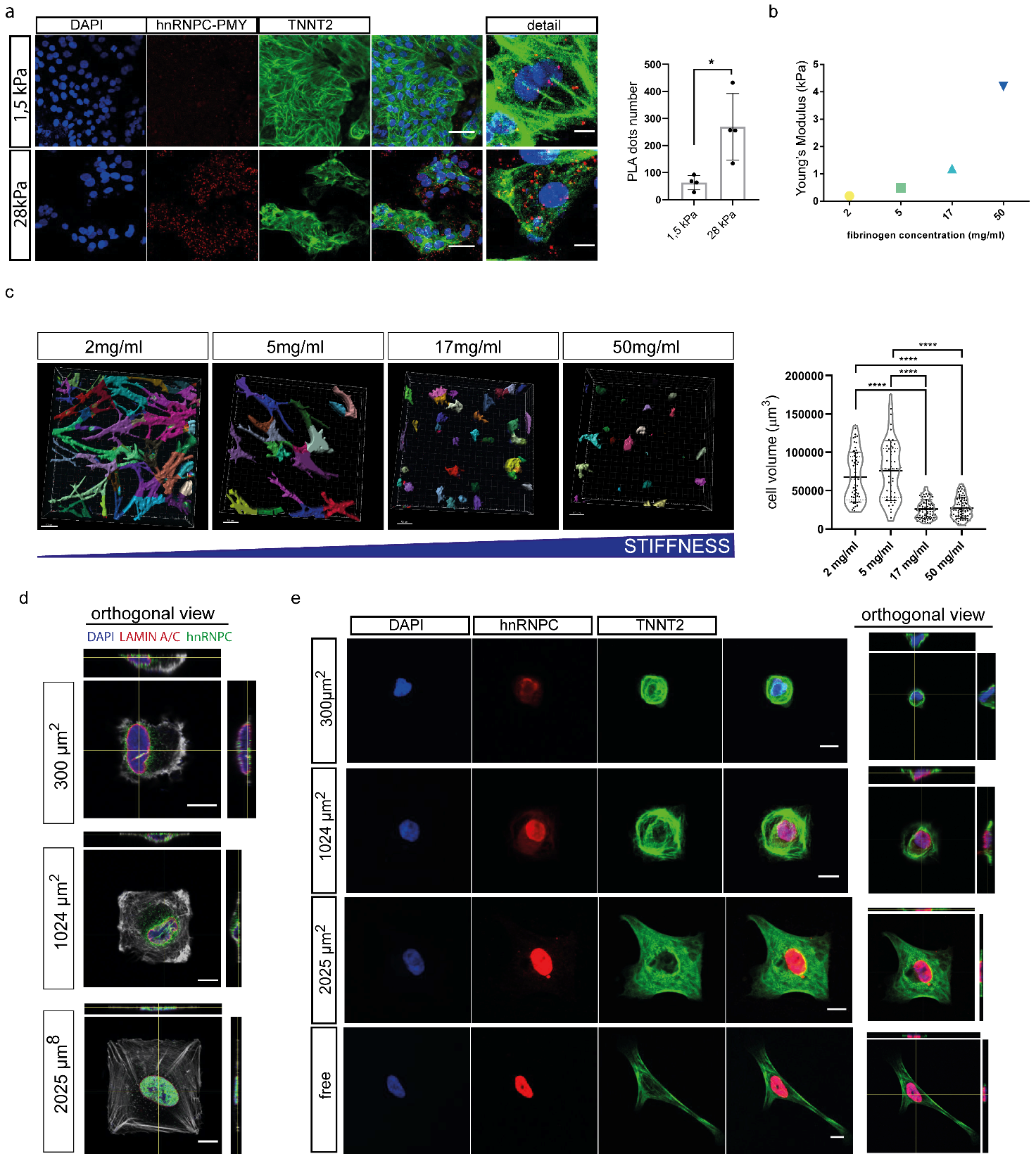

**Supplementary Figure 6**. **a)** Representative confocal microscopy images of the results obtained from PLA using antibodies against PMY and hnRNPC in iPSC-derived beating cardiomyocytes grown onto substrate with stiffness of 1.5 and 28 kPa. Cardiomyocytes are labeled for cardiac troponin T (TNNT2, green). The red signal identifies that the proteins are within 40 nm of each other. Nuclei are counterstained with DAPI (blue). Data are presented as mean ± S.D. (N =4 ; n = 5). *p < 0.05, Mann-Whitney test. Scale bar = 10 μm. **b)** The Young’s modulus (kPa) corresponding to the fibrin hydrogels used as calculated in Duong et al.,2009^1^. **c)** A 3D reconstruction (left) and cell volume measure (right) of the NHDF cells embedded in fibrin hydrogels with different stiffnesses, calculated using Imaris software. Data are presented as mean ± S.D. ( > 50 cells per condition). ****p < 0.0001; Kruskall Wallis test followed by Dunn’s multiple comparisons test. **d)** Orthogonal view of the representative confocal images showing the hnRNPC distribution in cells confined on fibronectin-coated squares with the indicated areas. The cells were stained with anti-hnRNPC (green), Alexa Fluor 647 Phalloidin, anti-LaminA/C (red) and the nuclei were counterstained with DAPI (blue). **e)** Representative confocal analysis of individual iPSC-derived cardiomyocytes grown onto 300 µm^2^, 1024 µm^2^, 2025 µm^2^ micropatterns or without patterns (free) on a fibronectin coating. The cells were stained with anti-hnRNPC (red), cardiac Troponin T (TNNT2), and the nuclei were counterstained with DAPI (blue).

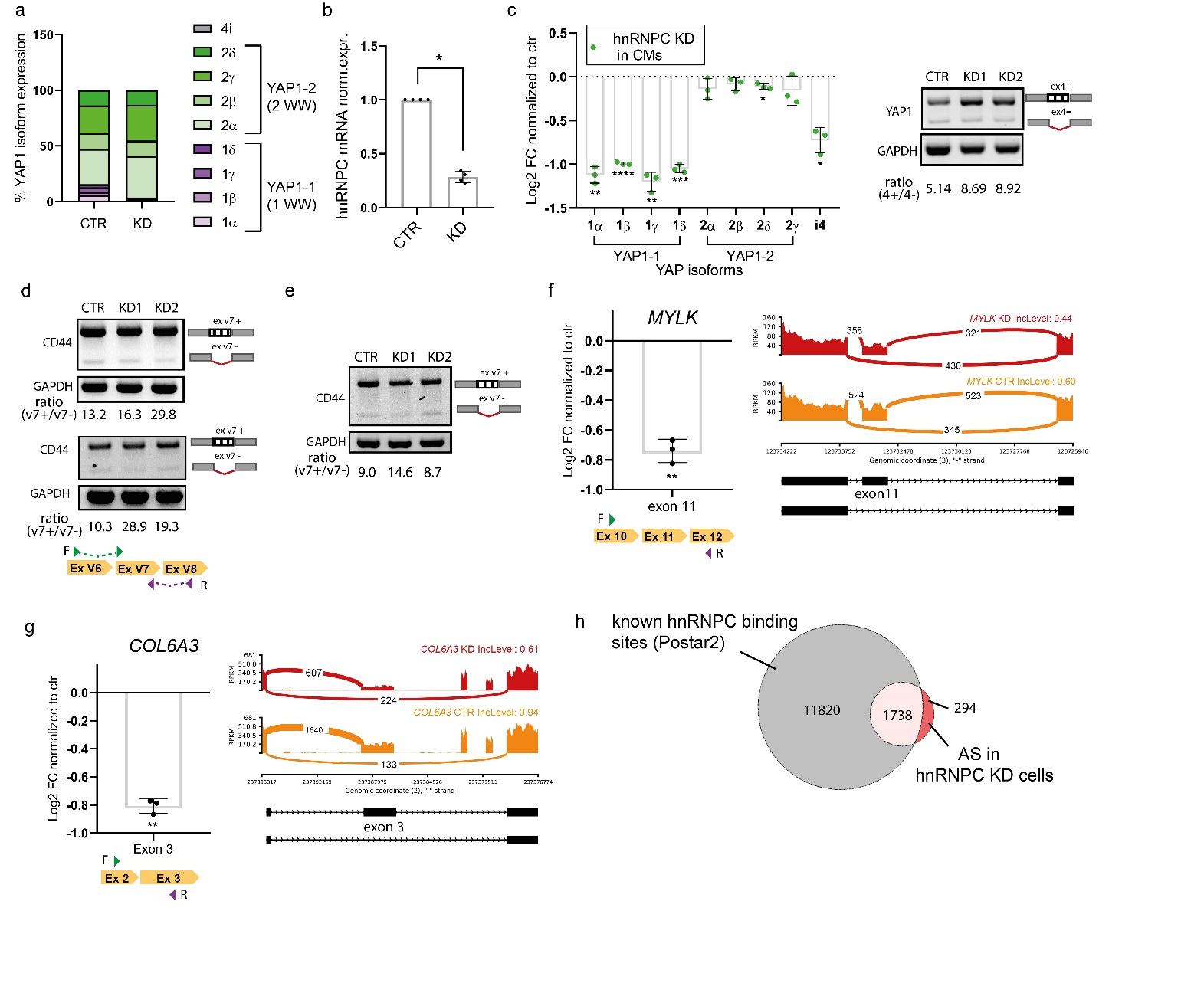

**Supplementary Figure 7**. **a)** Barplot representation of the relative expression of YAP1 splicing isoforms obtained by RT-qPCR in normal human dermal fibroblasts (NHDFs) depleted (KD) or not (CTR) for hnRNPC. The data are expressed as mean percentages of the indicated isoform ± S.D. and are normalized to GAPDH (N = 4). **b)** RT-qPCR analysis of hnRNPC mRNA expression in iPSC-derived cardiomyocytes (iPSC-CMs) 96 h after transfection with hnRNPC (KD) or control siRNAs (CTR). Data are presented as mean ± S.D. (N = 4). *p < 0.05, Mann-Whitney test. **c)** Left) Barplot representation of YAP1 isoforms mRNA expression (Log2FC) in iPSC-CMs upon hnRNPC-depletion. Data are presented as mean ± S.D. (N = 3). *p < 0.05; **p < 0.01; ***p < 0.001; ****p < 0.0001; One-sample t-test. Right) RT-PCR of YAP1 in iPSC-CMs upon hnRNPC-depletion. Top band: exon 4-included YAP1 mRNA. Bottom band: exon 4-excluded YAP1 mRNA. **d)** RT-PCR of CD44 in NHDF upon hnRNPC-depletion (Top) or upon 24h treatment with inhibitors of tension (Latrunculin and Y27632) (Bottom). Top band: variable exon 7-included CD44 mRNA. Bottom band: variable exon 7-excluded CD44 mRNA. **e)** RT-PCR of CD44 in iPSC-CMs upon hnRNPC-depletion **f)** Left) Barplot representation of MYLK exon11 expression (Log2FC) in hnRNPC-depleted NHDFs. Data are presented as mean ± S.D. and normalized to GAPDH (N = 3). **p < 0.01; One-sample t-test. Right) Sashimi plot depicting MYLK skipped exon (SE) AS event (exon 11) in hnRNPC KD cells. **g)** Left) Barplot representation of COL6A3 exon3 expression (Log2FC) in hnRNPC-depleted NHDFs. The data are presented as mean ± S.D. and normalized to GAPDH (N = 3). **p < 0.01; One-sample t-test. Right) Sashimi plot depicting representative COL6A3 skipped exon (SE) AS event (exon3) in hnRNPC KD cells. h) Venn diagram representing the genes that displayed AS events in hnRNPC KD cells (red) and the transcripts harboring at least one binding site for hnRNPC collected from POSTAR2 database (grey).

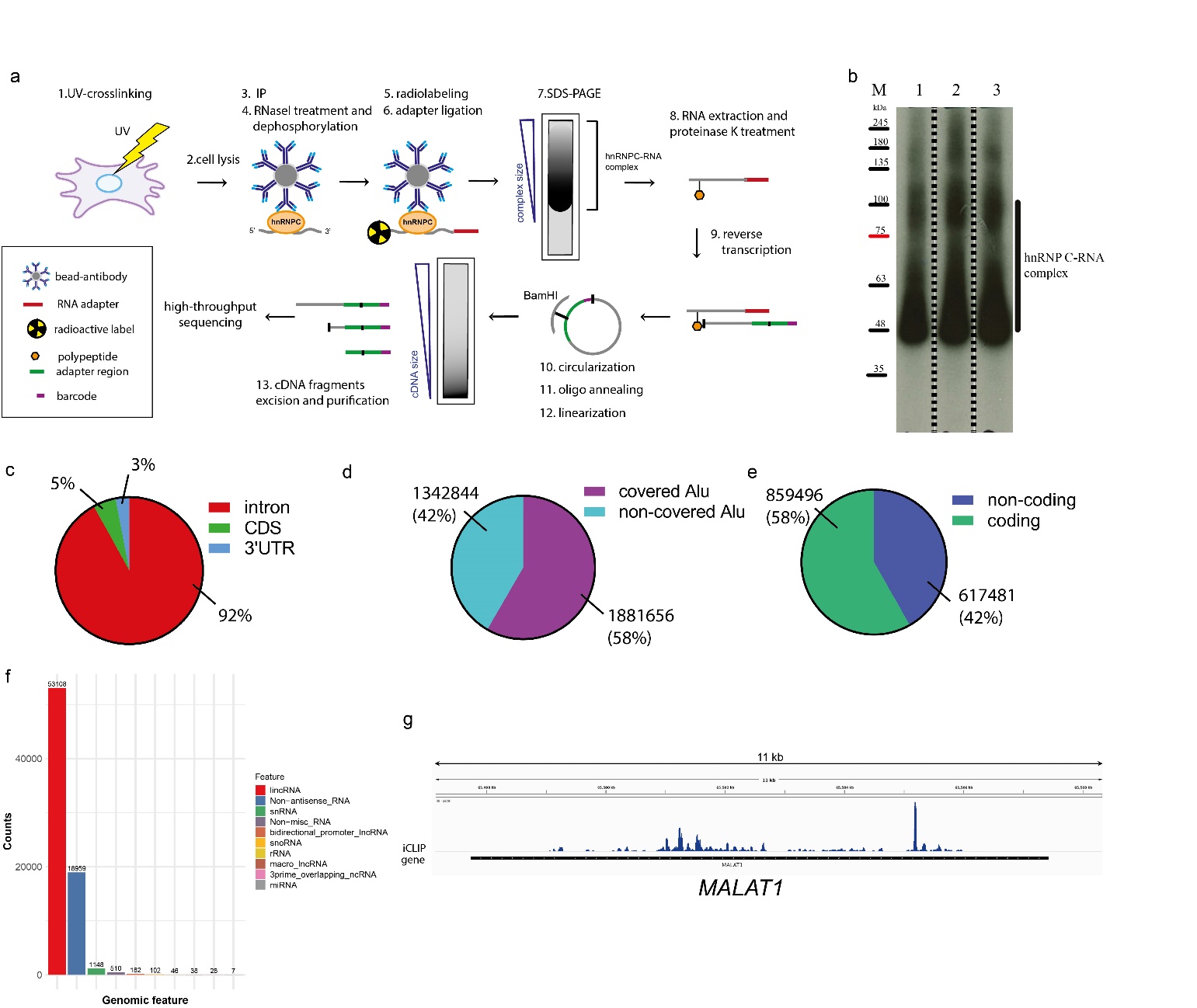

**Supplementary Figure 8.** **a)** Graphical representation of the strategy adopted to identify hnRNPC RNA targets (N=3) in NHDF cells by iCLIP followed by sequencing. **b)** Auto-radiography image of 10% IP beads used for labeling. (M) Molecular weight marker; (1)(2) and (3) Three replicates with protein RNA complex resolved on 4-12% gradient SDS PAGE. **c)** Piechart representation of the genomic distribution of the read counts showing the percentage of reads mapping on specific genomic regions (intron/exon, 5’UTR, 3’UTR, CDS) in immunoprecipitated samples (N = 3). **d)** Pie chart representation of read counts distribution on Alu elements in iCLIP samples (N = 3). **e)** Pie chart representation of read counts distribution on coding vs non-coding elements in iCLIP samples (N = 3). **f)** Bar plot representation of non-coding RNA categories bound by hnRNPC (N = 3). **g)** Individual gene representation hnRNPC binding on MALAT1: the iCLIP averaged (N = 3) is shown in blue.

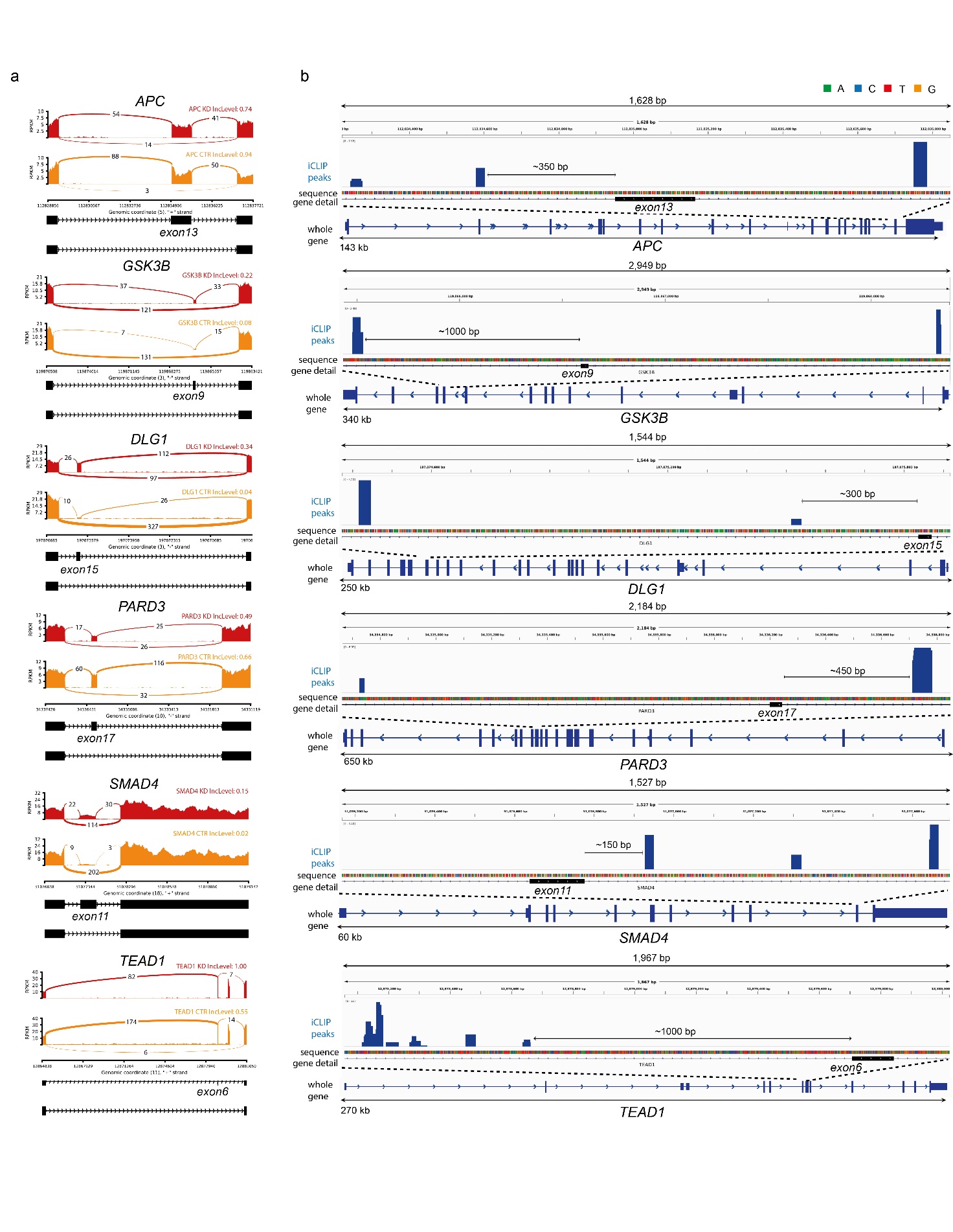

**Supplementary Figure 9.** **a)** Sashimi plots depicting representative skipped exon (SE) AS events occurring in transcripts belonging to the Hippo pathway in hnRNPC KD cells NHDFs (red) and controls (orange). The exon inclusion level (IncLevel) is indicated. **b)** Genome browser view of individual genes displaying the iCLIP data (crosslink events per nucleotide) of hnRNP C (blue) (average of N = 3) in proximity of specific exons.

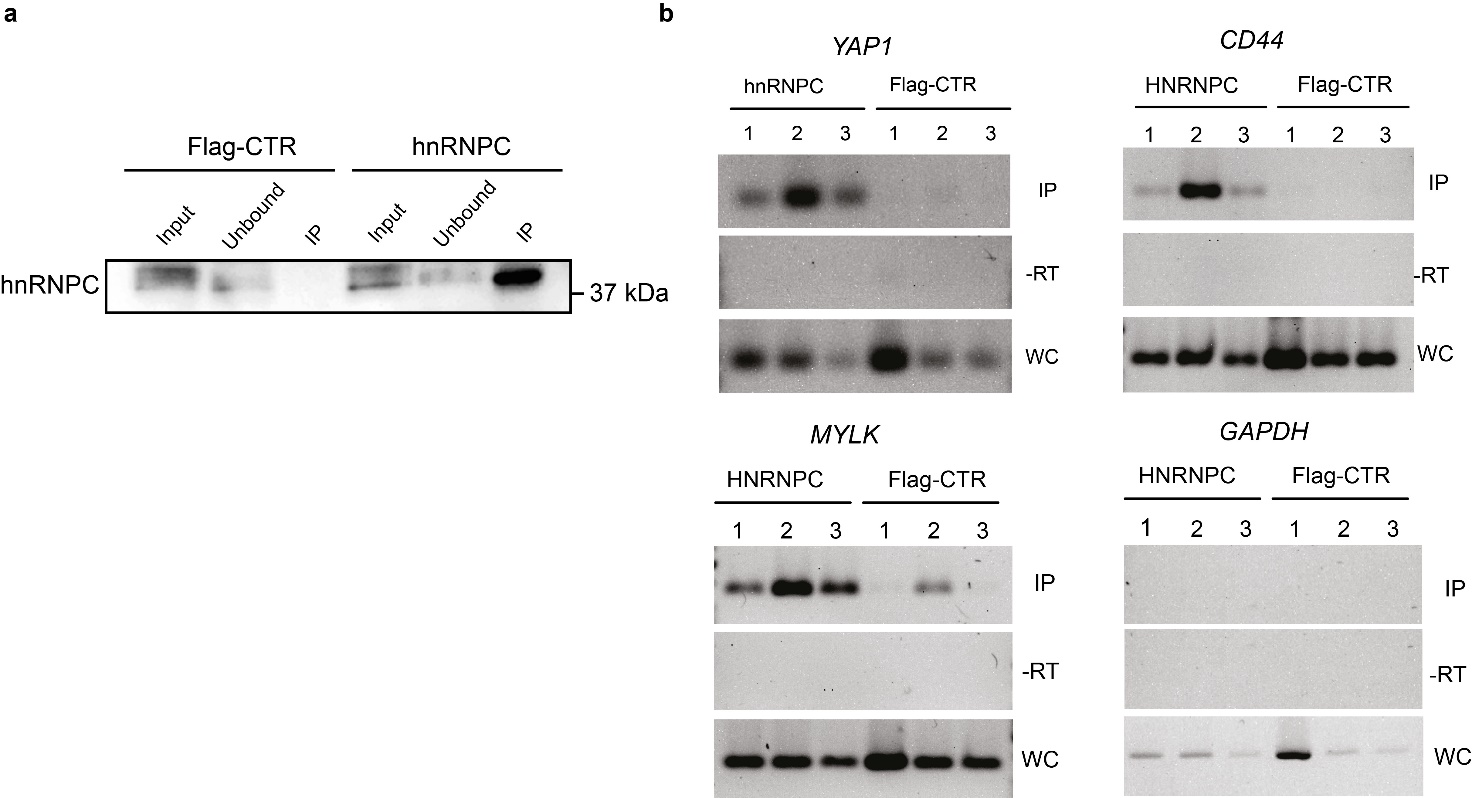

**Supplementary Figure 10. a)** Western blot analysis of hnRNPC protein in the total lysate of normal human dermal fibroblasts (input), in hnRNPC immunoprecipitated sample (IP) and unbound fraction following iCLIP protocol. Flag immunoprecipitation was used as negative control. **b)** Analysis of the indicated transcripts in PCR-amplified iCLIP cDNA libraries. Whole cell lysates (WC) and Flag-immunoprecipitated samples were used as positive and negative controls, respectively. -RT represents the negative control in the absence of amplification.

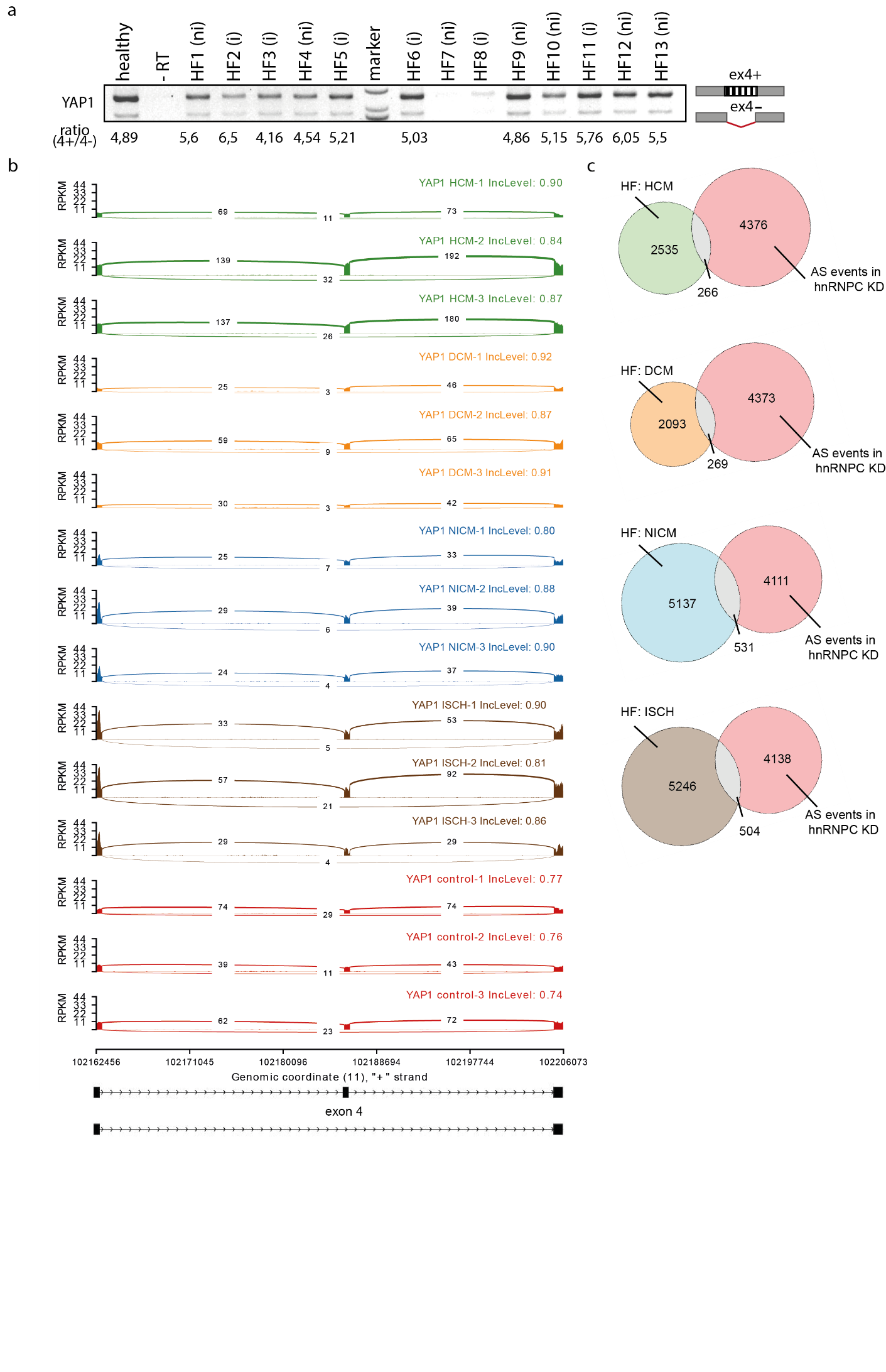

**Supplementary Figure 11. a)** RT-PCR of YAP1 in samples of human healthy and diseased (HF) hearts. HF (i) and HF (ni) indicate samples obtained from patients diagnosed with either ischemic or non-ischemic HF, respectively. Top band: exon 4-included YAP1 mRNA. Bottom band: exon 4-excluded YAP1 mRNA. -RT represents the negative control in the absence of amplification. **b)** Sashimi plot depicting the inclusion level of YAP1 exon 4 in human healthy (control) or failing hearts. Failing hearts are obtained from patient diagnosed with hypertrophy (HCM) (GSE141910), dilated cardiomyopathy (DCM) (GSE141910), non-ischemic (NICM) (GSE108157), or ischemic cardiomyopathy (ISCH) (GSE108157). Healthy human hearts are obtained from the dataset GSE141910. **c)** Venn diagram representations of the AS events determined by hnRNPC depletion in KD NHDF that are also found in failing human hearts (HCM, DCM, NICM, ISCH) and not in control heart. See also **Supplementary Data 9**.

**Supplementary Material and Methods:**

1. **Surgical induction of mouse myocardial infarction (MI)**

MI was induced by ligating the left anterior descending coronary artery in C57BL/6 adult mice (9-10 weeks), as previously described ^2^. Mice hearts were collected 4 days and 21 days after surgery, time points corresponding to acute MI and chronic MI, respectively. The infarcted area downstream of the ligation and the equivalent region in sham-operated control, were isolated. After collection, the isolated heart tissues were washed in PBS and processed for histology.

Experiments involving mice were approved by the Instituto de Biologia Molecular e Celular – Instituto de Engenharia Biomédica (IBMC-INEB) Animal Ethics Committee and the National Direção Geral de Veterinária (permit no: 022793), and conformed with Directive 2010/63/EU of the European Parliament.

2. **Protein immunoprecipitation from human heart tissue**

For immunoprecipitation of hnRNPC-binding proteins, 100 mg tissue from the human heart apex was used for each IP and control sample.

**2.1 Tissue’s lysate preparation:** Samples were thawed on ice, then washed in PBS before tissue dissection into small pieces on a glass plate placed on ice. The tissue was mechanically homogenized in 1 ml lysis buffer [150 mM NaCl, 50 mM TRIS pH 7.5, 0.5 % deoxycholate, 0.5% Tergitol (Sigma Aldrich, NP40S), protease and phosphatase inhibitor cocktail (10%, Sigma PPC1010)] using 3.0 mm Zirkonia beads (SKU:D1032-30) and a Beadbug homogenizer (Benchmark scientific, catalog number: D1030-E) for 2 min at 300 rpm. The sample was incubated on ice for 10 min after each homogenization cycle. The procedure was repeated until the sample was completely homogeneous (~60 min, ~6 cycles). Then, the lysate was centrifuged at 10,000 g for 10 min at 4 °C and the total lysate served as the input for western blot analysis.

**2.2 Beads preparation**

The storage buffer was removed from 50µl Dynabeads Protein G (**Supplementary Table 2**). Then, primary antibody (hnRNPC 4F4, FHL2, PDLIM5 antibody or isotype control IgG; 5 µg per sample) dilution in 200 µl Ab binding buffer (catalog number 10007D) was added and incubated with the Dynabeads for 45 min at 4°C with gentle rotation. The Ab-conjugated-beads were washed with 200 µl Ab Binding Buffer before proceeding with immunoprecipitation step.

**2.3 Immunoprecipitation**

An equal amount of protein lysate for each sample was incubated with antibody-conjugated-beads, overnight under rotation at 4°C. Then, the immunoprecipitated samples were washed three times in wash buffer (150 mM NaCl, 50 mM TRIS pH 7.5) and aliquots of the samples were collected for western blotting. For elution with 8 M urea, the samples were incubated with 100 µl urea on a shaker for 20 min at room temperature (RT); the elution was repeated three times and cumulatively collected.

**3. Mass spectrometry analysis**

Mass spectrometry was applied to hnRNPC immunoprecipitates from human healthy and failing (HF) heart samples to determine the composition of hnRNPC interactome. The samples were digested with trypsin using the filter-aided sample preparation (FASP) method ^3^. Briefly, samples were reduced with 100 mM dithiothreitol at 60°C for 30 min, transferred to 30 kDa MWCO Pall Nanosep centrifugation filters (Sigma-Aldrich), washed repeatedly with 8 M urea and once with digestion buffer [1% sodium deoxycholate (SDC) in 50 mM triethylammonium bicarbonate (TEAB)] prior to alkylation with 10 mM methyl methanethiosulfonate in digestion buffer for 30 min. Digestion was performed in digestion buffer supplemented with 0.5 µg Pierce MS grade trypsin (Thermo Fisher Scientific) at 37°C and incubated overnight. An additional portion of trypsin was added and incubated for a further 2 hours and peptides were collected by centrifugation.

**3.1 TMT-labeling, MS-analysis and database search**

Digested peptides were labeled using TMT pro16-plex isobaric mass tagging reagents (Thermo Scientific), according to the manufacturer instructions. The samples were combined into two TMT sets and sodium deoxycholate was removed by acidification with 10% TFA. The set included the following samples:

1) diseased 1 - IP hnRNPC

2) diseased 1 - IgG

3) diseased 2 - IP hnRNPC

4) diseased 2 - IgG

5) healthy 1 - IP hnRNPC

6) healthy 1 - IgG

7) healthy 2 - IP hnRNPC

8) healthy 2 – IgG

9) diseased 1 – IP FHL2

10) diseased 1 – IP PDLIM5

11) diseased 1 – IgG

12) diseased 2 – IP FHL2

13) diseased 2 – IP PDLIM5

14) diseased 2 - IgG

The sets were pre-fractionated into 20 fractions by basic reversed-phase chromatography (bRP-LC) using a Dionex Ultimate 3000 UPLC system (Thermo Fischer Scientific). Peptide separations were performed using a reversed-phase XBridge BEH C18 column (3.5 μm, 3.0x150 mm, Waters Corporation) and a linear gradient from 3% to 40% of solvent B over 18 min followed by an increase to 100% of solvent B over 5 min. Solvent A consisted of 10 mM ammonium formate buffer at pH 10.00 and solvent B was 90% acetonitrile and 10% 10 mM ammonium formate at pH 10.00. The fractions were concatenated into 10 sequential fractions (1+11, 2+12, up to 10+20), dried and reconstituted in a solution of 3% acetonitrile, 0.2% formic acid.

**3.2 nLC-MS/MS**

The fractions were analyzed on an orbitrap Fusion™ Lumos™ Tribrid™ mass spectrometer interfaced with an Easy-nLC1200 liquid chromatography system (Thermo Fisher Scientific). Peptides were trapped on an Acclaim Pepmap 100 C18 trap column (100 μm x 2 cm, particle size 5 μm, Thermo Fischer Scientific) and separated on an in-house packed analytical column (75 μm x 30 cm, particle size 3 μm, Reprosil-Pur C18, Dr. Maisch) using a linear gradient: 5% to 33% solvent B for 77 min, followed by an increase to 100% solvent B for 3 min, and 100% solvent B for 10 min at a flow of 300 nL/min. Solvent A was 0.2% formic acid and solvent B was 80% acetonitrile, 0.2% formic acid. Precursor ion mass spectra were acquired at 120,000 resolution and MS/MS analysis was performed in a data-dependent multi-notch mode where the CID spectra of the most intense precursor ions were recorded in an ion trap at a collision energy setting of 35 for 3 sec (‘top speed’ setting). Precursors were isolated in the quadrupole with a 0.7 m/z isolation window, charge states 2 to 7 were selected for fragmentation, and dynamic exclusion was set to 45 s and 10 ppm. MS3 spectra for reporter ion quantitation were recorded at 50,000 resolution with HCD fragmentation at a collision energy of 55 using the synchronous precursor selection.

**3.3 Proteomic Data Analysis**

The data files for each set were merged for identification and relative quantification using Proteome Discoverer version 2.2 (Thermo Fisher Scientific). The search was made against the *Human* Swissprot Database version June 2019 (Swiss Institute of Bioinformatics, Switzerland) using Mascot version 2.5.1 (Matrix Science) as a search engine with precursor mass tolerance of 5 ppm and fragment mass tolerance of 0.6 Da. Tryptic peptides were accepted with zero missed cleavage, variable modifications of methionine oxidation and fixed cysteine alkylation, and TMT-label modifications of the N-terminal and lysine were selected. The reference samples were used as a denominator and to calculate the ratios. A percolator was used to validate the identified proteins. TMT reporter ions were identified in the MS3 HCD spectra with 3 mmu mass tolerance. The quantified proteins were filtered at 1% FDR and grouped by sharing the same sequences to minimize redundancy. Only peptides unique for a given protein were considered for protein quantification, excluding those common to other isoforms or proteins of the same family.

Proteomics analysis was performed on hnRNPC-immunoprecipitated samples and IgG-immunoprecipitated samples (negative controls) from two healthy and two failing (HF) human heart tissues. The ratio of IP/IgG abundances was calculated for each protein found in each sample. A protein was considered successfully immunoprecipitated when IP/IgG > 1.5 in both biological replicates (healthy1-healthy2 or HF1-HF2). Proteomics analysis on FHL- and PDLIM5-immunoprecipitated samples and IgG- immunoprecipitated samples from two HF samples was performed using the same method.

Proteomaps **(Fig. 2c)** (<https://bionic-vis.biologie.uni-greifswald.de/>) ^4,5^ were used to represent the quantitative composition of the hnRNPC interactome in healthy and failing human hearts, with a focus on protein function. The polygonal areas in the proteomaps represent the protein abundances in the immunoprecipitated sample as detected by TMT-MS analysis. Functionally related proteins are distributed in similarly colored regions. Proteomaps are based on the KEGG Pathways gene classification.

**4. Proximity Ligation Assay**

The proximity ligation assay (PLA) was performed on fixed heart tissue slices or on fixed cells using DuoLink PLA technology probes and reagents (**Supplementary Table 2**), according to the manufacturer’s protocol. Fixed samples were permeabilized with 0.5% Triton X-100 in PBS for 10 min (tissue slices) or 0.2% Triton X-100 in PBS for 5 min (cells). The samples were then washed three times with PBS and incubated with blocking solution for 1 h at 37°C in a humidified chamber. Primary antibodies were prepared in antibody diluent and incubated overnight at 4°C (**Supplementary Table 2**). The samples were washed twice with buffer A for 5 min and then incubated with the PLA probes (anti-mouse MINUS and anti-rabbit PLUS) in antibody diluent for 1 h at 37°C. After two washes for 5 min each with buffer A, a ligation step was performed by incubating the samples with ligase enzyme diluted in ligation buffer for 30 min at 37°C in a humidified chamber. After 2 washes of 5 min each with buffer A, the samples were incubated with polymerase enzyme diluted in amplification buffer for 100 min at 37°C, in humid chamber. After two washes for 10 min each with buffer B, and a final wash of 1 min with 0.01% of buffer B, the slides were mounted with Duolink *in situ* mounting medium containing DAPI. Each experiment was performed with a pair of antibodies of different species (mouse and rabbit). Negative control experiments were performed where only one antibody or none were incubated with the PLA probes.

**5. iPSC cardiac differentiation**

The human induced pluripotent stem cell (iPSC) line DF 19-9-7T was purchased from WiCell (Madison, WI, USA). iPSCs were cultured on Matrigel® Matrix-coated (1:100 in DMEM/F12, Corning) Growth Factor Reduced plates in complete Essential 8™ Medium (Life Technologies) containing penicillin/streptomycin (0.5 %, VWR). For passaging, the cells were dissociated using TrypLE Express (Life Technologies) and supplemented with Rock Inhibitor Y27632 (2.5µM, Selleckchem).

Cardiac differentiation was performed as previously described ^6^, with slight modifications. The medium was changed daily until the cells formed a monolayer. At day 0 of cardiac differentiation, 100% confluent cells were incubated with mesoderm induction medium: RPMI 1640 (Sigma-Aldrich) supplemented with penicillin/streptomycin, L-glutamine (2 mmol/L, Biowest), B-27™ supplement minus insulin (1X, Thermo FisherScientific) and CHIR99021 (8 µM, Sigma-Aldrich). At day 2, the medium was replaced with RPMI plus B-27 supplement minus insulin and IWP-2 (5 µmol/L, Selleck chemicals). At day 4, the medium was changed to RPMI plus B27 supplement minus insulin and substituted every other day until the cells started beating. Around day 8-10, after the iPSC-derived cardiomyocytes (CMs) started beating, the medium was substituted with RPMI supplemented with B27 supplement plus insulin (1X, Thermo Fisher Scientific) and replaced every 2-3 days over the differentiation time to assure cell beating.

**6. Ribopuromycilation**

Ribopuromycilation assay was performed to visualize the active sites of translation following Procedure A in the Original RPM protocol, as previously described ^7,8^. A schematic representation of the protocol can be found in **Supplementary Fig. 5d**. All the buffers were freshly prepared before use.

Beating cardiomyocytes between day 20 and day 30 of differentiation were incubated with cycloheximide (CHX), a chain elongation inhibitor, at the final concentration of 355 µM for 5 min at RT. Labeling medium containing CHX (355 µM) and puromycin (PMY) (91 µM final concentration) was added for 5 min at 37 C. PMY is incorporated into nascent peptides while CHX prevents PMY-nascent chain release from the ribosomes. For each experiment, a control without puromycin addition was performed. After labeling, the cells were washed with cold PBS. All subsequent steps were performed on ice. Extraction buffer (Triton 0,01%, MgCl2 5mM, KCl 25nM, CHX 355 µM, protease and phosphatase inhibitors 1% v/v, Tris-HCl 50mM pH 7.5) was slowly added and incubated for 2 min. Then, the cells were washed with wash buffer (MgCl2 5mM, KCl 25nM, CHX 355 µM, protease and phosphatase inhibitors 1% v/v, Tris-HCl 50mM pH 7.5) and fixed with 4% PFA for 15 min at RT. After fixation, the cells were washed three times with PBS and immunostaining was performed.

**7. Decellularization of human myocardial tissue and cell seeding on decellularized ECM**

The decellularization protocol was performed as previously described ^9,10^ with slight modifications. First, the frozen healthy and diseased heart samples were thawed and sliced into small pieces using a 2x2 mm grid as a reference. The explants were then immersed in mild decellularization solutions and mixed by shaking at 300 rpm (Eppendorf Thermomixer C 80 with orbit diameter of 3 mm) at 25°C. The specimens were first immersed for 18 h in hypotonic buffer (10 mM Trizma Base Sigma / 0.1% EDTA 82 Applichem, pH 7.8), then for 24 h in detergent solution (0.5% Sodium Dodecyl Sulfate 83 (SDS, Sigma) / 10 mM Trizma Base Sigma, pH 7.8) and for 1 h in hypotonic wash buffer (10 mM TrizmaBase Sigma, pH 7.8). DNAse treatment [50 U/mL 85 DNAse I (StemCell Technologies) in 10 mM Tris HCl, pH 7.8] was performed for 3 h at 37°C. Finally, the samples were washed overnight in PBS (12 rpm, GFL Shaker 3014).

The decellularized samples were mechanically homogenized in 400 µl sterile PBS supplemented with 1% penicillin/streptomycin using 3.0 mm Zirkonia beads (SKU:D1032-30) and a Beadbug homogenizer (**Supplementary Table 2**). dECM matrices were homogenized until full dissociation with cycles of 1 min at 3,000 rpm and 2 min on ice. A Pierce BCA Protein Assay Kit (ThermoFisher) was used to quantify the proteins in the homogenized samples. Decellularized matrices were preserved in PBS supplemented with 1% Penicillin/Streptomycin (Diagnovum) at 4°C.

The coating was performed by incubating 12 mm coverslips with 100 µl/cm^2^ dECM solution (5µg/ml) at 37°C overnight in PBS-1%Penincilin/Streptomycin solution. Coating solution was removed immediately before cell seeding. Beating iPSC-CMs (day 15-20) were dissociated in TryPLE, re-suspended in RPMI medium supplemented with B27 supplement minus insulin and Y27632, and seeded onto ECM-coated-coverslips. Then, 24 h after seeding and when the cells re-started beating, the medium was replaced with RPMI supplemented with B27 supplement plus insulin. 48 h after seeding, the ribopuromycilation assay was performed as previously described (**Supplementary Method 6**). Cells were then fixed and kept at 4 °C until PLA was performed. Strategy depicted in **Fig. 5c**.

**8. Cell treatment, embedding in fibrin hydrogels and micropatterning**

**8.1 Cell treatments**

For the treatment with inhibitors of cytoskeletal tension, cells were incubated with Latrunculin-A (1 μM) for 30 min or with Rock inhibitor (Y27632) (50 μM) for 4 h, at 37°C and 5% CO_2_. After treatment, cells were washed twice with PBS, harvested with TryPLE and the proteins were extracted as described in Method 12.

**8.2 3D Fibrin hydrogel preparation, 2D elastic surfaces and cells seeding**

3D fibrin gels were produced as previously described ^1^. Fibrinogen and thrombin were purchased from Baxter. DMEM (50μL) containing 2x10^5^ normal human dermal fibroblasts (NHDFs) was embedded in 50μL of 40 U/mL thrombin-CaCl_2_ solution. The resultant solution was then quickly mixed 1:1 with fibrinogen 100 mg/mL, 34 mg/mL, 10 mg/mL or 4 mg/mL to generate clots with fibrinogen concentrations of 50 mg/mL, 17 mg/mL, 5 mg/mL and 2 mg/mL, respectively, with a constant thrombin concentration of 20 U/ml. The 3D fibrin gels with embedded cells were polymerized for 1 h at 37°C before the addition of complete growth medium (DMEM) supplemented with aprotinin (3000 KIU/mL). The 3D fibrin hydrogels were kept at 37 °C and 5% CO_2_ in a humidified incubator until processed.

Thirty-five mm high elastically supported surface (ESS) μ-Dishes with a stiffness of 28 or 1.5 kPa were purchased from iBIDI (Munich, Germany). iPSC-derived cardiomyocytes were seeded on Matrigel®- coated μ-Dish ESS in RPMI medium supplemented with B27 supplement minus insulin and Y27632. After 24 hours, the medium was changed to B27 supplement plus insulin and 48 hours after seeding the cells were fixed with 4% PFA for 15 min at RT. After fixation, the cells were washed three times with PBS and immunostaining was performed.

**8.3 Micropatterning**

Fibronectin-coated micropatterned slides featuring different areas or patterns (ref: 10–950–10–18; **Supplementary Table 2**) were purchased from CYTOO (CYTOO, Grenoble, France).  After 24 h culture in complete medium on the microarrays, the NHDFs were fixed in 4% PFA for 15 min at RT and analyzed by immunofluorescence; n≥8 squares of each type were analyzed *per* experiment (N=3).

**9. Cell siRNAs transfection**

*HNRNPC* knockdown was performed as previously described ^11^. In brief, NHDF cells and iPSC-derived cardiomyocytes (CMs) were transfected using two different human hnRNPC1/C2 stealth select small interfering RNA (siRNA) (Invitrogen, HSS179304 and HSS179305) and a stealth siRNA negative control low GC (Invitrogen) at a final concentration of 10 nM. The transfection was performed using Lipofectamine RNAiMAX (Invitrogen) transfection reagent according to the protocol “RNAiMAX Reverse Transfections Lipofectamine” provided by ThermoFisher Scientific (**Supplementary Table 2)**.

For each well of a 6-well plate to be transfected, siRNA duplex-Lipofectamine™ RNAiMAX complexes were prepared as follows: the growth media was replaced by 30 pmol siRNA diluted in 500 µl Opti-MEM® Medium (Gibco^TM^, ThermoFisher Scientific) without serum and was gently mixed. Lipofectamine™ RNAiMAX was gently mixed before use, and 5 µl was added to each well containing the diluted siRNA molecules. The resultant solution was gently mixed and incubated for 20 min at RT. NHDFs and iPSC-CMs were diluted in Opti-MEM® Medium at a concentration of 1x10^5^ cells/ml; then, 2.5 ml of the diluted cells were uniformly added to each well containing siRNA-Lipofectamine™ RNAiMAX complexes. Cells and transfection reagents were incubated at 37°C in a CO_2_ incubator for 24 h followed by the addition of 2 ml of complete growth medium (DMEM containing 10% FBS, 2mM L-glutamine and 1% penicillin/streptomycin for NHDFs; essential 8™ Medium containing penicillin/streptomycin 0.5% and supplemented with ROCK inhibitor Y27632 1:4000. The knockdown efficiency was assessed 96 h after transfection (**see Fig.7a and Supplementary Fig.7a**).

**10. RNA immunoprecipitation from human heart tissue**

For immunoprecipitation of hnRNPC RNA complexes, 100 mg tissue from the human heart apex was used for each IP and control sample. All the buffers were prepared before use. A schematic representation of the experimental strategy can be found in **Fig. 4a**.

**10.1 Tissue’s lysate preparation:** The samples were thawed on ice, washed twice in PBS and then the tissues were cut into small pieces (2mm diameter) on a glass plate placed on ice. Cross-linking was achieved by soaking the samples in 1% (v/v) formaldehyde for 15 min in 10 ml PBS at RT with gentle rotation. The cross-linking reaction was quenched by adding 200mM TRIS. The samples were centrifuged at 50 g for 1min at 4°C, washed twice in PBS and then mechanically homogenized in 1ml Polysome Lysis Buffer (PLB) [20mM Tris buffer (pH 7.5), 100 mM KCl, 5mM EDTA at pH 8, 0.5% Tergitol (Sigma Aldrich, NP40S), 2 mM DTT, 50 U/mL RNase OUT, heparin 0.2 mg/mL, protease and phosphatase inhibitor cocktail (10%, Sigma PPC1010)] using 3.0 mm Zirkonia beads (SKU:D1032-30) and a Beadbug homogenizer for 2 min at 300 rpm. The sample was cooled on ice for 10 min after each homogenization cycle. The procedure was repeated until the sample was completely homogeneous (~60 min, ~6 cycles). Then, the samples were sonicated (focused-ultrasonicator M220, Covaris) for five cycles, consisting of: peak power 50, duty factor 20%, 200 cycles, 20 sec, plus 30 sec delay, at 4 °C. The resulting lysates were centrifuged twice for 10 mins each at 4 °C and 14,000 x g and the supernatant collected. Aliquots (1:5 of total lysate) of the supernatant were collected to serve as the input for western blot analysis and total RNA extraction.

**10.2 Beads preparation:** The storage buffer was removed from 100 µl Dynabeads Protein G (immunoprecipitation Kit, catalog number 10007D) using a magnetic rack. Then, the beads were equilibrated by washing them three times with 400 µl NT2-coupling buffer [50 mM Tris–HCl at pH 7.5, 500 mM NaCl, 1 mM MgCl2, with 1% Tergitol, 5% BSA, 0.02% sodium azide, 0.02 mg/mL heparin]. The primary antibody (hnRNPC 4F4 antibody, sc-32308 or isotype control IgG mouse, sc-2025, Santa Cruz Biotech, 10 µg/sample; **Supplementary Table 2**) was then diluted in 400 µl NT2-coupling buffer and incubated with the Dynabeads for 1 h at 4°C with gentle rotation. The antibody-conjugated-beads were washed twice with 400µl NT2-RIP buffer [50 mM Tris–HCl pH 7.5, 500 mM NaCl, 1 mM MgCl_2_, with 1% Tergitol, 50 U/mL RNase OUT, 2 mM DTT, 30 mM EDTA at pH 8, heparin 0.02 mg/mL].

**10.3 Immunoprecipitation:** The antibody-conjugated beads were incubated with an equal volume of cross-linked lysate in NT2-RIP buffer (1:10) for each sample, overnight and under gentle rotation at 4°C. The IP-beads were resuspended in ice-cold NT2 buffer [50 mM Tris–HCl at pH 7.5, 500 mM NaCl, 1 mM MgCl_2_, and 1% Tergitol] and transferred to a new 1.5 mL conical microtube. The IP-beads were washed four times in NT2 buffer by shaking the tubes and aliquots (~150 µl) were collected for western blot analysis.

**10.4 Elution and reverse cross-linking:** The input samples were thawed (aliquot for total RNA extraction) and reverse cross-linking was carried out by resuspending the IP-beads samples and input sample in 250 µl SDS-EDTA elution buffer [50 mM Tris–HCl at pH 8, 100 mM NaCl, 10 mM EDTA, 1% (w/v) SDS] supplemented with 60 µg Proteinase K per sample. The samples were then incubated for 1.5 h at 55 °C with shaking (800 rpm).

**10.5 RNA extraction:** A total of 400 µl TRIzol (Invitrogen) was added to the samples and incubated for 5 min on ice. Then, 80 µl chloroform was added and incubated for 30 sec with shaking before centrifugation at 10,.000 g for 10min at 4°C. The upper aqueous phase was removed and transferred to a new tube before the addition of 50 µl Salt I (Millipore), 15 µl Salt II (Millipore), 4 µl Precipitate Enhancer (Millipore) and 850 µl absolute ethanol (**Supplementary Table 2)**. The RNA was precipitated overnight at - 80°C, then centrifuged at 14,000 g for 30 min at 4 °C. The supernatant was removed, and the pellet was washed twice with ice cold 75% ethanol by centrifugation at 5 min at max speed at 4°C. Finally, the pellet was air-dried for 2 min at RT and then resuspended in 15 µl nuclease free water.

**11. Quantitative real-time PCR (qPCR)**

For RNA extraction from human heart samples, tissues were thawed from -80°C and total RNA was extracted by standard phenol:chloroform (TRIzol protocol, according to manufacturer’s instructions) phase separation, followed by precipitation in isopropanol and EtOH wash. For RNA extraction from adherent cells, cells were harvested from culture plates by TryPLE, cell pellet was washed twice in DPBS and total RNA was isolated using a High Pure RNA Isolation Kit (Roche) according to the manufacturer's instructions. After RNA isolation, samples were treated with Dnase (Roche) to remove genomic DNA contamination and 1 μg of total RNA was transcribed into cDNA using a Transcription First Strand cDNA Synthesis Kit (Roche) and analyzed by qPCR using a SYBR Green I Master Kit (Roche) and a LightCycler 480 Real-Time PCR System (Roche). The samples were loaded in triplicate and quantified by ∆Ct method normalized to reference GAPDH expression using LightCycler 480 Software release 1.5.0. Non-template controls (NTC) containing no cDNA were included in each reaction. The primer sequences used are listed in **Supplementary Table 2**.

**12. Western blotting**

For total protein extraction from human heart tissue, samples were thawed from -80°C, placed on a glass plate on ice, finely cut into small pieces and mechanically disrupted by pipetting and vortexing. The homogenized heart tissue was then incubated with RIPA lysis buffer (Merk Millipore, 20-188) supplemented with protease and phosphatase inhibitor cocktails (1% v/v, Sigma-Aldrich) for 30 min on ice, vortexing every 10 min. The lysate was then centrifuged at 4°C for 15 min at 16,000 g. The supernatant was collected in a new eppendorf tube and stored at -80°C until further use.

For total protein extraction, the cells were detached with TrypLE Express (Life Technologies, 12604-013), and the pellet was washed with PBS and incubated in RIPA buffer as described above.

Compartment protein extraction was performed using an NE-PERtm Nuclear and Cytoplasmic extraction kit (**Supplementary Table 2**) following the manufacturer’s specifications. A Pierce BCA Protein Assay Kit (ThermoFisher) was used for protein quantification, with albumin serum bovine standards for calibration. The respective absorbances were read using a Multiskan™ GO Spectrophotometer.

Protein extracts (5-10 µg) were boiled for 5 min at 95 °C, loaded on 10% Mini-PROTEAN® TGX™ Precast Protein Gels and separated at 100 V in Tris/Glycine/SDS buffer in a Mini-Protean Tetra System (Bio-Rad). The proteins were transferred to polyvinylidene difluoride membranes (PVDF, Bio-Rad) using a semi-dry Trans-Blot Turbo transfer system (Bio-Rad). The membranes were immersed in 5% skim milk in TBST to block unspecific sites for 1 h, followed by primary antibody incubation in 5% BSA in TBST with gentle rotation overnight at 4°C. After three washes in TBST, the membranes were incubated for 1 h at RT with the secondary HRP-conjugated antibody (Sigma-Aldrich) in 5% BSA in TBST. Clarity™ Western ECL Substrate (Bio-Rad) and a ChemiDoc MP Imaging System (Bio-Rad) were used to detect Chemiluminescence. Band intensities were quantified using Bio-Rad Image Lab software.

**13. RNA sequencing, RIP-seq analysis and AS analysis**

**13.1 Sequencing**

A sequencing library was prepared using an NEBNext Ultra II Directional Kit (New England Biolabs, MA, USA). For RIP-sequencing, immunoprecipitated RNA was used as an input into the total RNA library preparation protocol; control input RNA was first rRNA depleted using a QIAseq Fastselect HMR Kit (Qiagen, Germany). For the alternative splicing analysis, 200-300 ng total RNA was used as an input into the polyA enrichment module protocol. The samples were fragmented and transcribed into cDNA. Following universal adapter ligation, the samples were barcoded using dual indexing primers. The RNA resultant from the RIP of heart tissue was sequenced to 25-35 million single-end 75bp reads while the RNA resultant from the NHDF transfection with hnRNPC siRNA and respective control constructs was sequenced to >50 million paired-end 75 bp reads/sample. The samples were sequenced on an Illumina Nextseq 550 sequencer (Illumina, CA, USA).

A quality check of raw fastq reads was carried out by FastQC ^12^. The adapters and quality trimming of the raw fastq reads was performed using Trimmomatic v. 0.36 ^13^ with settings CROP:250 LEADING:3 TRAILING:3 SLIDINGWINDOW:4:5 MINLEN:35. The trimmed RIP-Seq and RNA-seq reads were mapped against the human genome (hg38) and Ensembl GRCh38 v.94 annotation using STAR v. 2.5.3a ^14^ as splice-aware short read aligner and default parameters, with the exception of outFilterMismatchNoverLmax 0.1 and twopassMode Basic. Quality control after alignment including assessment of uniquely and multi-mapped reads, read coverage distribution, level of read duplication, rRNA contamination, genomic regions of mapped reads and gene biotypes was performed using several tools, namely RSeQC v2.6.2 ^15^, Picard toolkit v2.18.27 (GitHub Repository. http://broadinstitute.github.io/picard/), Qualimap v.2.2.2 ^16^ and FastQ Screen v0.13.0 ^17^.

**13.2 RIP-seq analysis**

Counts of mapped reads to specific gene features were made using a modified Ensembl GRCh38 v.94 annotation GTF file. The annotation file was modified as follows:

**13.2.1** Four individual GTF files were derived from original annotation GTF file: (1) only for regions specified as a feature "gene", (2) for regions specified as a feature "exon", (3) for regions specified as a feature "three_prime_utr" and (4) for regions specified as a feature "five_prime_utr" (only for chromosomes 1-23, X, Y and MT).

**13.2.2** Two derived GTF files for genes and exons were subtracted using bedtools subtract v2.29.0 (Quinlan et al.,2018) considering the strandness of the alignment as a parameter to create the GTF file with only intronic regions; regions annotated as RNA (antisense_RNA, bidirectional_promoter_lncRNA, 3prime_overlapping_ncRNA, lincRNA) were excluded from GTF file.

**13.2.3** As a next step, the 3'UTR and 5'UTR regions in the GTF files were subtracted from the exonic GTF file using bedtools subtract.

**13.2.4** The subtracted exonic GTF, 5'UTR GTF, 3'UTR GTF and intronic GTF were joined together into one GTF file, which was sorted by positions, resulting in a final GTF file that was used to count the mapped reads. Read counts for four specific genomic features (CDS - exons without UTR regions, introns, 3'UTR and 5'UTR) were gathered using featureCounts v1.5.2^18^. The set included the following samples:

1) diseased 1 (HF1) – input (total lysate)

2) diseased 1 (HF1) - IP hnRNPC

3) diseased 1 (HF1) - IgG

4) diseased 2 (HF2) - input

5) diseased 2 (HF2) – IP hnRNPC

6) diseased 2 (HF2) - IgG

7) diseased 3 (HF3) – input (total lysate)

8) diseased 3 (HF3) – IP hnRNPC

9) diseased 3 (HF3) - IgG

The input RNA samples were normalized on sequencing depth and further used as normalization factors for IP and IgG sample normalization. IP and IgG samples were further normalized on the length of each feature on a logarithmic scale with base 2. Exon counts were represented by the sum of CDS, 3'UTR and 5'UTR, while introns only included intronic reads. The total transcript expression was defined as the sum of the exonic and intronic reads. The immunoprecipitated transcripts were considered statistically significant when the IP/IgG ratio > 5 and the transcript total expression > 200 in at least two out of three replicates of the IP samples. To distinguish between exons-enriched targets and introns-enriched targets, we established a ratio of intronic/exonic reads on a logarithmic scale with base 2 and set the cutoff as log_2_(intron/exon) > 1 (intron-enriched) or log_2_ (intron/exon) < -1 (exon-enriched) and the total transcript expression > 200. To visualize all four genomic features (CDS - exons without UTR regions, introns, 3'UTR and 5'UTR) in pie plots, their counts were handled separately without any cutoffs or filtering. Volcano and bar plots were created using the ggplot2 v3.3.1 ^19^ library, and pie plots were generated using the plotrix v3.7-8 ^20^ library.

**13.3 Alternative splicing (AS)**

The experimental design included the following samples:

CTR x3 : NHDFs transfected with a stealth siRNA negative control low GC (Invitrogen)

KD1 x3: NHDFs transfected with hnRNPC1/C2 siRNA HSS179304 (Invitrogen)

KD2 x3: NHDFs transfected with hnRNPC1/C2 siRNA HSS17930 (Invitrogen).

Alternative splicing (AS) analysis was carried out by rMATS v3.1.0 ^21^, which detects differentially used exons by comparing exon-inclusion levels based on the number of junction reads. In this case, all replicates from the KD1 and KD2 conditions were treated as one condition containing 6 replicates and were compared to three replicates of CTR condition. rMATS was then run with settings -t paired --readLength 75 --libType fr-secondstrand. Sashimi plots depicting skipped exon AS events were generated using rmats2sashimiplot v2.0.2 (https://github.com/Xinglab/rmats2sashimiplot), whereby representative replicates of KD and CTR conditions can be visualized. The clusterProfiler v3.12.0 ^22^ R library was used for Gene Ontology (GO) analysis. The input list of genes uploaded for GO analysis consisted of a pool of genes from all individual rMATS AS event outputs (Alternative 5' start site, Alternative 3'start site, Skipped exons, Retained introns and Mutually exclusive exons), filtered based on FDR ≤ 0.01 and Inclusion level difference (ΔPSI) ≥ 0.1 cutoffs.

AS analysis in human heart samples was carried out by rMATS v3.1.0. Three samples for each diseased condition (HCM, DCM, ISCH, NICM) were analysed and compared to healthy controls (GSE141910). The samples analysed belong to the following published datasets: GSE108157 and GSE141910. The differentially spliced events were calculated based on FDR ≤ 0.05 and inclusion level difference (ΔPSI) ≥ 0.1 cutoffs.

**14. Cytoscape Analysis and GO analysis**

The EnrichR enrichment analysis tool was used (<http://amp.pharm.mssm.edu/Enrichr>) ^23,24^, for the molecular function GO analysis of upregulated genes in mouse and human pathological hearts (**Fig. 1a**). Cytoscape 3.7.2 software ^25^ was used to analyze the enrichment of gene annotation (GO terms) in GO networks or sub-networks on data obtained by RNA-seq. The Cytoscape plug-in ClueGO version 2.5.6 ^26^ was used to visualize the biological terms of genes clustered in functionally grouped networks. The ClueGO plugin CluePedia version 1.5.6 ^27^, was used to represent the genes and to integrate them in the ontology network primarily built with ClueGO. For the cellular component GO analysis of hnRNPC interactome in human healthy and pathological hearts (**Fig. 2d**), GO term fusion, GO-Tree interval of 4-10, p value ≤ 0,01 were selected as filters. The cellular component and biological process GO analysis of hnRNPC-RNAs targets obtained by RIP-seq GO term fusion, GO-Tree interval of 4-10, p value ≤ 0,05, were selected as filters.

The WikiPathways database ^28^ was used for pathway enrichment analysis performed on differentially spliced genes in hnRNPC silenced cells, and filtered by p-value ≤ 0.05. For all statistical analyses, the Kappa score was set to 0.3 and an Enrichment/Depletion (Two-sided hypergeometric test) test with Bonferroni pV correction was selected as an advanced statistical option in the software.

POSTAR2 database (<http://lulab.life.tsinghua.edu.cn/postar/>) ^29^ was used to analyze transcripts which harbor hnRNPC binding sites.

To visualize the components of the Hippo pathway found among hnRNPC interactors by RIP-seq, iCLIPseq and the ones displaying altered AS in hnRNPC KD cells KEGG mapper ^30^ was used (**Supplementary Data 6**). Comparative Toxicogenomics Database (CTD) ^31^ was used to detect the disease categories enriched using the list of transcripts common between the ones bound to hnRNPC in HF (RIP-seq analysis), the direct targets found by iCLIPseq and the transcripts displaying altered AS in hnRNPC KD cells (**Fig. 8**; **Supplementary Data 8**).

**15. Histology, immunohistochemistry and immunocytochemistry**

After collection, human heart tissues were dissected in cold DPBS, fixed for 4-8 h in 4% PFA at 4°C, cryoprotected in 30% sucrose overnight at 4°C, embedded in Tissue Freezing Medium (Leica Biosystems) and snap frozen using a dry ice/isopentane cooling bath. Frozen tissues were cut into 5 μm-thick sections using a cryostat (Leica).

Murine hearts were fixed for 4-8 h in 10% formalin neutral buffer (10%) (VWR BDH & Prolabo), cryoprotected by 30% sucrose immersion, embedded in Tissue Freezing Medium and cut into 3 μm-thick slices using a cryostat (Leica).

All cryosections were thawed at 4°C and rehydrated with DPBS for 5 min at room temperature (RT). Tissue permeabilization was performed with 0.5% Triton X-100 for 10 min at room temperature. After blocking non-specific sites, primary antibodies were incubated overnight at 4°C (**Supplementary Table 2**).

For immunocytochemical analysis of adherent cells, fixation was performed in 4% PFA for 15 min at room temperature, followed by 5 min permeabilization in 0.2% Triton X-100 (Sigma-Aldrich) and 1 h blocking in 2.5% BSA in DPBS solution at room temperature (RT). Cells were incubated with primary antibodies (**Supplementary Table 2**) for 2 h at RT or overnight at 4 °C. After three washes with DPBS, the samples were incubated with the respective Alexa fluorochrome-conjugated secondary antibodies (ThermoFisher Scientific) for 1 h at RT. DAPI (Sigma-Aldrich) was used to counterstain the nuclei. Specimens were then mounted using anti-fade Mowiol® 4-88 mounting media (Sigma-Aldrich) plus DABCO (Sigma-Aldrich). Masson’s Trichrome (Sigma-Aldrich) staining was performed according to the manufacturer’s instructions using celestine blue (Sigma- Aldrich) and bouin’s solution (VWR Chemicals).

**16. Image analysis**

A Zeiss LSM 780 confocal microscope with × 40 (1.3 NA) oil-immersion or x 20 air (0.8 NA) objective lenses were used for immunostaining read-out acquisition. Z-stack images were acquired with the optimal interval suggested by the software, followed by the application of maximum intensity algorithm. For the Masson’s Trichrome analysis, a slide scanner Zeiss Axio Scan Z1 microscope was used to visualize the whole tissue in bright-field mode and through a x10 objective. Super-resolution imaging was performed using a 3D-SIM Deltavision OMX with x 60 objective (1.42 NA) using 524 oil. Expression profiles were obtained by RGB profiler using ImageJ software.

**16.1 PLA quantification**

For each experiment and respective conditions, a minimum of six Z-stacks were acquired with a confocal microscope and a x40 oil objective. The same confocal acquisition settings were used within the same experiment. PLA dots quantification was performed using ImageJ software as follows: 10 regions of interest (ROIs) of the same size were defined for each Z-stack, a threshold of 50-150 (signal histogram) was set on the red channel (PLA signal) and all the slices of the Z-stack were processed as binary and watershield. The dots in the ROIs were quantified using the “3D Object counter”.

**16.2 Co-localization analysis**

Co-localization analysis was performed using Imaris 9.0 software. For each experiment a minimum of four Z-stacks *per* condition were acquired with a confocal microscope and a x40 oil objective. The same confocal acquisition settings were used across the experiment. A mask was applied to the channel corresponding to the sarcomeric structure (TNNT2) and automatic threshold was selected. Co-localization channel statistics were then automatically calculated by Imaris software. The threshold, the colocalization channel and the colocalization coefficient (Pearson’s r) were calculated by Imaris software.

**17. Individual-nucleotide resolution UV crosslinking and immunoprecipitation (iCLIP) of HNRNPC1/C2**

**17.1 Cell culture and crosslinking**

Normal Human Dermal Fibroblasts (NHDF) were grown in DMEM supplemented with FBS to 85-90% confluency. Three 150mm tissue culture dishes were used for each replicate of iCLIP. Cells were washed briefly with ice cold 1x PBS and exposed to 150 mJ/cm^2^ at 254 nm in a Stratalinker 2400 on ice. The cells were harvested by scraping, washed with 1x ice cold PBS. Pooled pellets from three dishes were flash frozen in liquid nitrogen and stored at -80°C until further use. Pellets were thawed and resuspended in 1ml of ice-cold lysis buffer (50 mM Tris-HCl, pH 7.4; 100 mM NaCl; 1% NP-40; 0.1% SDS; 40 U / ml of RNase inhibitor, 1x CompleteMini Protease Inhibitor Cocktail (Roche)), the suspension was passed through g20 syringe and incubated 10 min on ice. The insoluble fraction was removed by centrifugation at 20.000g for 20 min at 4°C and the supernatant was used for the iCLIP experiments.

**17.2 iCLIP**

100 µl of Protein G Dynabeads (Invitrogen) were coupled with 15 µg of anti HNRNPC1/C2 antibody (Santa Cruz) in a total volume of 100 µl of lysis buffer for 1 h at RT. The beads were washed two times with 1ml of low salt wash buffer (LSWB; 50mM Tris pH 7.4, 150mM NaCl, 0.02% NP-40, 1mM DTT, 1x CompleteMini Protease Inhibitor Cocktail), two times with 1ml of high salt wash buffer (HSWB 50mM Tris pH 7.4, 1M NaCl, 1mM EDTA, 1% NP-40, 0.1% SDS, 1x CompleteMini Protease Inhibitor Cocktail) and two times 1ml lysis buffer. Prior to protein IP, the beads were incubated for 1h at 4°C in lysis buffer containing 0.1 mg/ml heparin (Sigma-Aldrich) and washed as previously mentioned.

Antibody coupled beads were mixed with 1 ml of soluble cell lysate and incubated for 1hour on a rotator at 4°C. The unbound fraction was collected on a magnetic stand. The beads were consequently washed three times with 1ml of LSWB, HSWB and PNK buffer (100mM Tris pH 7.4, 50mM NaCl, 10mM EDTA), each time for 10 min at 4°C with rotation.

The beads were transferred into fresh tubes and resuspended in 500 µl of LSWB supplemented with 0.2 units of RNase I, 5 min at 37°C, with agitation at 1100 RPM. The reaction was stopped by incubation on ice and the beads were subjected to an additional round of high stringency washing. The bound fractions were treated with two units of alkaline phosphatase (Fast‐AP, Thermofisher) in 1X FastAP buffer in a reaction volume of 25 µl and incubated 20 min at 37°C. 1/10V of the beads were incubated with 8μl of radiolabelling mix (0.4 μl T4 PNK (NEB); 0.8 μl 32P-γ-ATP; 0.8 μl 10x PNK buffer (NEB)) for 20 mins at 37°C at 1100 RPM, for reference purpose and the rest of the beads were incubated with 20 μl of cold ATP mix (1μl T4 PNK, 1mM ATP, 20μl 10x PNK buffer). 20μM of L3App adapter ^32^ was ligated to the bound RNA in a final volume of 20μl ligation mix (1x RNA ligase buffer, 5% PEG400 (Sigma-Aldrich), 20 Units RNase inhibitor and 10 Units T4 RNA Ligase 2 (truncated) (NEB). The ligation reaction was incubated overnight at 16°C while rotating at 1100 RPM. The beads were washed with 1ml of HSWB and after each wash transferred to fresh non-sticky 1.5 ml eppendorf tubes to get rid of any non-ligated linkers.

The protein-RNA complexes were eluted by adding 20 μl of 1X NuPAGE loading buffer supplemented with 1x NuPAGE reducing agent and incubating 5 min at 90 °C. The eluted proteins and protein-RNA complexes were resolved by 4-12% gradient SDS-PAGE (NuPAGE, Invitrogen). The region above the migration position of the free protein till the position migrating at 100KDa was cut from the nitrocellulose membrane post transfer and subjected to proteinase K treatment (100mM Tris pH 7.4 50mM NaCl, 10mM EDTA, 100u proteinase K) for 20 min at 37°C. The reactions were stopped with 1V of Urea buffer (100mM Tris pH 7.4, 50mM NaCl, 10mM EDTA, 7M Urea) and incubated for additional 20 min. RNA was purified by adding equal volumes of Phenol chloroform isoamylalcohol and precipitated with 3V of 100 % ethanol overnight at -20°C.

**17.3 cDNA library preparation and sequencing**

The cDNA libraries were prepared as previously described ^33^ with slight modifications. Briefly, RNA pellets were washed 2x with ice cold 80% ethanol and resuspended in 8μl of RT annealing mixture (Rt1iClip, Rt2iClip and Rt3iClip-0.5μM each, 1mM dNTPs) (**Supplementary Table 2**). The samples were denatured three min at 70°C and cooled to 25°C for 10 min. 12 μl RT reaction mix was added (1x first strand buffer, 5mM DTT, 200u SSIII) and reverse transcription reaction was carried out in a thermal cycler (25°C for 20 min, 42°C for 30 mins, 50°C for 45 mins, 90°C for 3 mins).

The samples were snap-chilled on ice and passed through the QIAquick columns (QIAGEN) following the manufacturer’s protocol. DNA was eluted in 20μl ddH2O. DNA circularization was carried in a final reaction volume of 40 μl of the circligaseII reaction mix (1X CircLigase Buffer II (Lucigen), 500mM MnCl2, 60U CircLigase II (Lucigen)) at 60°C for 1hour. The circularised samples were supplemented with oligo annealing mix (11μl ddH2O, 4.5μl 10x Fast digest buffer (Thermofisher), 1.5μl Cut-Oligo **(Supplementary Table 2)**. The samples were incubated 4 min at 95°C and the temperature was subsequently reduced by 1°C every 30 seconds till final temperature of 25°C. Post hybridization the reaction mix was supplemented with 3μl of BamHI and incubated for 1hour at 37°C. Linearized cDNA was further purified using the QIAquick columns and eluted in final volume of 28 μl. cDNA PCR amplification was carried out with P3/P5 primer mix (1mM each) and 1x Phusion Flash High-Fidelity PCR Master Mix (Thermofisher). The PCR products were resolved by a 4% metaphor agarose gel electrophoresis and the fragments of 180-300 bp were excised and purified by QIAquick gel purification following the manufacturer’s protocol. The resulting DNA libraries were quality checked on a bioanalyzer. The libraries were pooled in equimolar concentration and analysed by 75bp single end high-throughput sequencing at Illumina HiSeq2500. A schematic representation of the protocol can be found in **Supplementary Fig. 7f**.

**17.4 iCLIP-seq analysis**

The iCLIP libraries contained 5-nt random Unique molecular identifier sequence and 4-nt sample barcode, allowing for demultiplexing of samples as well as for PCR duplicates removal. We gathered about 250 million of reads across three replicates. Quality check of raw fastq reads was carried out by FastQC ^12^. The adapters and quality trimming of raw fastq reads was performed using Trimmomatic v. 0.36 ^13^ with settings CROP:250 LEADING:3 TRAILING:3 SLIDINGWINDOW:4:5 MINLEN:20 and adapter sequences AGATCGGAAGAGCGG, ATCTCGTATGCCGTC, GGGGGGGGGGGGGGG were removed using ILLUMINACLIP setting. Trimmed iCLIP-Seq reads were mapped against the human genome (hg38) and Ensembl GRCh38 v.91 annotation using STAR v. 2.5.3a ^14^ as splice-aware short read aligner and default parameters except --outFilterMismatchNoverLmax 0.1 and --twopassMode Basic. Quality control after alignment was performed using several tools namely RSeQC ^15^, Picard toolkit (GitHub Repository. http://broadinstitute.github.io/picard/) and Qualimap ^16^. Unique molecular identifiers were extracted and used for PCR duplicates removal using UMI-tools v. 0.5.5 ^34^. Counts and ratio of raw reads, trimmed reads, uniquelly/multimapped reads and deduplicated reads can be found in **Supplementary Data 7**.

**17.5 Identification of genomic crosslink sites**

To obtain genomic crosslink sites, only uniquely mapped and deduplicated reads were used, to avoid an uncertainty of ambiguously mapped reads. Uniquelly mapped and deduplicated reads were converted into bedgraphs genomic coverage tracks using deeptools bamCoverage v. 3.3.0 ^35^ with settings -of bedgraph --binSize 1 --normalizeUsing None. Bedgraphs were further normalized on number of mapped reads per million (CPM). All of the covered positions were filtered in a way to keep only these regions, where there is more than one uniquelly mapped deduplicated read, to get rid of bias from possible random false positive mapping. All crosslink sites that were closer than 5 bases to each other were merged together to create more robust binding sites. All binding sites were annotated employing Ensembl GRCh38 v.91 annotation file. As a binding site were considered all crosslink regions which appeared in at least 2 out of 3 replicates. Altogether we observed 13,567 crosslink sites, affecting 5,536 different genes and 2,988 intergenic regions.

For the visualization of binding sites in a close proximity of observed skipped exons all binding sites were averaged across all replicates and visualize in Integrative genomics viewer ^36^.

To assess the ratio between binding sites in coding and non-coding regions, counts, produced using featureCounts tool ^18^ with settings -s 1 -T 10 -F GTF -Q 0 -d 1 -D 25000 and calculated over feature "gene" including all parts of gene region specified in annotation file - intronic parts of gene are included as well, were adopted. All genomic counts were again annotated using Ensembl GRCh38 v.91. As a non-coding features were considered all records with gene_biotype falling into this list: "3prime_overlapping_ncRNA", "antisense_RNA","bidirectional_promoter_lncRNA", "lincRNA", "non_coding", "processed_transcript", "sense_intronic", "sense_overlapping", "miRNA","macro_lncRNA", "misc_RNA", "Mt_rRNA", "Mt_tRNA", "rRNA","scaRNA", "scRNA" , "snoRNA", "snRNA", "sRNA" , "tRNA", "tRNA_pseudogene","vaultRNA".

All intergenic binding sites were considered as non-coding as well. The rest of genomic counts were treated as coding regions. Based on the same counts and using gene_biotype list containing: lincRNA, Non-antisense_RNA, snRNA, Non-misc_RNA, bidirectional_promoter_lncRNA, snoRNA, rRNA, macro_lncRNA, 3prime_overlapping_ncRNA and miRNA, the counts and ratio within these annotated non-coding genomic features were derived (**Supplementary Fig. 7i; Supplementary Data 7**).

Read counts of 4 specific genomic features (CDS - exons without UTR regions, introns, 3'UTR and 5'UTR) were gathered using featureCounts and were again based on modified Ensembl GRCh38 v.94 annotation GTF file described in RIP-Seq methods part. All iCLIP sample counts were normalized on sequencing library depth (CPM) (**Fig. 7e**).

We used Alu elements annotation based on RepeatMasker predictions (Smit, AFA, Hubley, R & Green, P. RepeatMasker Open-4.0. 2013-2015; http://www.repeatmasker.org) to obtain crosslink sites on the total of 1,131,306 Alu elements. Only deduplicated reads without filtering of multimapped reads were converted into bed files using deeptools bamCoverage. These bed files were further intersected with bed file of Alu elements annotated regions utilizing bedtools intersect v. 2.23.0 ^37^. We considered all annotated Alu elements with any mapped read as covered Alu and the rest as Not-covered Alu (**Supplementary Fig. 7h**).

All Pie plots were produced using R package plotrix ^20^ and all barplots plotted using R package ggplot2 ^19^.

17.6 iCLIP-PCR validation assay

NHDFs were crosslinked at 150mJ/cm^2^ UV and lysed using 3X pellet volume of lysis buffer supplemented with RNAse inhibitor and protease inhibitor. The pull down was carried out on three biological replicates as described for the iCLIP assay (see 17.1), excluding the RNAse treatment step. Post IP, the beads were subjected to DNAse treatment to remove genomic DNA contamination, followed by 3' linker ligation and proteinase K treatment to release the bound RNA. Immunoprecipitation with FLAG antibody was used as negative control. Total RNA isolated from lysates were used as positive controls.

|  |  | **Study groups** | |
| --- | --- | --- | --- |
|  |  | **Control** | **Heart Failure** |
|  |  | N = 3 | N = 17 |
|  | **Actual surgery** | Autopsy (100%) | LVAD (53%) |
|  |  |  | Heart transplant (47%) |
|  | **Age (years)** | 52±3 | 53 ± 15 |
|  | **Gender** | M (100%) | M (88%) F (12%) |
| **Physiology** | **BMI (kg/m^2^)** | 25 ± 2 | 27 ± 5 |
|  | **Heart rate (bpm)** | 80 ± 20 | 80 ± 16 |
|  | **Ejection Fraction LV (%)** | 60 ± 5 | 19 ± 12 |
|  | **NYHA I** | n/a | 0 |
| **Heart Failure** | **NYHA II** | n/a | 6 |
| **Functional Classification** | **NYHA III** | n/a | 29 |
| (%) | **NYHA IV** | n/a | 65 |
|  | **Arrhythmia** | n/a | 71 |
|  | **Congestive Heart Failure** | n/a | 65 |
| **Cardiac pathologies** | **Ischemic Heart Disease** | n/a | 53 |
| (%) | **Dilated Cardiomyopathy** | n/a | 65 |
|  | **Danon disease** | n/a | 6 |

**Supplementary Table 1. Patients enrolled in the study.** Data are presented as the ratio in percentage or mean ± SD. Abbreviations: F, female; LV, left ventricle; LVAD, left ventricular assist device; M, male; NYHA, New York Heart Association Functional Classification; n/r, not reported; n/a, not applicable.

| REAGENT or RESOURCE | SOURCE | IDENTIFIER |
| --- | --- | --- |
| Antibodies | | |
| Mouse monoclonal anti-hnRNPC (4F4) | Santa Cruz Biotechnology | Cat# sc-32308, RRID:AB_627731 |
| Rabbit monoclonal anti-hnRNPC (EP3034Y) | Abcam | Cat# ab75822, RRID:AB_1310320 |
| Rabbit monoclonal anti-hnRNPC (EPNCIR152) | Abcam | Cat# ab133607, RRID:AB_2860560 |
| Rabbit polyclonal anti-TNNT2 | Sigma Aldrich | Cat# HPA017888, RRID:AB_1858355 |
| Mouse monoclonal anti-alpha-actinin (EA-53) | Sigma-Aldrich | Cat# A7811, RRID:AB_476766 |
| Rabbit polyclonal anti-FHL2 | Proteintech | Cat# 21619-1-AP, RRID:AB_10860263 |
| Rabbit polyclonal anti-PDLIM5 | Sigma-Aldrich | **Cat# HPA016740, RRID:AB_1855153** |
| Mouse monoclonal anti-PDLIM5 | Sigma-Aldrich | **Cat#WH0010611M1** RRID:AB_1842923 |
| Mouse monoclonal anti-RPS6 (C8) | Santa Cruz Biotechnology | Cat# sc-74459, RRID:AB_1129205 |
| Mouse monoclonal anti-PMY (12D10) | Millipore | Cat# MABE343, RRID:AB_2566826 |
| Mouse monoclonal anti-Lamin A/C (4C11) | Sigma -Aldrich | Cat# SAB4200236, RRID:AB_10743057 |
| Mouse monoclonal anti-YAP (SPM227) | Santa Cruz Biotechnology | Cat# sc-101199, RRID:AB_1131430 |
| Mouse monoclonal anti-vinculin (63.7) | Abcam | Cat# ab18058, RRID:AB_444215 |
| Mouse monoclonal Anti-β-Actin | Sigma-Aldrich | Cat# A2228,  RRID:AB_476697 |
| Normal mouse IgG | Santa Cruz Biotechnology | Cat# sc-2025, RRID:AB_737182 |
| Normal rabbit IgG | Santa Cruz Biotechnology | Cat# sc-2027, RRID:AB_737197 |
| Mouse monoclonal anti-Flag (M2) | Sigma-Aldrich | Cat# F3165,  RRID:AB_259529 |
| Rabbit polyclonal anti-MYH7 | Cloud-Clone Corp. | Cat# PAD418Hu01 |
| F-actin-Alexa Fluor 555 | ThermoFisher Scientific | Cat# A34055 |
| F-actin-Alexa Fluor 647 | ThermoFisher Scientific | Cat# A22287, RRID:AB_2620155 |
| GAPDH-HRP (GA1R) | Invitrogen | Cat# MA5-15738-HRP, RRID:AB_2537659 |
| Chicken anti-mouse IgG-HRP | Santa Cruz Biotechnology | Cat# sc-2954, RRID:AB_639239 |
| Chicken anti-rabbit IgG-HRP | Santa Cruz Biotechnology | Cat# sc-2963, RRID:AB_639249 |
| Alexa Fluor 488 donkey anti-mouse IgG | ThermoFisher Scientific | Cat# A-21202, RRID:AB_141607 |
| Alexa Fluor Plus 555 goat anti-rabbit IgG | ThermoFisher Scientific | Cat# A32732, RRID:AB_2633281 |
| Alexa Fluor 555 donkey anti-rabbit IgG | ThermoFisher Scientific | Cat# A-31572, RRID:AB_162543 |
| Alexa Fluor 488 goat anti-mouse IgG | ThermoFisher Scientific | Cat# A-11001, RRID:AB_2534069 |
| TNNT-FITC | Miltenyi Biotec | Cat# 130-119-575, RRID:AB_2751735 |
| Biological Samples | | |
| Patient-derived heart tissues | Centre of Cardiovascular and Transplantation Surgery, Brno, Czech Republic | N.A |
| Chemicals, Peptides, and Recombinant Proteins | | |
| Lipofectamine RNAiMAX | Invitrogen | Cat# 13778075 |
| Dynabeads | Invitrogen TM | Cat# 10007D |
| Zirconia beads | Benchmark Scientific | Cat# D1032-30 |
| Precision Plus Protein™ Dual Color Standards | Biorad | Cat# 1610374 |
| Trizol reagent | Invitrogen | Cat# 15596026 |
| DTT | Bio-Rad | Cat# 1610611 |
| DNase I | Roche | **Cat# 04716728001** |
| Salt I | Millipore | **Cat#** CS203173 |
| Salt II | Millipore | **Cat#** CS203185 |
| Precipitate Enhancer | Millipore | **Cat#** CS203208 |
| Cycloheximide | Sigma Aldrich | **Cat#** C7698 |
| Proteinase K | Invitrogen TM | **Cat#** AM2548 |
| Puromycin | Santa Cruz Biotechnology | **Cat#** sc-108071 |
| DAPI | Sigma Aldrich (Roche) | **Cat#** 10236276001 |
| Mowiol® 4-88 mounting media | Sigma Aldrich | **Cat# 81381** |
| DABCO 33-LV | Sigma Aldrich | **Cat#** 290734 |
| Bouin’s solution | VWR Chemicals | **Cat#** 7000.1000 |
| Celestin blue | Sigma Aldrich | **Cat#** 206342 |
| Heparin (5000IU / ML) | Zentiva | **Cat#** 8594739026131 |
| Fibrinogen (Tisseel_ Sealant Kit) | Baxter Bioscience | **Cat#** 1504514 |
| Thrombin (Tisseel_ Sealant Kit) | Baxter Bioscience | **Cat#** 1504514 |
| Aprotinin (Tisseel_ Sealant Kit) | Baxter Bioscience | **Cat#** 1504514 |
| RIPA lysis buffer | Millipore | **Cat#** 20-188 |
| Rnase-OUT | Invitrogen | **Cat#** 0777019 |
| Tergitol | Sigma-Aldrich | **Cat#** NP40S |
| Latrunculin | Sigma-Aldrich | **Cat#** 76343-93-6 |
| Y27632 | Selleck Chemicals | **Cat#** S1049 |
| CHIR99021 | Sigma-Aldrich | **Cat#** SML1046 |
| IWP2 | Selleck Chemicals | **Cat#** S7085 |
| B27 supplement minus | Thermo FisherScientific | **Cat#** A1895601 |
| B27 supplement | Thermo FisherScientific | **Cat#** 17504044 |
| Lithium dodecyl sulfate | Sigma-Aldrich | Cat#L9781 |
| IGEPAL CA-630 | Sigma-Aldrich | Cat#I3021 |
| CompleteMini Protease Inhibitor Cocktail | Roche | Cat#11836170001 |
| RNAseI | Thermo Fisher | Cat# EN0601 |
| TurboDNase | Thermo Fisher | Cat# AM2239 |
| T4 RNA Ligase 2 (truncated) | New England Biolabs | Cat#M0242S |
| CircLigase II | Lucigen | Cat#CL9021K |
| PEG400 | Sigma-Aldrich | Cat#8074850050 |
| Linear Acrylamide | Thermo Fisher | Cat#AM9520 |
| Protein G | Thermo Fisher | Cat#10003D |
| Critical Commercial Assays | | |
| High Pure RNA Isolation Kit | Roche | **Cat#** 11 828 665 001 |
| SYBR Green I Master Kit | Roche | **Cat#** 04707516001 |
| Phusion Flash High-Fidelity PCR Master Mix | Thermo Fisher | Cat#F-548L |
| Transcription First Strand cDNA Synthesis Kit | Roche | **Cat#** 04896866001 |
| Pierce BCA Protein Assay Kit | ThermoFisher | **Cat#** 23225 |
| Duo-link in situ PLA Kit | Sigma-Aldrich | **Cat#** DUO92101 |
| NE-PERtm Nuclear and Cytoplasmic extraction kit | ThermoFisher Scientific | **Cat#** 78833 |
| QIAseq Fastselect HMR Kit | Qiagen | **Cat#** 334376 |
| M.O.M basic kit | Baria | **Cat#** BMK-2202 |
| MinElute Gel Extraction kit | Qiagen | Cat#28604 |
| QIAquick PCR purification kit | Qiagen | Cat#28104 |
| SuperScript III Reverse Transcriptase Kit | Sigma-Aldrich | Cat#18080044 |
| Dynabeads™ Protein G Immunoprecipitation Kit | Invitrogen TM | **Cat#** 10007D |
| Deposited Data | | |
| TMT-Mass spectrometry data (hnRNPC interactome) | This study | Deposited to PRIDE  PXD020663 |
| RIP-sequencing data (hnRNPC-RNAs targets) | This study | Deposited to GEO  GSE155472 |
| RNA-sequencing data (AS in hnRNPC KD cells) | This study | Deposited to GEO  GSE155473 |
| iCLIP-seq data (hnRNPC in NHDF cells) | This study | Deposited to GEO GSE169068 |
| Mouse 48h post-MI dataset | Harpster et al., 2006 | GSE4648 |
| Mouse heart failure dataset | Rowell et al., 2014 | GSE54681 |
| Human heart failure dataset | Barth et al., 2006 | GSE3586 |
| Human heart failure dataset (AS analysis) | Pepin et al., 2019^38^ | GSE108157 |
| Human heart failure dataset (AS analysis) | MAGNet; www.med.upenn.edu/magnet | GSE141910 |
| Experimental Models: Cell Lines | | |
| iPSCs | WiCell (Madison, WI, USA | DF 19-9-7T |
| NHDFs | ATCC | ATCC® PCS-201-012™ |
| Experimental Models: Organisms/Strains | | |
| C57BL/6 | IBMC-INEB | Nascimento et al.,2011^2^ |
| Oligonucleotides | | |
| hnRNPC Forward 5’- TCCCCTTCTTGTTTTCGGCT -3’ | Generi Biotech (custom) | N.A |
| hnRNPC Reverse 5’ – CTGAGTAGAGGGGACGGAGA -3’ | Generi Biotech  (custom) | N.A |
| GAPDH Forward 5’-AAGTATGACAACAGCCTCAA-3’ | Invitrogen (custom) | N.A |
| GAPDH Reverse 5’- TCCTTCCACGATACCAAAGT-3’ | Invitrogen (custom) | N.A |
| YAP-1α Forward 5’-GATGAACTCGGCTTCAGCCATGAA-3’ | Generi Biotech | Vrbsky et al., 2021^39^ |
| YAP-1α Reverse 5’-GCAGGGCTAACTCCTGCCGAAGCA-3’ | Generi Biotech | modified from Karystinou 2015^40^ |
| YAP-2α Forward 5’-CCTCTTCCTGATGGATGGGA-3’ | Generi Biotech | Vrbsky et al., 2021^39^ |
| YAP-2α Reverse 5’-GCAGGGCTAACTCCTGCCGAAGCA-3’ | Generi Biotech | Vrbsky et al., 2021; Karystinou 2015^39,40^ |
| YAP-1β Forward 5’-GATGAACTCGGCTTCAGCCATGAA-3’ | Generi Biotech | Vrbsky et al., 2021^39^ |
| YAP-1β Reverse 5’-GCAGGGCTAACTCCTGTGGCCTCA-3’ | Generi Biotech | Vrbsky et al., 2021^39^ |
| YAP-2β Forward 5’-CCTCTTCCTGATGGATGGGA-3’ | Generi Biotech | Vrbsky et al., 2021^39^ |
| YAP-2β Reverse 5’-GCAGGGCTAACTCCTGTGGCCTCA-3’ | Generi Biotech | Vrbsky et al., 2021^39^ |
| YAP-1γ Forward 5’-GATGAACTCGGCTTCAGCCATGAA-3’ | Generi Biotech | Vrbsky et al., 2021^39^ |
| YAP-1γ Reverse 5’-TATTCCGCATTGCCTGCCGAAGCA-3’ | Generi Biotech | Vrbsky et al., 2021; Karystinou 2015^39,40^ |
| YAP-2γ Forward 5’-CCTCTTCCTGATGGATGGGA-3’ | Generi Biotech | Vrbsky et al., 2021^39^ |
| YAP-2γ Reverse 5’-TATTCCGCATTGCCTGCCGAAGCA-3’ | Generi Biotech | Vrbsky et al., 2021; Karystinou 2015^39,40^ |
| YAP-1δ Forward 5’-GATGAACTCGGCTTCAGCCATGAA-3’ | Generi Biotech | Vrbsky et al., 2021^39^ |
| YAP-1δ Reverse 5’-ATTGCCTGTGGCCTCACCT-3’ | Generi Biotech | Vrbsky et al., 2021; Karystinou 2015^39,40^ |
| YAP-2δ Forward 5’-CCTCTTCCTGATGGATGGGA-3’ | Generi Biotech | Vrbsky et al., 2021^39^ |
| YAP-2δ Reverse 5’-ATTGCCTGTGGCCTCACCT-3’ | Generi Biotech | Vrbsky et al., 2021; Karystinou 2015^39,40^ |
| YAP-i4 Forward 5’-AGCCCACTCGGGATGTAACTTGA-3’ | Generi Biotech | Vrbsky et al., 2021; Karystinou 2015^39,40^ |
| YAP-i4 Reverse 5’-CTGGTGGGGGCTGTGACGTT-3’ | Generi Biotech | Vrbsky et al., 2021; Karystinou 2015^39,40^ |
| YAP1 Forward 5’- CCTGAACAGTGTGGATGAGA-3’ | Generi Biotech (custom) | N.A |
| YAP1 Reverse 5’- GCTTCAAGGTAGTCTGGGAA-3’ | Generi Biotech (custom) | N.A |
| CD44 Forward 5’- GGAGCAGCACTTCAGGAGGTTAC-3’ | Generi Biotech (custom) | Wang et al., 2017^41^ |
| CD44 Reverse 5’- GGAATGTGTCTTGGTCTCTGGTAGC-3’ | Generi Biotech (custom) | Wang et al., 2017^41^ |
| EX2-3 Yap1 (F)- ACTTCTTAAATCACATCGAT | Generi Biotech (custom) | N.A |
| EX5 Yap1 (R): TGCCGAAGCAGTTCTTGCTG | Generi Biotech (custom) | N.A |
| CD44 ex v6 (F): GCAACTCCTAGTAGTACAACG | Generi Biotech (custom) | N.A |
| CD44 ex v8 (R): GCGTTGTCATTGAAAGAGGTC | Generi Biotech (custom) | N.A |
| MYLK ex11 Forward 5’- GAGCCAAGATGTTGTGAGCA-3’ | Generi Biotech (custom) | Miao et al., 2010^42^ |
| MYLK ex11 Reverse 5’- ATTCAGCAGCCAAGTGATCC-3’ | Generi Biotech (custom) | Miao et al., 2010^42^ |
| COL6A3 ex3 Forward 5’- AGCAGCAAGCAGATGTCAAA-3’ | Generi Biotech (custom) | Arafat et al., 2011^43^ |
| COL6A3 ex3 Reverse 5’- TTTCTCCCACAGCTAAGGATTT-3’ | Generi Biotech (custom) | Arafat et al., 2011^43^ |
| HNRNPC siRNA | Invitrogen | HSS179304 |
| HNRNPC siRNA | Invitrogen | HSS179305 |
| stealth siRNA negative control low GC | Invitrogen | 12935200 |
| Software and Algorithms | | |
| ImageJ | Schneider et al., 2012^44^ | <https://imagej.nih.gov/ij/> ; RRID:SCR_003070 |
| Imaris 9.0 | Oxford Instruments | RRID:SCR_007370 |
| Cytoscape 3.7.2 | Shannon, 2003^25^ | RRID:SCR_003032 |
| Prism v 8.0.1 | GraphPad | <https://www.graphpad.com/> ; RRID:SCR_002798 |
| rMATS v3.1.0 | Shen et al., 2014^21^ | rnaseq-mats.sourceforge.  net/. |
| POSTAR2 | Zhu et al., 2019^29^ | http://lulab.life.tsinghua.edu.cn/postar/ |
| ClueGO v2.5.6 | Bindea et al., 2009^26^ | RRID:SCR_005748 |
| CluePedia v1.5.6 | Bindea et al., 2013^27^ | RRID:SCR_015784 |
| EnrichR | Chen et al., 2013; Kuleshov et al., 2016^23,24^ | <http://amp.pharm.mssm.edu/Enrichr> ; RRID:SCR_001575 |
| Integrative Genomics Viewer | Broad Institute | http://software.broadinstitute.org/software/igv/; |
| KEGG mapper | Kanehisa and Sato, 2020^30^ | RRID:SCR_018145 |
| WikiPathways | Slenter et al., 2018^28^ | RRID:SCR_002134 |
| Comparative Toxicogenomics Database (CTD) | Davis et al., 2018^31^ | http://ctdbase.org/ ; RRID:SCR_006530 |
| FastQC | Andrews, 2010^12^ | RRID:SCR_014583 |
| Trimmomatic v. 0.36 | Bolger et al.,2014^13^ | RRID:SCR_011848 |
| STAR v. 2.5.3a | Dobin et al.,2013^14^ | RRID:SCR_015899 |
| RSeQC v. 2.6.2 | Wang et al.,2012^15^ | RRID:SCR_005275 |
| Mascot version 2.5.1 | Matrix Science | RRID:SCR_014322 |
| Picard toolkit v. 2.18.27 | GitHub Repository | <http://broadinstitute.github.io/picard/>; RRID:SCR_006525 |
| Qualimap v.2.2.2 | Okonechnikov et al.,2016^16^ | RRID:SCR_001209 |
| FastQ Screen v. 0.13.0 | Wingett and Andrews,2018^17^ | RRID:SCR_000141 |
| bedtools subtract v2.29.0 | Quinlan et al.,2010^37^ | RRID:SCR_006646 |
| featureCounts v. 1.5.2 | Liao et al.,2014^18^ | RRID:SCR_012919 |
| Ggplot2 v3.3.1 | Wickham, 2011^19^ | RRID:SCR_014601 |
| plotrix v3.7-8 | Lemon, 2006^20^ | https://www.rdocumentation.org/packages/plotrix/versions/3.7-8 |
| Rmats2sashimiplot v2.0.2 | Xinglab, 2015a^45^ | https://github.com/Xinglab/rmats2sashimiplot |
| clusterProfiler v3.12.0 | Yu et al., 2012^22^ | RRID:SCR_016884 |
| Proteomaps | Liebermeister et al., 2014; Otto et al., 2010^4,5^ | https://bionic-vis.biologie.uni-greifswald.de/ |
| RepeatMasker | Smit, AFA, Hubley,  R & Green, 2013-2015 | http://www.repeatmasker.org |
| UMI-tools v. 0.5.5 | Smith et al., 2017^34^ | RRID:SCR_017048 |
| bamCoverage v. 3.3.0 | Ramírez et al., 2016^35^ | NA |
| Adobe Illustrator | Adobe | <https://www.adobe.com/> ; RRID:SCR_010279 |
| Others | | |
| Micropattern | CYTOO, Grenoble, France | ref: 10–950–10–18 |
| Micropattern | CYTOO, Grenoble, France | ref: 10–950–00–18 |
| Mini-PROTEAN® TGX™ Precast Protein Gels | BioRad | Cat# 4561034 |
| µ-Dish 35 mm, high ESS | ibidi GmbH, Munich, Germany | Cat# 81291 |

**Supplementary Table 2. Key Resource Table.** The table includes all the reagents, deposited data, assays and software used in this paper.

45.	Xinglab. Actin-associated hnRNP proteins as transacting factors in the control of mRNA transport and localization. *Available online at: https://github.com/Xinglab/rmats2sashimiplot* doi:10.4161/rna.6.2.8195.
